# Precision Targeting of Troponin I Phosphorylation Prevents Diastolic Dysfunction while Preserving Systolic Performance

**DOI:** 10.1101/2025.11.18.689162

**Authors:** Ying-Chi Chao, Debra J. McAndrew, David Revuelta, Rahul Yadav, Wan-Hua Hong, Jianshu Hu, Milda Folkmanaite, Edoardo Valli, Mariana Gee Olmedilla, Andreas Koschinski, Nicoletta C. Surdo, Jakub Tomek, Diederik W. D. Kuster, Helen Edwards, Connor Blair, Evaldas Girdauskas, David Henderson, Josh Hurst, Adam Linekar, Yiangos Psaras, Julia Gorelik, Nicola Smart, Christopher Toepfer, Helen Maddock, Jonathan M. Elkins, Jolanda van der Velden, George Baillie, Craig A. Lygate, Cristina E. Molina, Manuela Zaccolo

**Affiliations:** Department of Physiology Anatomy and Genetics, University of Oxford, UK; Radcliffe Department of Medicine, University of Oxford, UK; Institute of Experimental Cardiovascular Research, University Medical Center Hamburg-Eppendorf, Germany; Department of Physiology, Amsterdam Cardiovascular Sciences, Amsterdam University Medical Centre (UMC), Amsterdam, the Netherlands; College of Veterinary, Medical and Life Sciences, University of Glasgow, UK; Department of Cardiovascular Surgery, University Heart Center Hamburg, Germany; Coventry University Group, Coventry University, UK; Imperial College, London, UK; InoCardia Ltd, Coventry, UK; Centre for Medicines Discovery, University of Oxford, UK

## Abstract

**Background:** Defective cardiac relaxation (diastolic dysfunction) is prevalent in heart failure, particularly in the context of diabetes, obesity, hypertension, and ageing, and is associated with increased mortality. Yet there is no available treatment that directly targets this abnormal relaxation. Phosphorylation of troponin I by cAMP-dependent protein kinase A (PKA) promotes relaxation, but global PKA activation has widespread, undesirable effects. Here we investigated whether selectively enhancing troponin I phosphorylation could improve relaxation without engaging the broader PKA signalling network.

**Methods:** To resolve cAMP signalling at subcellular scales, we combined real-time cAMP measurements with FRET-based genetically encoded reporters targeted to defined nanodomains, alongside biochemical and genetic approaches. Protein–protein interaction surfaces were mapped using peptide-array technology, which guided the design of a disruptor peptide to displace specific interactions. The peptide’s effects on cardiomyocyte function were assessed *in vitro* using rodent and human cardiomyocytes with biochemical assays, real-time imaging, and work-loop analysis. *In vivo* effects were evaluated through echocardiography and haemodynamic measurements.

**Results:** We identified a previously unrecognized regulatory mechanism in which the cAMP-hydrolyzing enzyme PDE4D9 interacts with troponin I and restricts local cAMP levels within a nanometre-scale domain, thereby selectively controlling troponin I phosphorylation. We found that PDE4D9–troponin I association is markedly increased in cardiac disease in both rodents and humans. A peptide that displaces PDE4D9 from troponin I in a mouse model of heart failure selectively enhances troponin I phosphorylation, accelerates cardiomyocyte relaxation, and prevents diastolic dysfunction, without compromising systolic contractility.

**Conclusions:** These findings reveal a highly specific regulatory nanodomain governing troponin I phosphorylation and points to a first-in-class, mechanism-based therapeutic strategy with the potential to address a major unmet contributor to heart failure.

**Clinical Perspective:** *What Is New?:*

- We identify a previously unrecognized, nanometre-scale regulatory mechanism in which PDE4D9 binds troponin I to locally suppress cAMP, limit troponin I phosphorylation and attenuate relaxation.
- We demonstrate that this mechanism becomes maladaptive in disease: PDE4D9–troponin I interaction is markedly upregulated in failing hearts in both rodents and humans, which can mechanistically explain the impaired cardiac relaxation.
- We show that this interaction is druggable: a rationally designed peptide can selectively displace PDE4D9, restore localized cAMP–PKA signalling, and normalize myocardial relaxation without global cAMP perturbation.
- Displacement of PDE4D9 from TPNI improves relaxation with no impact on the calcium transient amplitude and without adversely affecting systolic output and cardiac reserve.

*What Are the Clinical Implications?:*

- Diastolic dysfunction, a major cause of heart failure with no targeted treatments, may be amenable to therapies that modulate cAMP signalling with subcellular precision rather than global pathway modulation.
- Selective disruption of PDE4D9–troponin I binding offers a mechanism-based approach with potential to restore diastolic function while minimizing off-target effects typical of GPCR-directed therapies.
- By enhancing relaxation while preserving systolic function and cardiac reserve, this approach may overcome limitations of myofilament-targeted therapies currently under evaluation for diastolic dysfunction that exhibit negative inotropic effects.
- More broadly, this work establishes a platform for targeting localized signalling domains, a concept that may extend beyond cardiovascular disease to other pathological conditions in which dysregulated cAMP contributes to dysfunction.

## Introduction

Cardiac contraction (systole) and relaxation (diastole) are driven by contractile proteins arranged in sarcomeres consisting of interdigitated thick myosin and thin actin filaments (Fig. 1b). The sliding of thick and thin filaments past each other causes the sarcomeres to shorten during contraction and re-lengthen during relaxation. This process is regulated by the binding and unbinding of Calcium ions to and from troponin C (TPNC) which, together with troponin T (TPNT) and troponin I (TPNI), forms the regulatory troponin complex associated with the thin filament. During sympathetic stimulation, activation of the β-adrenergic receptor (β-AR)/cAMP/ PKA pathway leads to phosphorylation of multiple targets, including proteins that enhance contraction and relaxation, allowing the heart to adapt to higher workload demands. PKA phosphorylation of phospholamban (PLB) at the sarcoplasmic reticulum (SR) enhances the activity of the sarco-endoplasmic reticulum Calcium ATPase (SERCA), promoting Calcium reuptake into the SR and ensuring a greater Calcium reserve is available for release during the next contraction. PKA phosphorylation of the thick filament protein myosin-binding protein C (MyBPC) enhances contraction by extending the force-generating state of the sarcomere. Phosphorylation of TPNI facilitates relaxation by lowering the affinity of TPNC for Calcium, ensuring timely deactivation of myofilaments to match faster contraction rate^1^. In pathological conditions, such as heart failure (HF), chronic β-AR overstimulation disrupts cAMP signaling, impairing contractility and relaxation^2^. β-blockers mitigate these effects and improve cardiac efficiency by reducing oxygen demand and preventing further myocardial damage^3^. However, β-blockers are ineffective when the primary issue is not impaired contraction but inadequate cardiac relaxation^4^.

**Figure 1.**
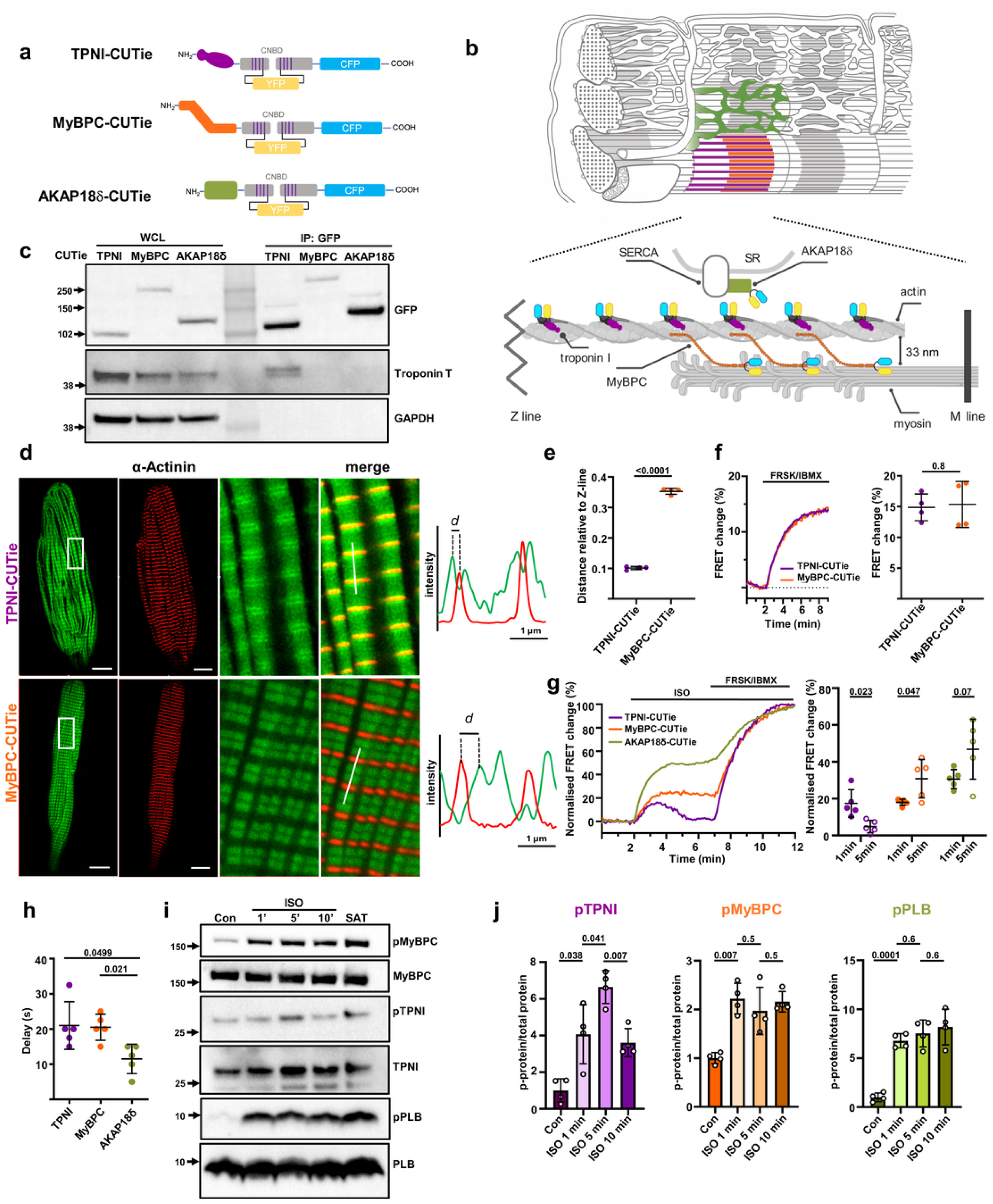
Unique cAMP regulation at TPNI. (a) Schematic representation of the targeted FRET reporters TPNI, MyBPC-, and AKAP18δ-CUTie and (b) their subcellular localisation in cardiac myocytes. CNBD, cAMP-binding domain. (c) Western blot of GFP immunoprecipitation from lysates of NRVM expressing TPNI-, MyBPC-, or AKAP18δ-CUTie and probed with GFP or troponin T antibodies. Representative of three independent experiments. (d) Representative confocal images of ARVM expressing TPNI-CUTie or MyBPC-CUTie showing sensor localisation relative to α-actinin. Enlarged views and merged images of sensor localisation (green) and α-actinin (red) are shown. Fluorescence intensity profiles measured across a single sarcomere along the indicated line are shown on the right. Dotted lines indicate peak intensity of α-actinin and sensor signals used to determine the distance (d) between sensor localisation and the Z-line, quantified in (e). Statistical significance was assessed using Welch’s t-tests. (f) Representative kinetics and average maximal FRET responses of TPNI- or MyBPC-CUTie expressed in HEK293 cells following saturating stimulation (25 μM forskolin and 100 μM IBMX). Statistical significance was assessed using Welch’s t-tests. (g) Representative kinetics (left) and quantification of FRET responses at 1 and 5 min following stimulation with 5 nM ISO (right) in ARVM expressing the indicated targeted CUTie reporters. Forskolin (25 μM) and IBMX (100 μM) were applied at the end of each experiment to achieve sensor saturation. Statistical significance was assessed using paired t-tests. (h) Average time required to achieve 5% of the maximal FRET response following stimulation with 5 nM ISO. Statistical significance was assessed using Welch’s t-tests with Benjamini–Hochberg correction. (i) Representative western blot of ARVM lysates showing total protein abundance and phosphorylation levels of MyBPC, TPNI, and PLB at 1, 5, and 10 min following stimulation with 5 nM ISO. (j) Quantification of experiments shown in (i). Statistical significance was assessed using Welch’s t-tests with Benjamini–Hochberg correction. Each data point represents an independent biological replicate. Data are presented as mean ± SD. n ≥ 4 biological replicates per group.

The cAMP/PKA signalling pathway operates through a complex network of subcellular signalling nanodomains^5^. Elevation of cAMP following G-protein–coupled receptor (GPCR) activation is not uniformly distributed within cardiomyocytes. Instead, receptor-specific, nanometer-scale cAMP gradients are established across the cell, governed by the localized activity of cAMP-hydrolyzing phosphodiesterases (PDEs)^6^. The spatially confined cAMP nanodomains activate a selected subset of PKA enzymes, typically tethered in proximity of specific phosphorylation targets via scaffolding proteins named A-kinase anchoring proteins (AKAPs)^7^. This spatial organization allows selective phosphorylation of local targets, resulting in cellular effects that exhibit a distinct profile, ensuring signalling specificity^8^. While drugs targeting the GPCR influence the entire intracellular network of the receptor-associated cAMP nanodomains and the full spectrum of its functional effects, the compartmentalized nature of cAMP signalling offers the opportunity for a more precise intervention^9^. Such an approach could selectively modulate cAMP levels within an individual compartment achieving nanometer subcellular precision and stringent functional selectivity^5^.

We previously demonstrated that the cAMP signal generated upon β-AR activation at TPNI is distinctly regulated compared to the cAMP signals that the same stimulus generates at the plasmalemma or the SR^6^. This observation prompted us to investigate whether TPNI defines a distinct cAMP nanodomain that could be selectively targeted to enhance relaxation for the treatment of diastolic dysfunction.

## Methods

### Materials & Reagents

Isoproterenol, cilostamide, rolipram, forskolin, and 3-isobutyl-1-methylxanthine (IBMX) were from Sigma-Aldrich; N-terminal stearic-scrambled peptide (SC, KTELKAERGIKSDVAKERELHDLGK) and disrupting peptide (P33, GTRAKESLDLRAHLKQVKKEDIEKE) were synthesised by GenScript. Collagenase A was from Roche; pancreatin was from Sigma; laminin (mouse) was from BD Biosciences; PBS, DMEM high glucose, MEM199, horse serum, newborn calf serum (NCS), 1% penicillin/streptomycin (10,000 units/ml of penicillin and 10,000 μg/ml of streptomycin), and L-glutamine were from Invitrogen. Adenoviral vectors and self-complementary AAV vectors carrying the target reporter and sequences were generated by Vector Biolabs (Malvern, PA, USA).

### Generation of MyBPC-CUTie targeted reporter

Full length mouse cardiac myosin binding protein C (MyBPC, NM_008653.2) was PCR-amplified to incorporate NheI restriction sites at both 5’ and 3’ ends. The resulting fragment was cloned, in frame, into the NheI site of the CUTie sensor^6^. Clones were screened for correct orientation of insert and selected clones were sequenced. Adenoviral vector carrying the MyBPC-CUTie targeted reporter was generated by Vector Biolabs (Malvern, PA, USA)

### Primary cardiac myocyte isolation

ARVMs were isolated from 300-350 g male Sprague Dawley rats as described^10^. Briefly, hearts were perfused on a Langendorff apparatus at 37 °C for 4-5 min through the coronary arteries with an oxygenated Calcium-free Tyrode Buffer solution (130 mM NaCl, 5 mM Hepes, 0.4 mM NaH_2_PO_4_, 5.6 mM KCl, 3.5 mM MgCl_2_, 20 mM Taurine, 10mM Glucose), and subsequently digested with 0.06 mg/ml Liberase TH (Roche Diagnostics Limited, UK) in a 10 μM Calcium Tyrode Buffer solution with addition of 10 μM Blebbistain (T6038, TargetMol Chemicals, USA) for a further 15-20 min until the heart became flaccid. After perfusion, the left ventricle was removed, minced, and resuspended in equal volume of 1% BSA Calcium-free Tyrode Buffer solution. Extracellular Calcium was added incrementally to reach a final concentration of 1.0 mM. Cells were cultured in Minimum Essential Medium (MEM) (Sigma-Aldrich, UK), supplemented with 2.5% foetal bovine serum (FBS), 1% penicillin-streptomycin, 1% L-Glutamine and 9 mM NaHCO_3_ and plated on laminin (40 μg/ml) coated culture dishes. ARVMs were left to adhere for 2 h in a 5% CO_2_ atmosphere at 37 °C, before the medium was replaced with FBS-free MEM. Myocytes were infected for 3 h at multiplicity of infection (MOI) of 10–100 with adenovirus encoding for FRET reporters. The medium was then replaced with fresh FBS-free MEM including 0.5 μM cytochalasin D (MP Biomedicals, UK). Infected cells were kept in culture for less than 36 h past infection. NRVM were cultured as described before^11^.

### Immunostaining

ARVMs plated on coverslips were infected for 2 h at a MOI of 10-100 with adenovirus encoding for CUTie sensors. After 48 h of infection, cells were washed 3x with PBS, and fixed with 4% PFA in PBS at room temperature for 15 min. After permeabilisation with PBS+0.2% Triton-X-100 (PBST) for 30 min, the coverslips were blocked in PBST+1% BSA for 1 h at room temperature. Primary α-Actinin antibody (Abcam, ab9465, 1:400) incubation was performed overnight at 4°C, followed by 3 washes at room temperature with PBS+0.025% Tween 20 and incubation with an Alexa Fluor 647 (Invitrogen, A-21235, 1:500) conjugated secondary antibody for 1 hour. Coverslips were mounted with Ibidi mounting medium (Thistle Scientific). Confocal images were acquired using a Leica Stellaris TauSTED Xstend microscope (Leica, Wetzlar, Germany), and the images were analysed using Leica LAS X software (Leica, Wetzlar, Germany).

### Primary cardiomyocytes siRNA knockdown

Silencer pre-designed PDE4B siRNA (Assay ID: 200641), PDE4D siRNA (Assay ID: 200644), and Silencer negative control No.1 siRNA (AM4611) were purchased from Thermo Fisher Scientific. 2×10^6^ NRVM were plated onto 6-well plate coated with laminin (0.75 µg/cm^2^). One day after plating the medium was replaced with low-serum (0.5% NCS) DMEM medium. Individual siRNA with indicated concentrations were transfected into NRVM using Lipofectamine RNAiMax transfection reagent (133778030, Thermo Fisher Scientific) following the manufacture’s protocol. 48 h after transfection, the cells were harvested for Western blots or FRET experiments.

### FRET imaging

FRET imaging experiments were performed 24-48 h after transduction with adenovirus carrying each sensor, as previously described^12^. For the measurements, cells were maintained at room temperature in a modified Ringer solution (125 mM NaCl, 20 mM Hepes, 1 mM Na_3_PO_4_, 5 mM KCl, 1 mM MgSO_4_, 5.5 mM Glucose, CaCl_2_ 1mM, pH 7.4). ARVMs were imaged 18 h after infection and kept at ∼35 °C in a modified Tyrode solution containing 1.4 mM Calcium. An inverted microscope (Olympus IX71) with a PlanApoN, 60X, NA 1.42 oil immersion objective, 0.17/FN 26.5 (Olympus, UK), was used for image acquisition. The microscope was equipped with a CoolSNAP HQ2 monochrome camera (Photometrics) and a DV2™ optical beam-splitter (MAG Biosystems, Photometrics). Images were acquired and processed using MetaFluor^®^ 7.1, (Meta imaging series, Molecular Devices). FRET changes were measured as changes in the background-subtracted 480nm/535nm fluorescence emission intensity on excitation at 430 nm and expressed as R/R_0_, where R is the ratio at time t and R_0_ is the average ratio of the first 8 frames.

### Immunoprecipitation

For pull-down experiments, ARVM isolated from one heart were suspended in Tyrode buffer with 1.4 μM Calcium and treated as described. For NRVM pull-down, 6×10^6^ NRVM were plated onto 10-cm Petri dishes coated with laminin (0.2 µg/cm^2^) and infected with the indicated adenovirus for 36-48 hours and treated as described. For hiPSC-CMs pull-down, 1×10^6^ hiPSC-CMs were plated to 6-cm Petri dishes 30 days after differentiation and treated as describe. For heart tissue, freshly isolated ventricles (≈50–100 mg) were homogenized on ice, while cultured cells were harvested directly in ice-cold IP buffer (50 mM HEPES pH7.4, 150 mM NaCl, 2 mM EDTA, 10% glycerol, 0.5% Triton-X-100 or 50 mM Tris pH7.4, 150 mM NaCl, 1 mM EDTA, 10% glycerol, and 1% nonyl phenoxypolyethoxylethanol [NP40]) supplemented with Complete EDTA-free protease inhibitor cocktail tablets and a ‘PhosSTOP’ Phosphatase Inhibitor Cocktail Tablet (Roche Diagnostics Limited) for 5min on ice. The cells were then collected and placed on a rotating wheel for 20 min at 4°C. Insoluble material was removed by centrifugation at 17,000g for 10 min at 4° C and total protein was quantified by Micro BCA Protein Assay Kit (Pierce Biotechnology Inc.). 500-1000 μg proteins were rotated for 1 h at 4°C with 20 μl Pierce protein A Plus agarose (Thermo Fisher Scientific, 22810). The precipitates were removed by centrifugation at 1,000g for 3 min. Supernatants were then rotated overnight with a monoclonal anti-cardiac troponin I (Abcam, ab10231, 1:250), followed by 30 μl protein G agarose beads (Cell signalling, 37478) for 2-4 h. The precipitates were than collected by centrifugation at 1,000g for 3 mins and beads were washed three times with ice-cold IP buffer. Bound proteins were then eluted in 60 μl of 2X SDS loading buffer (Life Technologies) and released from the beads at 96°C for 6 min.

### Western blotting

Pull-down proteins and samples equivalent to 20-50 μg proteins were separated on NuPage 4-12% Bis-Tris gradient gel (Invitrogen) and transferred onto PVDF blotting membrane (Amersham Hybond P 0.45 PVDF, Cytiva). After transfer, the membrane was blocked for 1 h at room temperature with 5% skim milk (VWR Chemicals) or 5% PhosphoBLOCKER blocking reagent (Cell Biolabs), and then incubated overnight at 4°C with the following primary antibodies: mouse-anti-cardiac TPNI (Abcam, ab10231, 1:2000), rabbit anti-Phospho TPNI (S22+23) (Abcam, ab190697, 1:2000), mouse anti-PLN (Badrilla, A010-14, 1:4000), rabbit anti-phospho PLN (pSer16) (Badrilla, A010-12AP, 1:4000), mouse anti-MyBPC (Santa Cruz, sc-137237, 1:500), rabbit anti-phospho MyBPC (pSer282) (kindly provided by Dr. Sakthivel Sadayappan, Department of Cellular & Molecular Medicine, University of Arizona College of Medicine, Tucson, USA, 1:5000), rabbit anti-PDE4D9, goat anti-PDE4B and goat anti-PDE4D (kindly provided by Dr. George Baillie, University of Glasgow, UK, 1:5000). The blots were washed and incubated with appropriate horseradish peroxidase-conjugated secondary antibodies for 1 h at room temperature. Immunoreactive bands were visualized by ECL western blotting detection kit. Antibody dilutions and exposure times were adjusted to ensure that for each antibody the signal detected was within the linear range. If necessary, membranes were then stripped of bound antibodies by incubation with stripping buffer (Thermo Fisher Scientific, 21059) for 30 min at room temperature with gentle agitation. For western blots analysis, the results from each gel were normalised to the control group from the same gel and the band intensities were quantified by densitometry using ImageJ software.

### qPCR

Quantitative PCR was conducted as previously described^13^. In brief, RNA extraction from cardiac tissue was performed using a mixed protocol using phenol:chlorophorm extraction followed by Qiagen RNeasy kits for cleanup. qPCR reactions were perfomed using the ABI Prism 7300 qPCR thermocycler and analysis software (60 °C for 30 s, 95 °C for 10 min followed by 40 cycles of 95 °C for 15 s, 60 °C for 1 min). Reactions were conducted in qPCR Fastmix (VWR, Lutterworth, UK), as previously described. Primers used are as in^13^.

### Peptide array

Peptide array experiments were performed by automatic SPOT synthesis as described^14–16^. Human TNI peptides were synthesised onto PEG-derivatized continuous cellulose membrane supports *via* 9-fuorenylmethyloxycarbonyl chemistry (Fmoc) using the MultiPep 2 Robot (CEM). A far western blot approach was utilised to detect PDE4D9 (derived from lysate of PDE4D9 overexpressing HEK293 cells), whereby TNI arrays (consisting of 20-25mer peptides) were (i) blocked for 2 h at room temperature in 1X Intercept (TBS) Blocking Buffer (LI-COR), (ii) incubated overnight at 4 °C in [1000 μg] PDE4D9-HEK293 lysate (diluted in 1X TBS, pH 7.4), (iii) incubated overnight at 4 °C in αPDE4D9 rabbit primary antibody (1:1000, bespoke PDE4D9 antibody raised against the unique N-term PDE4D9sequence in rabbits), (iv) incubated for 1 h at room temperature in IRDye 800nm donkey α-rabbit secondary (1:10,000, LI-COR) and (v) visualised using Odyssey CLx (LI-COR). Arrays were washed three times in 1X TBS-T following protein, primary and secondary antibody incubation steps. The antibody without lysate was used as a negative control. Antibodies were diluted in 1x Intercept (TBS-T) Antibody Diluent (LI-COR).

### *In vitro* work-loop protocol

Sprague-Dawley Rats: Sex; Male: Body Mass; 250 – 400g: Animals: (5 for P33, 4 for SC, 6 for controls) were purchased from Charles River (Margate, UK) and housed at Warwick University Biomedical Services Unit (BSU) in a 12:12 day/night cycle in rooms held at 22 °C. Cardiac myocytes were isolated as previously described^17^. Isolated cardiac myocyte work-loop experiments were performed using an adapted methods as described previously^18–20^. In brief, myocytes were placed in a perfusion chamber and carefully attached to fine glass rods (high-speed length controller or highly sensitive cantilever force transducer respectively) (IonOptix, Milton, MA, USA). Once attached fresh buffer was cycled through the chamber to remove excess cells. Isometric and force-length protocols were implemented for separate sets of cells incubated in the presence of scrambled or disrupted peptide or DMSO vehicle control. Work-Loop involved electrically stimulating an isometric contraction; the length was held constant until the afterload value was achieved; this was followed by an isotonic phase during which the cell was shortened to maintain a constant level of force. The work-loop protocol was initiated, and the myocytes were paced at 1 Hz using a field stimulator (Myopacer; IonOptix, Milton, MA). Isometric contractions were recorded for 5 min to obtain a baseline/control period. The work-loop protocol relies on the force length relationship and when mimics the cardiomyocyte length change cycle providing realistic evaluation of *in vivo* ‘like’ myocardial contractile mechanics which involve force feedback work-loops with stepwise preload increases and ramped afterload increases over time. Intervals between loop phases were filled by continuous isometric activations at 1Hz 2hrs prior to the work-loop experiment and the maximal isometric force (Fmax) and the relaxation velocity were recorded.

### Fura-2 measurements

Isolated ARVMs were plated onto laminin (2 μg/cm^2^)-coated coverslips and cultured overnight. Cells were loaded with 1 μM Fura-2 AM (Molecular Probes, OR, USA) in the dark at 37°C for 15 min and then washed three times. Coverslips were mounted in the perfusion chamber, continuously perfused with a Tyrode solution containing 1.4 mM Calcium at 37 °C, and paced at 1 Hz using a field stimulator (Myopacer; IonOptix, Milton, MA). When at steady state, cells were perfused with ISO 5 nM for about 10 min. The background-subtracted Fura-2AM emission at 510 nm was recorded and expressed as a ratio of fluorescent light on excitation at 340 and 380 nm (R340/380). The recorded transients were analysed using the IonWizard software (IonOptix, Milton, MA).

### hiPSC-derived cardiac myocytes

Monolayer differentiation of the PGP1 hiPSC cell line^21^ was performed via Wnt pathway modulation with small molecule inhibitors. Once stem cell confluence reached ∼80%, the cells were induced to form cells of the mesodermal layer with 5μM CHIR99021in RPMI1640/ B27 minus insulin for 24 h. This was day 0 of differentiation. At the end of the 24 h, the medium was replaced with RPMI1640/B27 minus insulin. 48 h later, the medium was replaced with 5 µM IWP2 in RPMI1640/B27 minus insulin for 48 h. Subsequently the medium were replaced with fresh RPMI1640/NB27 minus insulin every 48 h until day 10, when the medium was changed to glucose-free RPMI1640/B27 plus insulin supplemented with 4mM lactate for two 48-h cycles. Spontaneous contraction was observed on day 8-10 of differentiation.

On day 20, iPSC-CMs were dissociated with 10x TrypLE enzyme for 12 min. The dissociation reagent was quenched with plating media (20% fetal bovine serum, 10 μM Thioglycerol in RPMI 1640 supplemented with B27 plus insulin). Cells were collected in suspension, centrifuged at 180g for 3 min and resuspended in plating media. Cardiomyocytes were plated at a density of 1 × 10^6^ on Matrigel-coated Ibidi 35mm imaging plates. On day 28 cardiomyocytes were transduced with Adenovirus-RGECO^22^. On day 29, a second transduction was carried out with Adenovirus-α-Actinin-GFP to enable simultaneous Calcium and contractility imaging.

### Peptide treatment of iPSC-CMs

On day 30 cardiomyocytes were incubated for 15 min with P33 or scrambled peptide (SC). Controls were provided by equivalent concentrations of dimethylsulfoxide (DMSO). This was repeated in four biological replicates (separate differentiations). A minimum of 30 cells expressing both fluorescent reporters were recorded per plate. Time lapse videos acquired at 1 Hz electrical pacing at 20V and 5msec bipolar current.

Imaging was performed with a Nikon Ti-2 Eclipse inverted microscope (Nikon, Japan) at x100/1.45NA magnification with a Nikon Plan Apo λ (Nikon, Japan). Excitations at 550nm (RGECO) and 470nm (a-Act-GFP) were provided with a CooLED pE4000 (CooLED, UK). Fluorophores were imaged simultaneously using dual illumination through a Quad-Filter (LED-DAPI/FITC/TRITC/Cy5, Semrock, USA). Emissions were passed through a TwinCam dual camera splitter (Cairn Research, UK), split with a T565lpxr and filtered through an ET632/60 (RGECO) or ET520/40 (GFP) filter (Semrock IDEX, NY, USA). Emissions were collected by two aligned Teledyne Photometrics Kinetix sCMOS cameras (Teledyne Photometrics, USA). Time lapses were acquired at 10 ms exposure (100fps) at 37°C.

Time lapse videos were analysed for Calcium using the published computational pipeline CalTrack^22^. Contractility videos were uploaded into the University of Oxford Advanced Research Computing (ARC) facility cluster and analysed using the automated motion-tracking pipeline SarcTrack^23^.

### AAV Vector Design and Production

Self-complementary adeno-associated virus serotype 9 (scAAV9) vectors were generated by Vector Biolabs (Malvern, PA, USA). Transgene expression is driven by the cardiac troponin T (cTnT) promoter to ensure cardiomyocyte-specific expression. Two constructs were generated: scAAV9-cTnT-SC-IRES-mCherry-WPRE3 and scAAV9-cTnT-P33-IRES-mCherry-WPRE3. The The SC and P33 inserts encode the sequence of 25 amino acids, using the nucleotide sequences 5’-*ATGAAGACCGA ACTCAAGGCCGAGCGGGGGATCAAGTCCGACGTGGCCAAGGAAAGGGAGCTCCACGACCTGCAGAAGTGA-3’* and 5’-*ATGGGGACCCGGGCCAAGGAATCCTTGGACCTGAGGGCCCACCTCAAGCAG GTGAAGAAGGAGGACATTGAGAAGGAATGA-3’*, respectively. For both vectors, an internal ribosome entry site (IRES) was placed downstream of the primary open reading frame to enable bicistronic expression of mCherry. A WPRE3 element was included to enhance transcript stability and expression efficiency^24,25^.

### Surgery

All in vivo experiments were approved by the Committee for Animal Care and Ethical Review at the University of Oxford and comply with the UK Animals (Scientific Procedures) Act 1986, as amended 2012. C57BL/6J^OlaHsd^ mice (Envigo, Huntingdon, UK) were group housed in individually ventilated cages under specific pathogen-free conditions at 21 °C with controlled humidity of 55 % and a 13–11 h light-dark cycle.

Water and chow were available ad libitum using irradiated 2016 Teklad Global 16% Protein Rodent Diet. TAC and MI surgery was performed aseptically as described previously^26^ under isoflurane general anaesthesia (2% in medical oxygen with anesthetic depth assessed by loss of pedal reflex)^27^. Buprenorphine analgesia was given subcutaneously immediately prior to surgery (0.8 mg/Kg) and again the morning after surgery (0.4 mg/Kg). MI surgery was performed in female mice, as male mice showed a higher incidence of sudden cardiac rupture within the first week after surgery (approximately 25 %) compared with females (approximately 5 %), consistent with prior reports. The use of female mice therefore improved postoperative survival and reduced variability in infarct size. All the MI mice were inspected and found to have large infarcts.

### Echocardiography and haemodynamics

Mouse transthoracic echocardiography was performed using a Visualsonics Vevo 3100 ultrasound system (Visualsonics, Toronto, Canada) equipped with 22–55 MHz transducer (MS 550D). Mice were maintained on isoflurane anaesthesia (1.0–1.5% in medical O2), placed on a homeothermic table in a supine position. B-mode images were obtained in a parasternal short and long-axis view and dimensions of the left ventricle were measured in a short-axis view in diastole and systole. Diastolic functions were obtained in Apical four-chamber with colour flow imaging for optimal alignment of pulsed-wave Doppler (PWD) and Tissue Doppler imaging (TDI). Data analysis was performed by a single operator blinded to genotype using Vevo Lab, Edition 3.1.1 (Visualsonics, Toronto, Canada). Retrograde LV cannulation was performed under isoflurane anaesthesia via the right carotid artery using a 1.4F solid-state pressure catheter (SPR-839, Millar Instruments, Texas, USA) as described previously^28^. The jugular vein was cannulated using flame-stretch polyethylene tubing (Portex 0.96 mm OD, 800/100/200, Smiths Medical, UK) for administration of dobutamine hydrochloride at 32 ng/g BWt/min. Hearts were removed under anesthesia at the end of the protocol.

### Histology

For histological analyses, mouse hearts were harvested and fixed in 4% paraformaldehyde (PFA) at room temperature overnight. For paraffin embedding, samples were dehydrated using a series of graded ethanol concentrations (50-100%, each for at least 2 h), and cleared in butanol overnight. Samples were incubated at 60° C in 1:1 butanol: molten pastillated fibrowax for 30 mins, then three times in 100% molten wax, each for 4 h to overnight, before embedding and sectioning (10 μm transverse sections). Paraffin sections were stained with the Trichrome Stain Kit (AB150686, Abcam) following the manufacture’s protocol. Slides were imaged with a Nanozoomer S210 Slide Scanner (Hamamatsu Photonics, Hamamatsu, Japan) and analysed using Qupath-0.6.0^29^.

### Human ventricular samples, isolation and culture of human ventricular myocytes

Human ventricular samples were obtained from patients with and without heart failure and reduced ejection fraction at the time of transplantation, valvular surgeries or myectomy and informed consent was obtained from the donors. The study was conducted in accordance with the Declaration of Helsinki and approved by the Ethical Committees of the Aerztekammer Hamburg (WF-088/18). After surgical excision, tissue samples were placed into Custodiol® solution (Dr. Franz Köhler Chemie GmbH) and transported to the laboratory. Of the 26 samples obtained, 10 were used for western blot (Extended Data table 6), and 16 for myocytes isolation (Extended Data table 5). Human ventricular myocytes were isolated as previously described (doi: 10.1093/eurheartj/ehad086). Briefly, tissue was cut into small pieces and digested in Stop Ca^2+^ free solution (containing in mM: 88 Sucrose, 88 NaCl, 5.4 KCl, 4 NaHCO3, 0.3 NaH2PO4, 1.1 MgCl2, 10 HEPES, 20 Taurine, 10 Glucose, 5 Na+ pyruvate. 7.4 pH at room temperature; 5% BSA, and 12 µM blebbistatin) plus 0.5 mg ml-1 collagenase (Worthington type 1, 240 U), 0.5 mg ml-1 proteinase (Sigma type XXIV, 11 U) and 2% bovine serum albumin (BSA; Sigma, St. Louis, Missouri, USA) by spinning it at 200 rpm and 37 °C 30 min. Then tissues were transferred to Stop Ca^2+^ free solution alone and moved up and down in a Pasteur pipette for mechanical dissociation. The remaining tissue was digested for 5 more rounds in Stop Ca^2+^ free solution with 0.4 mg ml-1 collagenase and 2% BSA at 200 rpm and 37 °C 15 min, until the tissue was completely digested. Human ventricular myocytes were harvested by centrifugation at 500 rpm during 5 min and resuspended in MEM medium (M 4780; Sigma-Aldritch) containing 2 mmol/L Ca^2+^, 2.5% fetal bovine serum (FBS, Invitrogen), 1% penicillin-streptomycin and 12 µM blebbistatin. Myocytes were then plated on Laminin-coated dishes (Cellvis). After 2 h at 37 °C and 5% CO_2_, medium was changed to FBS-free MEM medium containing adenovirus (MOI 200 PFU/cell) and keep them in culture at 37°C and 5% CO_2_ until FRET experiments were performed 24-48 h after.

### Statistical analysis

Statistical analysis was performed using GraphPad Prism 10 and R. For western blot analyses, band intensity values for each sample were normalised to the corresponding control group on the same gel. Comparisons involving three or more independent groups were analysed using one-way Welch’s ANOVA, followed by the Dunnett T3 post-hoc test. Two-group comparisons were analysed using unpaired Welch’s t-test, or paired t-test, as appropriate. When multiple pairwise comparisons were performed within the same comparison family, p-values were adjusted using the Benjamini–Hochberg procedure. All t-tests were two-tailed. Statistical significance was defined as p < 0.05. When providing summary statistics, X ± Y represents the mean (X) and standard deviation (Y). For experiments using human cardiomyocytes, data are expressed as x/y, where x denotes the total number of individual cells analysed and y denotes the number of independent patient-derived samples. Each patient (y) represents one biological replicate, and multiple cells (x) were analysed per sample.

## Results

### Distinct cAMP regulation at TPNI

We first sought to confirm whether TPNI designates a nanodomain at the myofilament where cAMP is uniquely regulated. For this, we used the FRET-based cAMP reporter TPNI-CUTie^6^ and the newly developed MyBPC-CUTie (Fig. 1a,b) to monitor the cAMP signal at TPNI and MyBPC, a PKA target proximal to TPNI on the myofilaments (Fig. 1b). Correct localisation of TPNI-CUTie and MyBPC-CUTie was confirmed by co-immunoprecipitation and immunostaining (Fig. 1c-e). Overlapping kinetics and maximal amplitudes of the cAMP-dependent FRET changes at saturating cAMP (Fig. 1f) allowed direct comparison of the FRET change reported by the two sensors. Adult rat ventricular myocytes (ARVM) expressing TPNI-CUTie, MyBPC-CUTie or the previously characterised AKAP18δ -CUTie^6^, a sensor targeted to the SR and forming a complex with PLB and SERCA (Fig. 1b,c), were then challenged with the β-AR agonist isoproterenol (ISO, 5nM). Consistent with our previous findings^6^, the cAMP response at AKAP18δ was faster, significantly larger and sustained for at least 5 min compared to the cAMP response at TPNI, where cAMP elevation was slower and transient, peaking at 1 min after addition of ISO and returning close to baseline after 5 min (Fig. 1g,h). Remarkably, the cAMP response at MyBPC was also distinct, with maximal amplitude and velocity similar to TPNI at 1 min but with sustained kinetics for at least 5 min (Fig. 1g,h). Similar differences were confirmed in neonatal rat ventricular myocytes (NRVM) (Extended Data Fig. 1). The time course of PKA-dependent phosphorylation showed a consistent divergence in kinetics, with TPNI phosphorylation peaking at about 5 min after the application of 5 nM ISO and declining significantly after 10 min, reflecting the expected delay due to the time required for inactivation of PKA and subsequent dephosphorylation of the TPNI site after cAMP levels decline. In contrast, phosphorylation of MyBPC and PLB showed a steady increase up to 10 min after ISO application (Fig. 1i-j). Thus, β-AR activation generates a cAMP signal at TPNI that is transient in nature and clearly distinct from the signal generated at the SERCA/PLB complex and at MyBPC, despite the close proximity of these targets.

### A PDE4D uniquely modulates cAMP at TPNI

We previously demonstrated that the transient cAMP response to ISO at TPNI is abolished in the presence of the non-selective PDE inhibitor 3-isobutyl-1-methylxanthine (IBMX), resulting in a sustained elevation of the second messenger at this site^6^. Proteomic studies indicate that both PDE2A and PDE3A isoforms are associated with cardiac myofilaments^30^. To assess whether either of these enzyme families is responsible for the unique transient nature of the cAMP signal at TPNI, NRVM expressing TPNI-CUTie or MyBPC-CUTie were treated with 1 nM ISO and either the PDE2A inhibitor BAY 60-7550 (BAY, 10 nM) or the PDE3 inhibitor cilostamide (Cilo, 10 μM), and local cAMP levels recorded over time. Elevation of the FRET signal on inhibitor application indicates that both PDE2A and PDE3A contribute to cAMP hydrolysis at both locations (Fig. 2a,b,e). However, both inhibitors failed to eliminate the transient nature of the response at TPNI (Fig. 2a,b,e), even upon simultaneous inhibition of the two enzymes (Fig. 2c). By contrast, inhibition of PDE4 with rolipram (Roli 10 μM) resulted in a sustained response at TPNI (Fig. 2d,e), indicating a PDE4 isoform is the main determinant of the transient cAMP elevation at this location.

**Figure 2.**
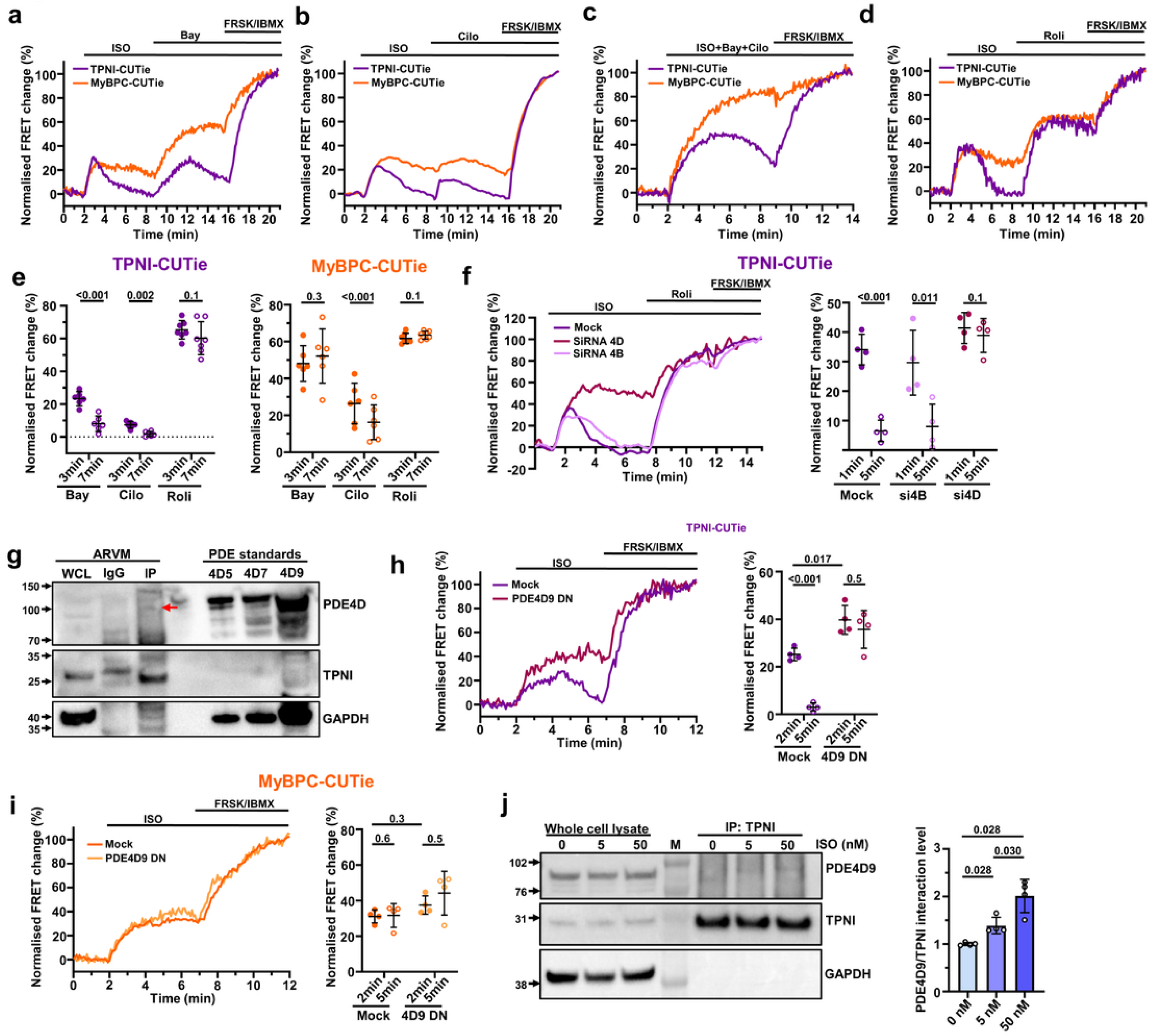
PDE4D9 regulates cAMP at TPNI. Representative kinetics of FRET change recorded in NRVM expressing TPNI- or MyBPC-CUTie and treated with 1 nM ISO followed by either (**a**) BAY (10 nM), (**b**) cilostamide (10 μM), (**c**) BAY (10 nM) + cilostamide (10 μM), or (**d**) rolipram (10 μM). (**e**) Summary of cAMP responses measured at 3 and 7 min after inhibitor application in experiments shown in (a), (b), and (d). Statistical significance was assessed using paired t-tests. (**f**) Representative kinetics and summary of FRET change recorded at 1 and 5 min following stimulation with 1 nM ISO in NRVM expressing TPNI-CUTie following siRNA-mediated knockdown of PDE4D or PDE4B. Cells were subsequently treated with rolipram (10 μM) and saturating stimulation (forskolin 25 μM and IBMX 100 μM). Statistical significance was assessed using paired t-tests. (**g**) Representative Western blot of TPNI immunoprecipitates probed with a pan-PDE4D antibody. Lysates obtained from CHO cells overexpressing PDE4D5, PDE4D7, and PDE4D9 are shown as molecular weight references. (**h**) Representative kinetics and summary of FRET change recorded at 2 and 5 min following stimulation with 1 nM ISO in NRVM expressing a dominant-negative PDE4D9-mCherry construct and either TPNI-CUTie or (**i**) MyBPC-CUTie. Statistical significance was assessed using paired t-tests and Welch’s t-tests with Benjamini–Hochberg correction, as appropriate. (**j**) Representative Western blot of whole-cell lysates (WCL) and TPNI immunoprecipitates (IP), with quantification of PDE4D9 associated with TPNI following treatment with the indicated concentrations of ISO. Statistical significance was assessed using Welch’s t-tests with Benjamini–Hochberg correction. Each data point represents an independent biological replicate. Data are presented as mean ± SD. n ≥ 4 biological replicates per group.

Of the four PDE4 families (PDE4A, PDE4B, PDE4C, and PDE4D), PDE4B and PDE4D are the most abundantly expressed in rat cardiac myocytes^31^. To identify the PDE4 subfamily involved, small interference RNA was applied to NRVM to knock down (KD) either PDE4B or PDE4D subfamilies^32^ (Extended Data Fig. 2). While the cAMP response to ISO at TPNI was not affected by PDE4B KD, PDE4D KD resulted in a sustained response (Fig. 2f). Given the extremely localised nature of the cAMP regulation at TPNI, we next investigated whether a PDE4D enzyme might be in a complex with TPNI. Immunoprecipitation of TPNI from ARVMs co-precipitated PDE4D enzymes with a molecular weight consistent with PDE4D9 (Fig. 2g). As PDE4D9 migrates at a similar size as PDE4D3 and PDE4D8^33^, we used quantitative PCR to distinguish between them and found that PDE4D9 is the most abundantly expressed in NRVMs (Extended Data Fig. 3). We therefore focused our subsequent analyses on this enzyme. The interaction of TPNI and PDE4D9 was further confirmed by co-expression of the two proteins in HEK-293 cells and co-immunoprecipitation (Extended Data Fig. 4) and by microscale thermophoresis (MST) analysis of purified TPNI and PDE4D9 (Extended Data Fig. 5). To confirm that PDE4D9 operates at TPNI in cardiac myocytes, we expressed a catalytically inactive mutant of PDE4D9 in NRVM. This mutant acts as a dominant-negative (DN) construct that displaces the native enzyme from its anchoring sites, thereby enhancing ISO-induced cAMP signaling at the location where the endogenous enzyme is normally localised^34^. As shown in Fig. 2h, expression of PDE4D9-DN resulted in a larger and sustained cAMP response to ISO at TPNI relative to control, while no significant difference was observed at MyBPC (Fig. 2i). Notably, we found that the amount of PDE4D9 detected in the TPNI immunoprecipitate was dependent on the extent of β-AR activation (Fig. 2j). Samples treated with ISO also displayed an upward band shift of PDE4D9 present in the TPNI immunoprecipitate (Fig. 2j). Long PDE4 isoforms, which include PDE4D9, undergo SUMOylation following β-AR activation, a modification that enhances their cAMP-hydrolytic activity and appears as an upward band shift in western blots^35^. To test whether SUMOylation may account for the shifted band of the PDE4D9 associated with TPNI, ARVM whole-cell lysates and TPNI pull-downs from untreated or 50 nM ISO-treated NRVM were prepared in the presence of N-ethylmaleimide (NEM) to preserve SUMOylation and probed with PDE4D9- and SUMO-1-specific antibodies after western blotting. As shown in Extended Data Fig. 6, PDE4D9 appeared as a doublet in whole-cell lysates, with the upper band corresponding to the SUMO-1-positive species. This upper band was the prevalent band in the TPNI pull-down and was further enriched following ISO treatment. Together, these findings are consistent with β-AR activation promoting recruitment of PDE4D9 to the troponin complex through a mechanism that involves PDE4D9 SUMOylation.

### PDE4D9 directly interacts with TPNI

To further confirm direct interaction of TPNI with PDE4D9 and to explore the interacting surface, we used peptide array technology^36^. TPNI arrays consisting of 20- (Extended Data Fig. 7a) or 25-mers (not shown) covering the entire sequence of rat TPNI, were probed with purified GST-tagged PDE4D9. A strong interaction was found for several peptides that were further analysed by alanine scanning, where each of the amino acids in the selected peptide was substituted with alanine or, if originally an alanine, with glycine. Peptide alanine scanning of peptide 33 from the original array identified lysine residues at position K175, K178, K179 (Extended Data Fig. 7b) and K184 (not shown) as required for interaction with GST-PDE4D9. Interestingly, a lysine to glutamate mutation at position 178 of human TPNI (corresponding to rat K179), causes an inherited restrictive cardiomyopathy^37^. To confirm the role of this residue in the TPNI/PDE4D9 interaction, we introduced the K178E mutation in TPNI-CUTie (TPNI_K178E_-CUTie) (Extended Data Fig. 8a,b) and probed the ability of the mutated TPNI moiety in this sensor to interact with PDE4D9 compared to TPNI-CUTie. As shown in Extended Data Fig. 8c, the K178E mutation significantly reduced this interaction and, when used to measure cAMP levels, TPNI_K178E_-CUTie detected a significantly larger and sustained cAMP signal in response to ISO stimulation compared to TPNI-CUTie (Extended Data Fig. 8d). Comparable results were obtained using a TPNI-CUTie mutant (TPNI_mut_-CUTie) carrying the mutations K177A, K178A and K183A (corresponding to rat K184) (Extended Data Fig. 8a-d). Consistently, ISO treatment of NRVM expressing TPNI_mut_-CUTie resulted in sustained phosphorylation of the TPNI moiety, in clear contrast to the transient phosphorylation of TPNI-CUTie (Extended Data Fig. 8e). Collectively, the data show that a carboxyl-terminal region of TPNI encompassing K177-K183 interacts with PDE4D9, and the TPNI peptide P33, with sequence G_161_TRAKESLDLRAHLKQVKKEDIEKE_185_ (Extended Data Fig. 8f), was selected for further studies.

### P33 selectively boosts local cAMP levels and phosphorylation of TPNI

We next assessed whether P33 can displace PDE4D9 from TPNI in cardiac myocytes. Pre-treatment of isolated NRVM with stearylated P33 (10 μM) for 2 h significantly reduced TPNI/PDE4D9 co-immunoprecipitation compared to treatment with a control peptide with scrambled amino acid sequence (SC, 10 μM), both in unstimulated cells and upon ISO stimulation (Fig. 3a,b). Consistently, cAMP levels measured in NRVM expressing TPNI-CUTie and pre-treated with P33 showed increased amplitude and duration of the cAMP signal compared to cells pretreated with vehicle DMSO or SC peptide (Fig. 3c). P33 had no effect on the hydrolytic activity of PDE4D9 (Extended Data Fig 9), confirming that the local elevation of cAMP at TPNI results from P33 displacing the cAMP-hydrolysing enzyme from the troponin complex. The effect of P33 is highly specific for TPNI, as no difference in the cAMP response between SC and P33 pre-treatment was observed at MyBPC (Fig. 3d) or AKAP18δ (Fig. 3e), where the response at 5 min was significantly higher than at 1 min upon either treatment and no different from DMSO control. Consistently, treatment with P33 resulted in a sustained phosphorylation of TPNI in response to ISO, while it had no effect on the phosphorylation of MyBPC or PLB (Fig 3f-g). As expected, the enhanced TPNI phosphorylation after P33 treatment increased the myofilament calcium sensitivity, as determined by force-Calcium relations measurements (Extended Data Fig 10). In cardiac myocytes differentiated from human inducible pluripotent stem cells (hiPSC-CM), where the level of TPNI phosphorylation in response to ISO stimulation also showed a transient elevation (Extended Data Fig. 11a), treatment with P33 significantly enhanced TPNI phosphorylation (Extended Data Fig. 11b) and significantly reduced TPNI/PDE4D9 interaction (Extended Data Fig. 11c,d) compared to treatment with SC control.

**Figure 3.**
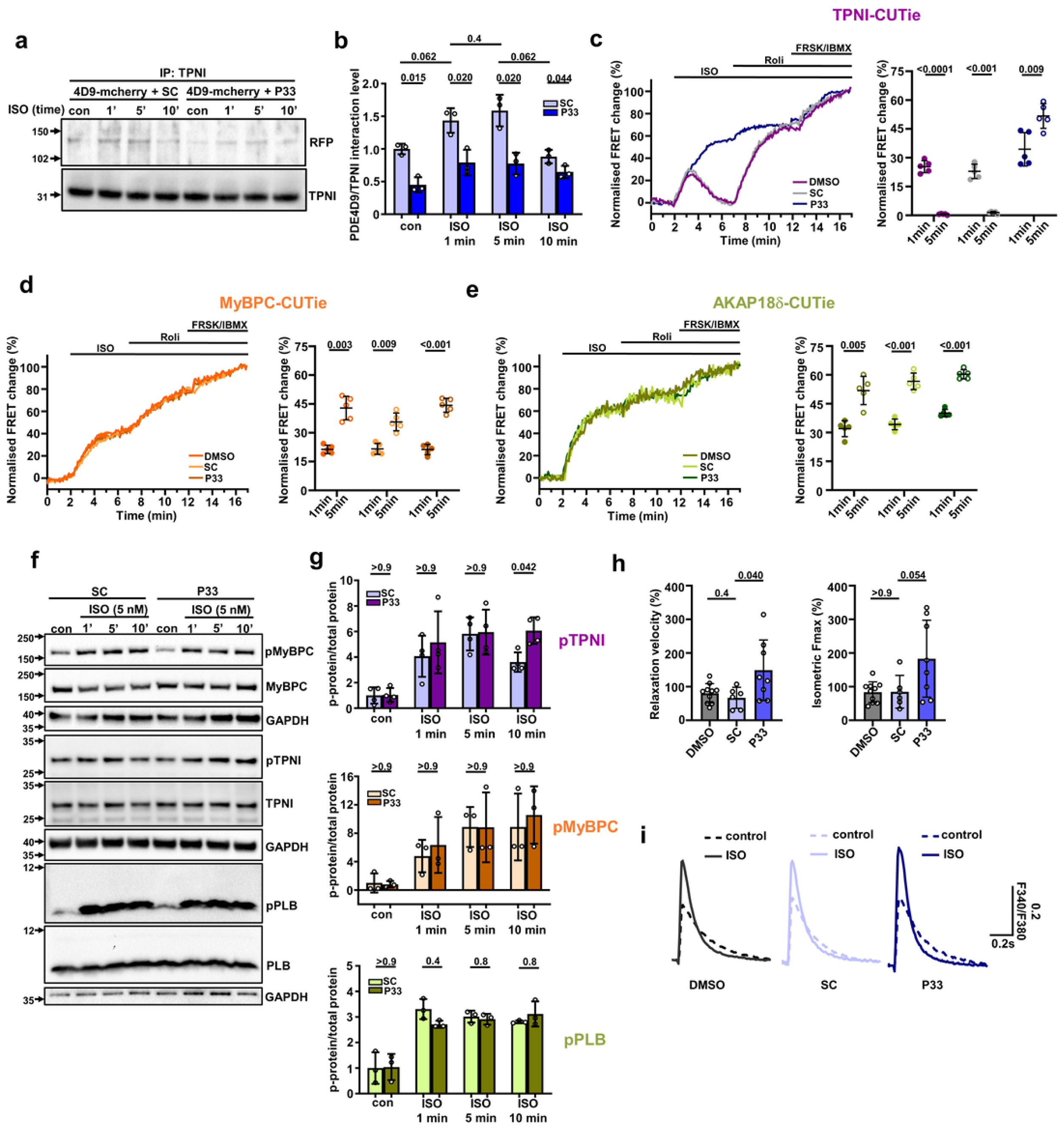
P33 selectively enhances cAMP at TPNI and improves cardiac contractility *in vitro*. (**a**) Representative Western blot and (**b**) quantification of co-immunoprecipitation from NRVM overexpressing PDE4D9-mCherry treated with ISO (1 nM) and P33 peptide (10 μM) or scrambled control peptide (SC, 10 μM). Protein interaction was assessed under basal conditions and following ISO stimulation for 1, 5, or 10 min. Statistical significance was assessed using Welch’s t-tests with Benjamini–Hochberg correction. Representative kinetics and summary of FRET responses recorded at 1 and 5 min in NRVM overexpressing (**c**) TPNI-CUTie, (**d**) MyBPC-CUTie, or (**e**) AKAP18δ-CUTie following stimulation with ISO (1 nM). Cells were pre-treated with DMSO, SC (10 μM), or P33 (10 μM), as indicated. Statistical significance was assessed using paired t-tests. (**f**) Representative Western blot and (**g**) quantification showing total protein expression and phosphorylation time courses of TPNI and MyBPC in ARVM pre-treated with P33 peptide (10 μM) or SC peptide (10 μM) and lysed 1, 5, or 10 min after stimulation with ISO (5 nM). Statistical significance was assessed using Welch’s t-tests with Benjamini–Hochberg correction. (**h**) Maximal isometric force response and relaxation velocity measured in isolated ARVM. Prespecified comparisons between DMSO and SC and between SC and P33 were analysed using Welch’s t-tests. (**i**) Average calcium transients recorded in ARVM under basal conditions or following stimulation with ISO (5 nM). Cells were pre-treated with DMSO, SC (10 μM), or P33 (10 μM), as indicated. Representative traces were generated from paced transients (≥10 contractions) recorded from a single cell and normalised to the fluorescence ratio at time 0 s. Each data point represents an independent biological replicate. Data are presented as mean ± SD. n ≥ 3 biological replicates per group.

Treatment with P33 did not affect the cAMP response to ISO measured at the plasmalemma using the targeted AKAP79-CUTie sensor^6^ (Extended Data Fig. 12a,b), or in the bulk cytosol using the untargeted cytosolic CUTie sensor^6^ (Extended Data Fig. 12c,d). Moreover, P33 had no impact on global PDE4 or PDE activity, as measured with the cytosolic cAMP sensor EPAC-H187^38^ upon application of rolipram (10 μM) or IBMX (100 μM), respectively (Extended Data Fig. 12e,f).

### P33 enhances relaxation and contraction *in vitro*

Given the observed effect of displacing PDE4D9 with P33 on TPNI phosphorylation, we expected treatment with P33 to enhance relaxation and force generation^39–43^. To verify this, we measured the velocity of relaxation using a work-loop technique in single isolated ARVM upon pacing at 1Hz^19,20^. We measured an average 2.2-fold increase (149.07 ± 89.90 for P33 vs 67.00 ± 31.29 for SC) in relaxation velocity in samples treated with P33 compared to SC (Fig. 3h). This was accompanied by an average 2.1-fold increase (183.23 ± 114.26 for P33 vs 84.77 ± 48.66 for SC) in isometric contractile force (Fig. 3h). When P33 was used to pre-treat hiPSC-CM, we found enhanced sarcomere length at relaxation (Extended Data Fig. 13a) and enhanced contraction (Extended Data Fig. 13b). Notably, the effects of P33 on lusitropy and force generation were independent of changes in the amplitude and duration of the Calcium transient both in ARVM and hiPSC-CM (Fig. 3i, Extended Data Fig. 13c-i and Extended Data Fig. 14).

### Aberrant PDE4D9/TPNI interaction in cardiac disease

Defective phosphorylation of TPNI has been reported in animal models of cardiac disease^44–47^ and in patients suffering from cardiomyopathies^48–52^, although the underlying aetiology remains unclear. We therefore explored whether the novel mechanism we uncovered for the selective regulation of TPNI phosphorylation may be altered in disease. We found a 5.1-fold larger amount of PDE4D9 interacting with TPNI in cardiac myocytes from a mouse surgical model of transverse aortic constriction (TAC) (Extended Data table 1 and Fig. 4a,b) relative to sham control. Similarly, we found 2.9-fold more PDE4D9 associated with TPNI in unstimulated cells obtained from failing cardiac myocytes from a mouse model of myocardial infarction (MI) compared to sham controls (Fig. 4c,d). Consistently, and in line with previous reports, we found attenuated TPNI phosphorylation compared to cells from age-matched healthy controls, while similar level of phosphorylation was found for MyBPC (Extended Data Fig. 15a,b).

**Figure 4.**
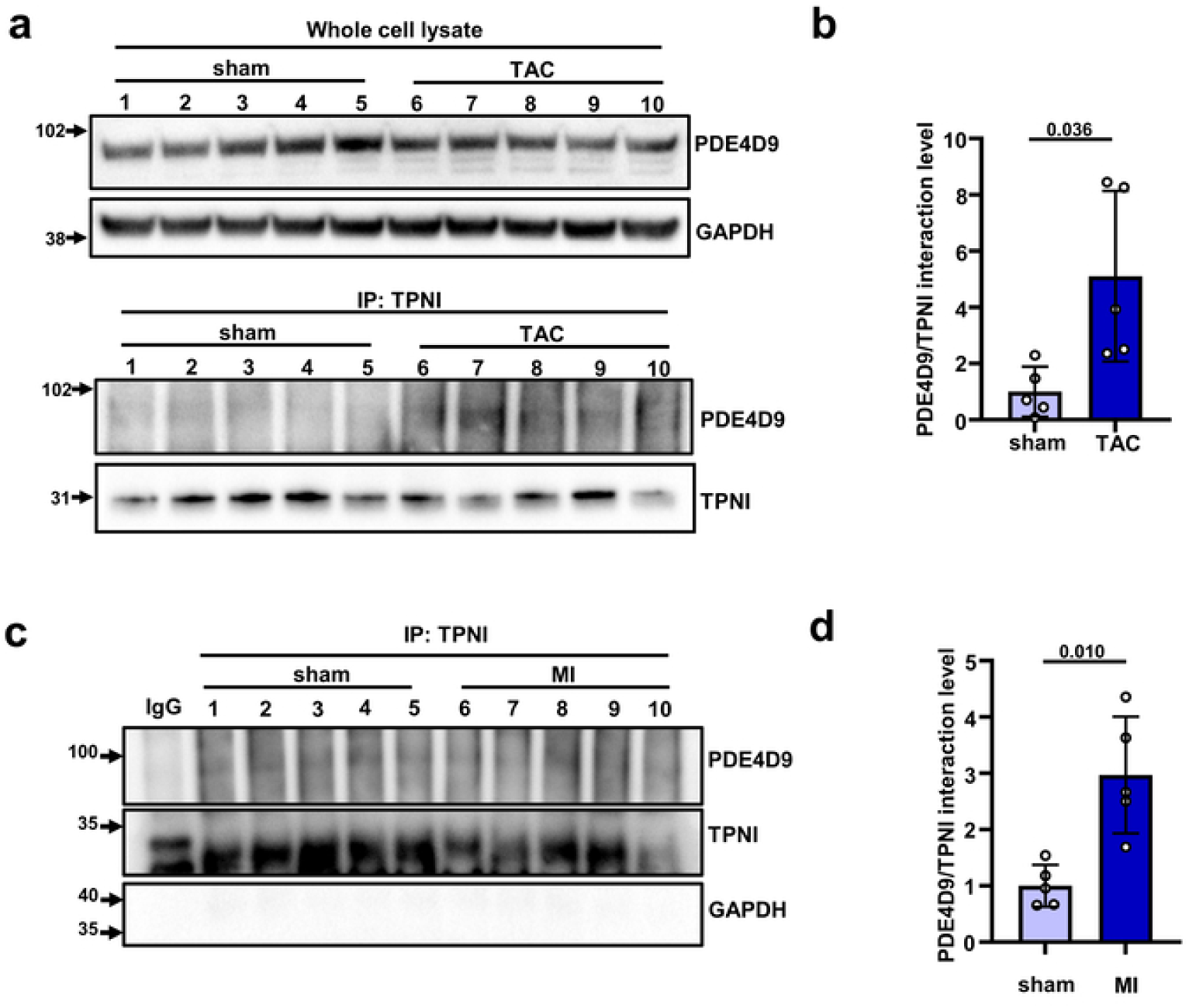
Interaction of PDE4D9 with TPNI is enhanced in cardiac disease. (**a**) Representative co-immunoprecipitation and (**b**) quantification of PDE4D9–TPNI interaction in protein lysates isolated from sham-operated or transverse aortic constriction (TAC) mouse hearts. (**c**) Representative co-immunoprecipitation and (**d**) quantification of PDE4D9–TPNI interaction in protein lysates isolated from sham-operated or myocardial infarction (MI) mouse hearts. Statistical significance was assessed using Welch’s t-tests. Lane numbers and data points represent individual animals. n = 5 animals per group. Data are presented as mean ± SD.

### P33 disrupts the TPNI/PDE4D9 interaction *in vivo*

To test whether P33 may attenuate diastolic dysfunction *in vivo*, we designed a self-complementary recombinant adeno-associated virus^53^ carrying the sequence for the control SC (scAAV9-SC) or P33 (scAAV9-P33) peptides followed by the coding sequence for mCherry under the control of an internal ribosome entry site (IRES) (Extended Data Fig. 16a). Expression of both constructs in healthy mice was confirmed by detection of mCherry in cardiac myocytes isolated 3 weeks after tail vein injection of 10^12^ vector particles (Extended Data Fig. 16b). In the same cells, efficient expression of P33 was confirmed by a 60% reduction in the amount of PDE4D9 co-immunoprecipitated with TPNI in cardiac myocytes from mice injected with scAAV9-P33 compared to scAAV9-SC (Extended Data Fig. 16c). Consistently, we found a 2-fold increase in TPNI phosphorylation in mice injected with scAAV9-P33 compared to scAAV9-SC (Extended Data Fig. 16d). Compared to scAAV9-SC, injection of scAAV9-P33 did not affect overall cardiac performance as assessed by echocardiography (Extended Data table 2 and Extended Data Fig. 17) and LV haemodynamics (Extended Data table 3) at 3 weeks post-injection.

P33 had no effect on the expression levels of PKA regulatory or catalytic subunits (Extended Data Fig. 18a,b), CaMKII expression (Extended Data Fig. 18c), or PKA- and CaMKII-dependent phosphorylation of PLB in healthy mice (Extended Data Fig. 18d,e).

### P33 improves diastolic function and preserves systolic performance *in vivo*

To test whether P33 protects from the development of diastolic dysfunction, mice were TAC or sham operated and, 1 week after surgery, tail vein injected with 10^12^ scAAV9- or scAAV9-P33 vector particles. Two separate studies, conducted at different points in time, were carried out to investigate the effect of P33 in males and female mice. Three weeks after injection, the cardiac phenotype was studied (Fig. 5a, Extended Data table 4-7). In the male cohort, a 3.5-fold increase (3.04 ± 2.11 for sham SC vs 10.91 ± 2.53 for TAC SC) in fibrosis in TAC animals was limited to a 1.7-fold increase (5.40 ± 2.28 for TAC P33) by injection of scAAV9-P33 (Fig. 5b, c). In female mice, a 6.1 fold increase (2.39 ± 0.52 for sham SC vs 14.71 ± 3.41 for TAC SC) in fibrosis was observed in TAC compared to sham operated mice injected with scAAV9-SC, which was limited to a 3.8 fold increase (9.1 ± 2.05 for TAC P33) in TAC mice injected with scAAV9-P33 (Fig. 5d and Extended Data Fig 19a). The anti-fibrotic effect of P33 does not appear to result from direct inhibition of pro-fibrotic pathways. Acute treatment of cultured cardiac fibroblasts with P33 showed no effect on TGF-β–induced SMAD2/3 phosphorylation (Extended Data Figure 20a). However, P33 normalised the elevated SMAD2/3 phosphorylation observed in cardiac myocytes from TAC mice (Extended Data Figure 20b). Together, these findings suggest that the anti-fibrotic effect of P33 is likely secondary to its direct improvement of myocardial relaxation and the consequent reduction in mechanical stress.

**Figure 5.**
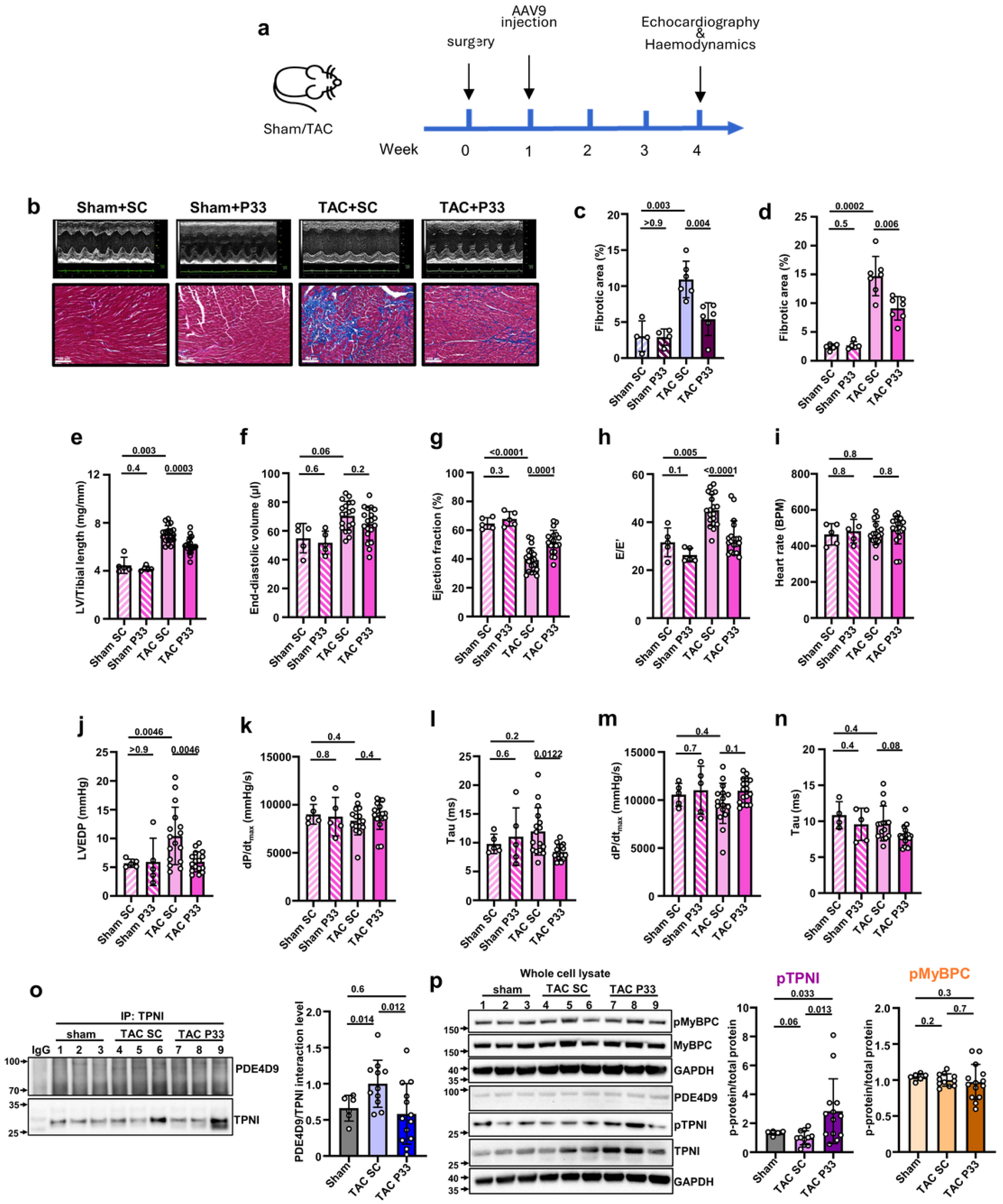
P33 protects from pressure overload-induced cardiac dysfunction. (**a**) Treatment protocol: wild-type mice were treated with scAAV9-SC or scAAV9-P33 via tail vein injection one week after sham or transverse aortic banding (TAC) surgery. Organ weights, echocardiographic parameters and left ventricular (LV) haemodynamics were obtained three weeks post-injection. (**b**) Representative M-mode echocardiography scans (top) and Masson’s trichrome staining of cardiac sections 4 weeks after TAC in male mice (scale bar = 100 μm). (**c**) Quantification of normalised fibrotic area (%) in male mice. Sham-SC, n = 5; Sham-P33, n = 5; TAC-SC, n = 7; TAC-P33, n = 7. (**d**) Quantification of normalised fibrotic area (%) in female mice. Sham-SC, n = 4; Sham-P33, n = 4; TAC-SC, n = 6; TAC-P33, n = 6. (**e**) Left ventricular weight normalised to tibial length in female mice. (**f**) Left ventricular end-diastolic volume. (**g**) Left ventricular ejection fraction. (**h**) E/e′ derived from pulse-wave Doppler and tissue Doppler imaging. Echocardiographic analyses were performed in female mice (Sham-SC, n = 5; Sham-P33, n = 5; TAC-SC, n = 19; TAC-P33, n = 19). LV haemodynamic measurements obtained under basal conditions are shown in (**i–l**) for heart rate, left ventricular end-diastolic pressure (LVEDP), maximum rate of pressure rise (dP/dtmax), and the isovolumetric relaxation constant tau. LV haemodynamic measurements obtained during dobutamine stimulation (32 ng/g BW/min) are shown in (m–n). Haemodynamic analyses were performed in female mice (Sham-SC, n = 5; Sham-P33, n = 5; TAC-SC, n = 16; TAC-P33, n = 17). Prespecified comparisons were performed between Sham-SC and Sham-P33, Sham-SC and TAC-SC, and TAC-SC and TAC-P33 using Welch’s t-tests with Benjamini–Hochberg correction. (o) Representative western blot and quantification of PDE4D9–TPNI interaction in cardiac tissue collected 4 weeks after sham or TAC surgery from male mice treated with SC or P33. (p) Representative western blot and quantification of TPNI phosphorylation levels in the same samples as in (o). Statistical significance for panels (o–p) was assessed using Welch’s ANOVA followed by Dunnett’s T3 multiple-comparison test. Lane numbers and data points represent individual animals. n ≥ 6 animals per group. Data are presented as mean ± SD.

In female mice, elevated LV weight normalised to tibial length (Fig. 5e), without evident increase in lung weight (Fig. 5e) in the TAC-SC group compared to sham-SC, indicates robust cardiac hypertrophy, in the absence of pulmonary congestion (Extended Data table 4). A trend for diastolic LV dilatation (Fig. 5f) along with a 39% lower ejection fraction (65 ± 4.0 for sham SC vs 39 ± 8.3 for TAC SC) in TAC-SC compared to Sham-SC (Fig. 5g), indicated contractile dysfunction consistent with early-stage HF. Injection of scAAV9-P33 resulted in 22% reduction in LV hypertrophy (7.07 ± 0.71 for TAC SC vs 6.12 ± 0.67 for TAC P33, Fig. 5e) and acted to preserve ejection fraction (Fig. 5g). A 1.4-fold elevation (31.59 ± 6.01 for sham SC vs 45.00 ± 6.58 for TAC SC) of E/e’ in TAC mice treated with SC, suggesting an increase in diastolic filling pressure, was reduced to a 1.1-fold increase (33.48 ± 7.09 for TAC P33) in P33 treated mice (Fig. 5h). LV haemodynamic measurements (Extended Data table 5) confirmed the beneficial effect of P33 in TAC mice. As expected, LV end-systolic pressure was greatly elevated in both TAC groups. Treatment with P33, but not SC, completely normalized LV end-diastolic pressure (Fig. 5j), in keeping with the echocardiographic E/e’ findings (Fig. 5h). Baseline values for the maximal rate of pressure rise (dP/dtmax, Fig 5k)) were not significantly different between groups, however, this parameter is sensitive to preload, so an elevated left ventricular end diastolic pressure (LVEDP) in the TAC-SC group may have maintained a deceptively high dP/dt_max_. The time constant of isovolumetric relaxation (tau) was reduced by 30% in the TAC-P33 group compared to TAC-SC (12.0 ± 4.1 for TAC SC vs 8.4 ± 1.3 for TAC P33) (Fig. 5l). The beneficial effect of P33 remained evident when mice were subjected to dobutamine stress (32 ng/g BW/min), with normalisation of the LVEDP (Extended Data table 5) and reduced values for tau (Fig. 5n). Notably, all groups were capable of increasing dP/dt_max_ upon dobutamine stimulation, indicating that P33 treatment did not blunt contractile reserve (Fig. 5m versus Fig. 5k). Comparable results were found in the male cohort (Extended Data Fig. 19b-p and Extended Data table 6-7).

In TAC mice, P33 did not alter the expression of PKA subunits (Extended Data Fig. 21a,b) or PKA-dependent PLB phosphorylation (Extended Data Fig. 21d). Although the TAC-induced increase in CaMKII expression was normalized by P33 treatment (Extended Data Fig. 21c), this was not accompanied by changes in CaMKII-dependent PLB phosphorylation (Extended Data Fig. 21e). Furthermore, P33 had no effect on CREB or NFAT phosphorylation (Extended Data Fig. 22).

Mechanistically, the beneficial effect of P33 is explained by the finding that treatment with P33 of TAC animals reduced by 40% the interaction of PDE4D9 with TPNI compared to SC peptide treatment (Fig. 5o), resulting in a 2.8-fold increase in TPNI phosphorylation (Fig. 5p). Correlation analysis confirmed a clear association between TPNI phosphorylation level and cardiac functional parameters (Extended Data Figure 23). By contrast, no effect on MyBPC phosphorylation was detected in SC or P33 treated TAC animals compared to sham (Fig. 5p).

Collectively, these findings indicate that P33 does not exert its cardioprotective effects through broad modulation of PKA, CaMKII, or downstream hypertrophic signalling pathways. Instead, the selective increase in TPNI phosphorylation observed with P33 is consistent with a local effect on PDE4D9-dependent regulation of the troponin complex, rather than a global enhancement of kinase activity within the cardiomyocyte.

### Translational applications of P33

While humans display a much slower heart rate than mice, simulations of excitation-contraction coupling using T-World^54^, a state-of-the-art virtual human cardiomyocyte, confirm a robust effect of displacing PDE4 from TPNI and normalization of diastolic function on disruption of PDE4/TPNI interaction, supporting a beneficial effect also at human-relevant heart rates (supplemental information and Extended Data Fig. 24).

Unlike rodents, human cardiac myocytes express predominantly PDE3 and relatively less PDE4 isoforms^55^. To confirm a role of PDE4 isoforms in the regulation of cAMP at TPNI in human cardiac myocytes, we compared, in myocytes obtained from HF patients (Extended Data table 6), the cAMP response on inhibition of PDE4 with Roli in the bulk cytosol, using the untargeted cAMP reporter Epac-1^56^, and at the troponin complex, using the TPNI-CUTie sensor (Fig. 6a). As expected, we found that the contribution of PDE4 isoforms to the regulation of bulk cytosolic cAMP is modest (Fig. 6b). By contrast, PDE4 inhibition had a more prominent effect (4.3-fold larger on average, 8.93 ± 7.66 for bulk cytosol vs 38.42 ± 21.76 for TPNI) on the cAMP response measured at TPNI (Fig. 6b). Similar to that was observed in rodent myocytes, the cAMP response to ISO (1 nM) was sustained at MyBPC but transient at TPNI (Fig. 6c), where subsequent inhibition of PDE4 led to a sustained elevation (Fig. 6c). As previously reported, analysis of left ventricular tissue from HF patients (Extended Data table 7) confirmed that TPNI phosphorylation is significantly reduced in failing hearts relative to healthy controls (Fig. 6d-e, Extended Data table 7). Notably, and consistent with our observations in rodents, this phosphorylation defect is explained by a prominent increase (1.8-fold on average) in the amount of PDE4D9 interacting with TPNI in HF patients relative to healthy controls, as revealed by co-immunoprecipitation experiments (Fig. 6f-g). To assess the potential beneficial effect of displacing PDE4D9 from TPNI in HF patients, we pre-treated human ventricular myocytes isolated from HF samples with 10 μM of either P33 or SC peptide for 2 h. At this time point we measured the effect on the local level of cAMP using TPNI-CUTie at 0.5 and 5 min after addition of 5 nM ISO and found a significantly larger cAMP response in P33-treated myocytes compared to SC control (Fig. 6h).

**Figure 6.**
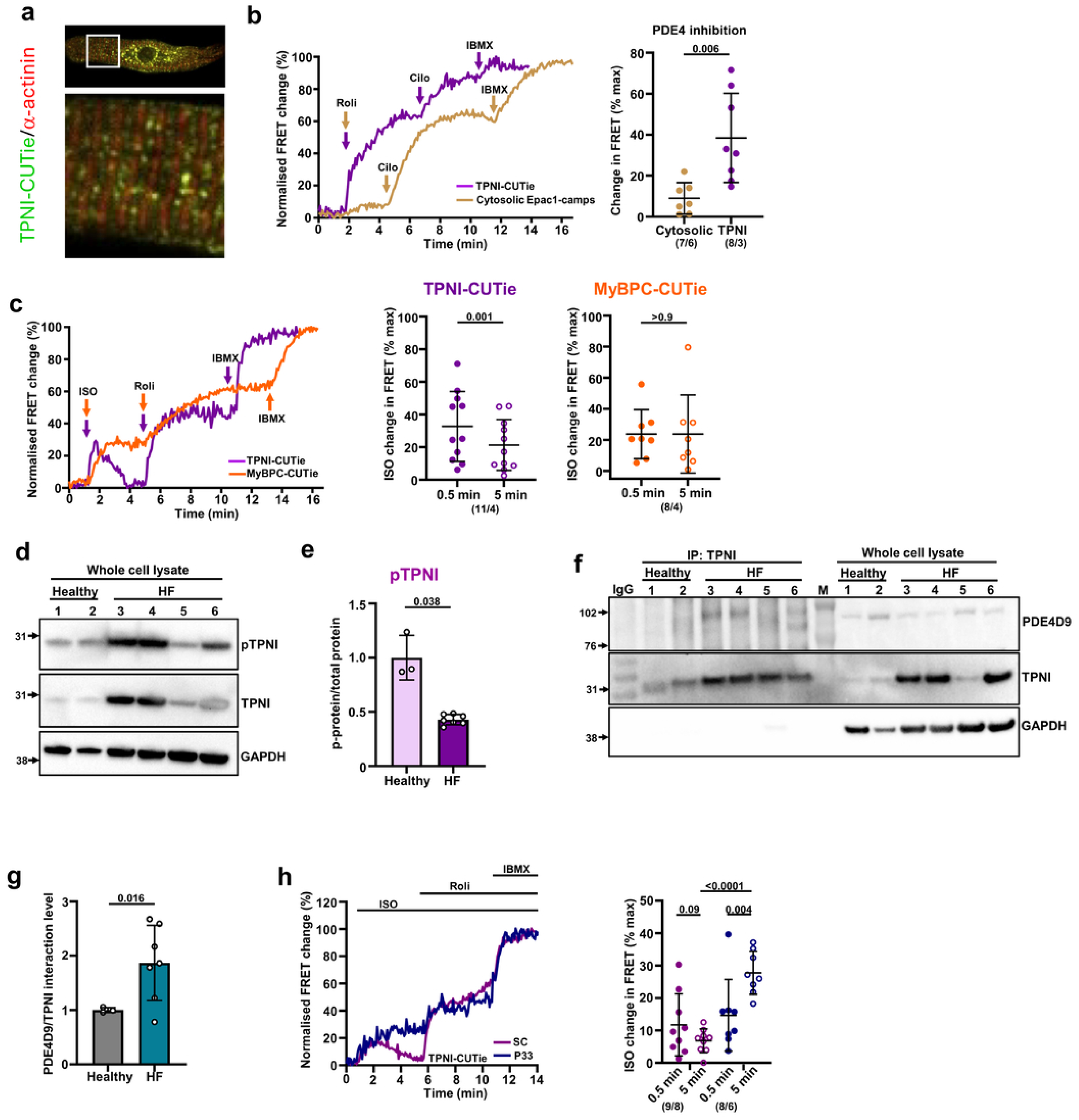
Role of PDE4D and modulation by P33 in human cardiac myocytes. (**a**) Representative image of a human ventricular myocyte expressing TPNI-CUTie (green) and stained with an α-actinin-specific antibody (red). (**b**) Representative kinetics and summary of FRET responses to the PDE4 inhibitor rolipram (10 μM) recorded in ventricular myocytes isolated from patients with heart failure with reduced ejection fraction (HFrEF) and expressing either TPNI-CUTie or the cytosolic Epac1-camps sensor. Responses to the PDE3 inhibitor cilostamide (10 μM) and IBMX (100 μM) are also shown. Compounds were added sequentially without washout. Arrows indicate the time of compound addition. Statistical significance was assessed using Welch’s t-tests. (**c**) Representative kinetics and summary of FRET responses measured at 0.5 and 5 min following stimulation with ISO (1 nM) in human ventricular myocytes isolated from HFrEF patients and expressing either TPNI-CUTie or MyBPC-CUTie. Responses to rolipram (10 μM) and IBMX (100 μM) are also shown. Statistical significance was assessed using paired t-tests. Each data point represents an independent biological replicate. Cells were obtained from four individual patients. (d) Representative western blot and (**e**) quantification of TPNI phosphorylation levels in lysates from healthy donors and patients with reduced ejection fraction. Statistical significance was assessed using Welch’s t-tests. (f) Representative western blot and (**g**) quantification of TPNI immunoprecipitates probed with a PDE4D9-specific antibody using the same samples as in (d). Statistical significance was assessed using Welch’s t-tests. Each data point represents an individual patient. Healthy donor n = 3; HFrEF n = 7. (**h**) Representative kinetics and summary of FRET responses measured at 0.5 and 5 min following stimulation with ISO (5 nM) in human ventricular myocytes isolated from HFrEF patients, pre-treated with scrambled control peptide (SC) or P33 and expressing TPNI-CUTie. Responses to rolipram (10 μM) and IBMX (100 μM) are also shown. Statistical significance was assessed using paired t-tests and Welch’s t-tests with Benjamini–Hochberg correction. Each data point represents an independent biological replicate. For both treatment groups, cells were obtained from six individual patients. The number of cells analysed is indicated below the graphs. Data are presented as mean ± SD.

## Discussion

Heart failure remains a major global health challenge, contributing substantially to morbidity, mortality, and healthcare costs worldwide^57^. Limited understanding of HF pathophysiology has so far hampered development of therapeutics that target the molecular mechanisms underpinning this condition^58^. This is particularly true for diastolic dysfunction, for which current medications are often ineffective^59^. Individuals with diastolic dysfunction face a 2–3-fold higher risk of symptomatic heart failure and a 50–60% greater risk of all-cause mortality compared to those with normal diastolic function^60,61^. Despite this substantial burden, no pharmacologic therapy currently improves diastolic function or reduces the associated mortality^62^. Extensive evidence indicates that the contractile apparatus exhibits heightened sensitivity to Calcium in end-stage HF^48,51,63^, making a strong case for targeting myofilament Calcium sensitivity for therapeutic purposes. However, proposed interventions^64–67^ are encumbered by lack of specificity and technical complexity^66,68^.

Here, we discovered a novel mechanism that modulates Calcium sensitivity by precisely regulating cAMP/PKA-dependent phosphorylation of TPNI with nanometer subcellular precision. This mechanism involves the direct interaction of PDE4D9 with TPNI, leading to localized cAMP degradation and reduced TPNI phosphorylation by PKA. The action range of the TPNI-anchored PDE4D9 is tightly confined, with its hydrolytic activity becoming ineffective beyond approximately 30 nm from the TPNI/PDE4D9 complex, where MyBPC is localised. Our findings align with the current understanding that cAMP signalling is highly compartmentalized and functions through the formation of localized signalling domains. In these structures, PKA is anchored near specific targets and is activated by confined pools of cAMP. Although not directly addressed in this study, TPNT is the likely local docking site for PKA, as it has been reported to function as an AKAP^69^.

We found that the interaction of PDE4D9 with TPNI is augmented in cardiac disese. While the mechanisms responsible remain to be established, our data suggest regulation by sympathetic stimulation and possible involvement of a SUMOylated form of PDE4D9. HF is often associated with chronically elevated sympathetic drive and aberrant SUMOylation^70^, including increased TPNI SUMOylation^71^. These mechanisms may therefore contribute to the excessive recruitment of PDE4D9 to the troponin complex observed in cardiac disease. Further studies will be required to test this hypothesis.

We demonstrate that, by selectively displacing PDE4D9 from TPNI, the highly localized nature of this regulatory mechanism can be leveraged to achieve local elevation of cAMP, local activation of PKA, and selective phosphorylation of TPNI. As expected, this phosphorylation promotes cardiac myocyte relaxation, as seen in isolated cells. Consistently, we also observed enhanced inotropy, likely due to increased crossbridge cycling rate and shortening velocity^39–41,72^. Importantly, these effects extend *in vivo*, where displacement of PDE4D9 from TPNI enhances speed of relaxation, normalises ventricular end-diastolic pressure and improves ejection fraction in a mouse TAC model of heart failure.

A key advantage of locally elevating cAMP by displacing PDE4D9 from the myofilament is the exquisite selectivity of this intervention, both at the cellular and subcellular level. TPNI expression is confined to striated muscle and PKA phosphorylates TPNI exclusively in cardiac myocytes. Thus, the effects of disrupting the TPNI/PDE4D9 interaction are expected to be highly specific to the heart. The same targeted effect of displacing PDE4D9 from TPNI could not be achieved pharmacologically, as available PDE4 inhibitors are not isoform-selective, and would inhibit all the 25 PDE4 isoforms simultaneously, both in cardiac myocytes and other tissues. We demonstrate that, instead, elevating cAMP through PDE4D9 displacement from TPNI does not affect the overall hydrolytic activity of PDE4 or other PDEs in the cell, potentially avoiding off-target effects. Moreover, displacing PDE4D9 from TPNI does not affect local cAMP signalling at the AKAP79-L-type Calcium channel complex at the plasmalemma or at the AKAP18δ-SERCA/PLB complex at the SR, supported by the fact that treatment with P33 has no effect on the amplitude of the Calcium transient. Thus, unlike treatments that globally elevate cAMP in the cell, this selective displacement is expected to offer the additional benefit of improving cardiac performance without increasing SERCA activity and, consequently, ATP consumption. As evidenced by the response under dobutamine stress, it also preserves cardiac reserve, which instead is lowered by β-blockers.

We observed a marked reduction of fibrosis in mice treated with P33. Our data suggest that this is unlikely to be the primary mechanism of action of the disrupting peptide. P33 selectively displaced PDE4D9 from troponin I and increases TnI phosphorylation without altering global PKA or CaMKII signalling, phosphorylation of phospholamban or MyBPC, or activation of pro-hypertrophic transcriptional pathways. In contrast, the anti-fibrotic effect appears to be secondary, as P33 does not directly inhibit TGF-β-induced SMAD2/3 activation in cardiac fibroblasts and has no effect on SMAD2/3 phosphorylation in healthy hearts, while it normalises the elevated SMAD2/3 phosphorylation observed in TAC myocardium. While further studies will be required to establish the relative contribution to cardiac function improvement of reduced fibrosis, our data are consistent with a model in which local enhancement of TnI phosphorylation improves cardiomyocyte relaxation, reduces myocardial mechanical stress, and thereby attenuates activation of stress-responsive pro-fibrotic pathways.

The observation that the expression of PDE4D isoforms is elevated in human failing hearts is exciting and can explain the defective phosphorylation of TPNI widely reported in cardiac patients^48–52^. While PDE4 isoforms are relatively less abundant than PDE3 isoforms in human cardiac myocytes, we show that they significantly contribute to the regulation of cAMP at TPNI, and that displacement of PDE4D9 from TPNI with the disrupting peptide P33 results in local elevation of cAMP in human ventricular myocytes.

Currently, no effective treatment exists that selectively targets the mechanisms underpinning myofilament relaxation^73^. The high morbidity and mortality associated with diastolic dysfunction make this a major unmet medical need worldwide^4^. Our work indicates that targeting PDE4D9 displacement from the myofilament may provide a new, mechanism-based avenue for the treatment of heart disease. More broadly, it lays the foundation for the development of precision therapeutics in other organ systems where cAMP compartmentalisation orchestrates cellular function.

## Acknowledgements

M.Z acknowledges funding by the British Heart Foundation (RG/12/3/29423, RG/17/6/32944, PG/23/11321, PG/21/10611), the Oxford BHF Centre of Research Excellence by grant code (RE/08/004, RE/18/3/34214) and the Oxford NIHR Biomedical Research Centre. C.E.M. acknowledges funding by the Deutsche Forschungsgemeinschaft (ES 569/2-2 and ES 569/3-1), the Gertraud und Heinz-Rose Foundation and the German Centre for Cardiovascular Research (DZHK). The work-loop experiments were supported by InoCardia Ltd and Coventry University. C.L. acknowledges funding from British Heart Foundation grant (RG/18/12/34040).

## Author contributions

Y-C.C. and M.Z. designed the study, with M.Z. supervising the overall project and acquiring funding. Experimental work included imaging experiments performed by Y-C.C., W-H.H., E.V., M.G.O., A.K., N.C.S., J.H., M.F.; biochemical work performed by Y-C.C., R.Y., M.G.O.; MST experiments were performed by R.Y and J.M.E.; qPCR experiments were conducted by D.H.; calcium-tension curves were generated by D.W.A.D. and J.v.d.V.; in vitro work-loop experiments were conducted by J.H., A.L. and H.M.; peptide arrays were conducted by H.S, C.B. and G.B; work related to in vivo animal models was designed by C.L. and carried out by D.J.M, C.L., Y-C.C. and J.G.; work using hiPSC-derived cardiomyocytes was carried out by Y.P. and C.T.; histological analyses were conducted by W-H.H. and N.S.; the modelling was carried out by J.T.; work with human ventricular myocytes was carried out by D.R, E.G. and C.E.M; Y-C.C. and MZ wrote the manuscript.

## Additional information

Supplementary Information is available for this paper. Correspondence and requests for materials should be addressed to Manuela Zaccolo

## Supplementary information

To confirm the potential of disrupting the PDE4-Troponin I interaction in human setting, we simulated this in the newest advanced model of human excitation-contraction coupling T-World^1^. First, we developed a model of diastolic dysfunction from the baseline healthy myocyte model, using the following parameters: 180% I_NaL_ and tau of inactivation gate thL, 40% I_to,f_, 68% I_K1_, 80% I_NaK_, and 80% I_NaCa_, similar to^2^, 9-fold leak of Calcium from the sarcoplasmic reticulum^3^, and 80% SERCA pumping rate, reflecting its inhibition^4^. To represent enhanced PDE4-Troponin I interaction and thus facilitated Calcium binding, we reduced the ca50 of troponin-Calcium binding to 80% of the healthy myocyte value. Conversely, to represent treatment which disrupts this interaction, we increased the ca50 value of troponin-Calcium binding to 120% of the healthy myocyte value.

The resulting model of diastolic dysfunction shows elevated resting tension at 60 beats per minute compared to a healthy cell (Suppl Figure 24a), which increases further as the pacing rate increases, consistent with human data on diastolic dysfunction^5^. Using a different indicator of diastolic dysfunction, the time from peak contraction to 90% relaxation, the model of diastolic dysfunction likewise shows a clear impairment of relaxation (Suppl Figure 24b). Importantly, both indices of diastolic dysfunction are normalised by the emulated disruption of PDE4-Troponin interaction (Suppl Figure 24a-b).

### Extended Materials and Methods

#### Microscale thermophoresis (MST) assay

PDE4D9 and TPNI constructs in pET-24a were transformed into BL21(DE3) Rosetta2 cells and expressed in LB medium supplemented with kanamycin (50 µg/mL) and chloramphenicol (34 µg/mL). Cultures were grown at 37 °C to an OD600 of 0.4–0.5, cooled to 18 °C, induced with 0.5 mM IPTG, and incubated overnight. Cells were harvested, lysed by sonication in Tris/NaCl buffer containing imidazole, TCEP, detergent, and protease inhibitors, and clarified lysates were purified using Ni-NTA affinity chromatography. Eluted proteins were treated with TEV protease, dialysed overnight, and subjected to a second Ni-NTA purification followed by size-exclusion chromatography on a Superdex S200 16/60 column. Purified fractions were pooled and concentrated by ultrafiltration. Proteins were labelled with Red NHS reactive dye (NanoTemper Technologies) according to the manufacturer’s instructions. Briefly, proteins were adjusted to 10 µM in labelling buffer and incubated with NHS dye for 30 min at room temperature in the dark, followed by removal of unreacted dye using the supplied filtration columns. MST measurements were performed on a MONOLITH NT.115 instrument (NanoTemper Technologies) in binding buffer using ∼7.75 nM labelled TPNI and serial dilutions of PDE4D9 (∼14 µM starting concentration). Measurements were repeated using independent protein preparations. Binding curves were generated by plotting protein concentration against fraction bound and analysed in Prism 9 (GraphPad Software).

#### SUMO affinity enrichment and immunoblotting

ARVMs were treated with vehicle or isoproterenol (ISO, 50 nM) for 10 min and lysed in ice-cold lysis buffer supplemented with 20 mmol/L N-ethylmaleimide (NEM, ThermoFisher, 040526.03) and protease inhibitors. SUMOylated proteins were enriched using SUMO-Tag Trap Agarose (ChromoTek, suta) according to the manufacturer’s instructions. Following extensive washing, bound proteins were eluted in SDS sample buffer and subjected to SDS-PAGE and immunoblot analysis. Whole-cell lysates and SUMO-enriched fractions were probed with antibodies against PDE4D9, SUMO-1(Proteintech, 67559-1-Ig, 1:1000), and TPNI.

#### TGF-β1 stimulation of neonatal rat cardiac fibroblasts

Neonatal rat cardiac fibroblasts were obtained as a by-product of the NRVM isolation procedure and used at passages 0–2. Cells were pre-treated with scrambled control peptide (SC, 10 μM) or P33 peptide (10 μM) for 2 h prior to stimulation with recombinant TGF-β1 (Life Technoloiges, 100-21, 5 ng/mL) for 30 min. Vehicle-treated cells (DMSO) served as controls in the presence of TGF-β1, whereas mock-treated cells received PBS without TGF-β1 stimulation. Phospho-SMAD2/3 (Cell Signaling, 8828, 1:1000) and total SMAD2/3 (Abcam, ab305325, 1:1000) levels were subsequently analysed by western blotting as described above.

#### CREB phosphorylation analysis

NRVMs were pre-treated with scrambled control peptide (SC, 10 μM) or P33 peptide (10 μM) prior to stimulation with isoproterenol (ISO, 10 μM) or a saturating stimulus consisting of forskolin (25 μM) and IBMX (100 μM) for 10 min. Vehicle-treated cells (DMSO) served as controls. Cells were subsequently lysed and phospho-CREB (Cell Signaling, 9198, 1:1000) and total CREB (Cell Signaling, 9104, 1:1000) levels were analysed by western blotting as described above. Phospho-CREB signals were normalised to total CREB levels and expressed relative to unstimulated basal controls.

#### NFAT phosphorylation analysis

NRVMs were transfected with an NFAT-GFP expression plasmid using Transfectin reagent (1703351, Bio-Rad) and cultured for 36–48 h prior to experimentation. Cells were pre-treated with scrambled control peptide (SC, 10 μM) or P33 peptide (10 μM) and subsequently stimulated with isoproterenol (ISO, 10 μM) or a saturating stimulus consisting of forskolin (25 μM) and IBMX (100 μM) for 10 min. Cells were lysed and GFP-tagged proteins were enriched using GFP-Trap Agarose (ChromoTek, gta-100) according to the manufacturer’s instructions. Immunoprecipitates were analysed by western blotting as described in the main Methods section and probed with antibodies against phospho-PKA substrates (Cell Signaling, 9624, 1:1000) and GFP (Abcam, ab183735, 1:1000). Phosphorylation signals were normalised to immunoprecipitated NFAT-GFP and expressed relative to unstimulated basal controls.

#### Analysis of PKA and CaMKII signalling proteins

Left ventricular (LV) tissue was collected from healthy mice 3 weeks after scAAV9-SC or scAAV9-P33 injection, or from mice subjected to sham or transverse aortic constriction (TAC) surgery and treated as illustrated in Fig. 5a. Protein abundance of PKA RIα (Proteintech, 20358-1-AP, 1:1000), PKA RIIα (Santa Cruz, SC-137220, 1:1000), PKA catalytic subunit (Santa Cruz, SC-28315, 1:500), and CaMKII (BD Biosciences, 611293, 1:1000), together with phospholamban (PLB) phosphorylation at Ser16 (Badrilla, A010-12AP, 1:4000) and Thr17 (Antibodies.com, A93027, 1:1000), were assessed by western blotting using the antibodies indicated above. Protein expression and phosphorylation levels were quantified by densitometry and normalised as described in the main Methods section.

#### PDE activity assay

HEK293 cells were transiently transfected with PDE4D9 or left non-transfected as controls. 36-48 hours after transfection, cells were treated with vehicle (DMSO), scrambled control peptide (SC, 10 μM), or P33 peptide (10 μM) for 2 h. Cells were subsequently stimulated with forskolin (10 μM) for 10 min at 37 °C and harvested for phosphodiesterase activity measurements. PDE activity was quantified using a commercially available phosphodiesterase activity assay kit (AkrivisBio/Enzo; Cat. No. MA-0160) according to the manufacturer’s instructions.

#### Myofilament calcium sensitivity measurements

Cardiomyocytes were enzymatically isolated from 5 rat hearts (Wistar; 3 males, 2 females) as described above, and incubated in Tyrode buffer (0,5 ml, 0.1 mM Ca^2+^) for 2 hrs with 10 µM P33 or 10 µM SC at 37°C. Subsequently, cardiomyocytes were incubated with 10 nM isoprenaline for 0, 1, 5 or 10 minutes, and samples were collected for analyses of cTnI phosphorylation (3 experiments), and assessment of myofilament Ca^2+^-sensitivity (2 experiments). Blots were stained with total protein stain to correct for minor loading differences, and TPNI phosphorylation signal was divided by total protein signal in each sample.

For force measurements cardiomyocytes were chemically permeabilized by incubation for 5 min in relaxing solution containing 0.5% (v/v) Triton X-100 and glued between a force transducer and a piezoelectric motor, as described previously^6^.

## Extended Data Figures and Tables

**Extended Data Fig. 1.**
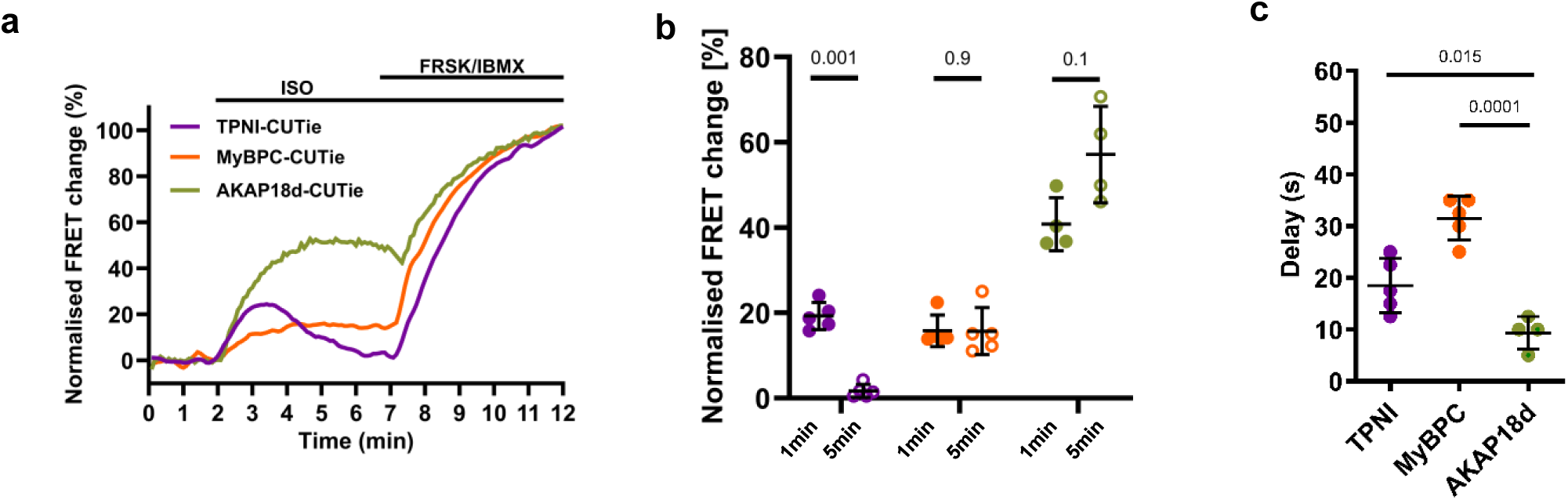
Local cAMP regulation in neonatal cardiomyocytes. **(a)** Representative kinetics of FRET responses recorded in NRVM expressing the indicated targeted sensors and stimulated with ISO (1 nM), followed by saturating stimulation with forskolin (25 μM) and IBMX (100 μM). (**b**) Summary of cAMP responses measured at 1and 5 min following ISO stimulation from experiments shown in (a). Statistical significance was assessed using paired t-tests. (**c**) Average time required to achieve 5% of the maximal FRET response following ISO stimulation. Statistical significance was assessed using paired t-tests and Welch’s t-tests with Benjamini–Hochberg correction. Each data point represents an independent biological replicate. Data are presented as mean ± SD. n ≥ 4 biological replicates per group.

**Extended Data Fig. 2.**
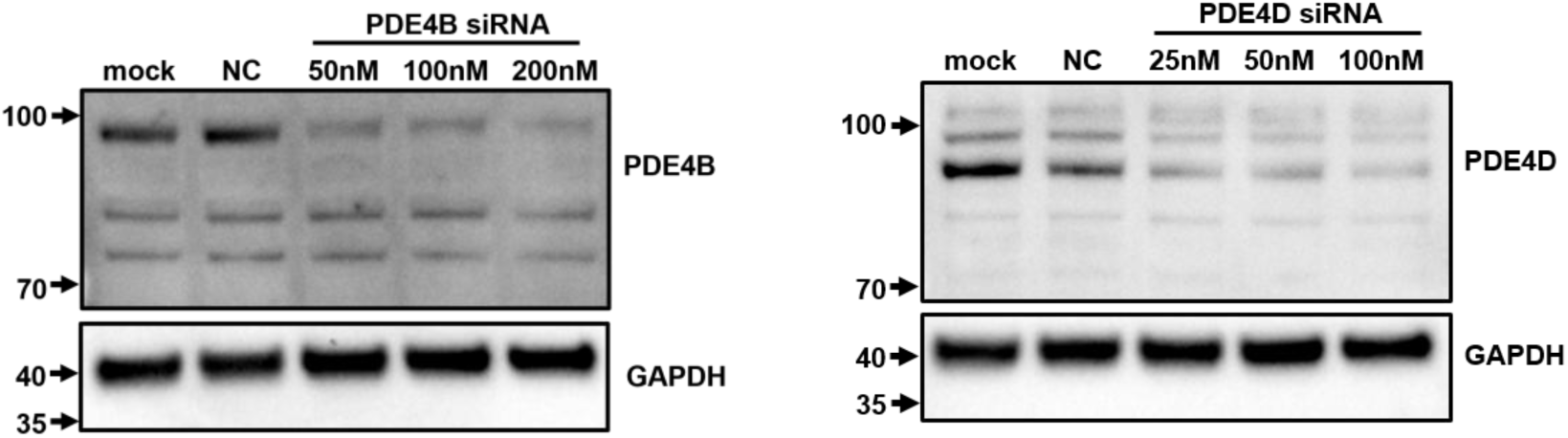
siRNA knockdown attenuates expressions of endogenous PDE4B and 4D in NRVMs. Western blot analysis of cell lysates from NRVM treated with different concentrations of PDE4D and PDE4B siRNA, as indicated, or with the non-targeting siRNA as negative control (NC). Mock indicates untreated cells.

**Extended Data Fig. 3:**
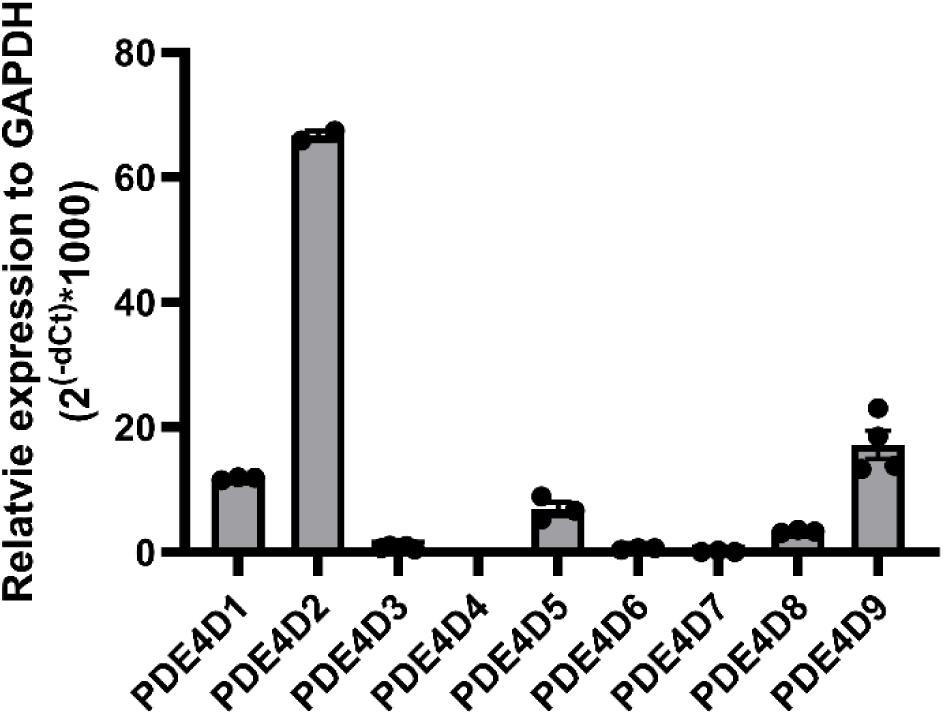
PDE4D long form expression in NRVMs. Quantitative PCR analysis showing the relative mRNA expression for different PDE4D isoforms in neonatal rat ventricular myocytes. n ≥ 3 biological replicates.

**Extended Data Fig. 4:**
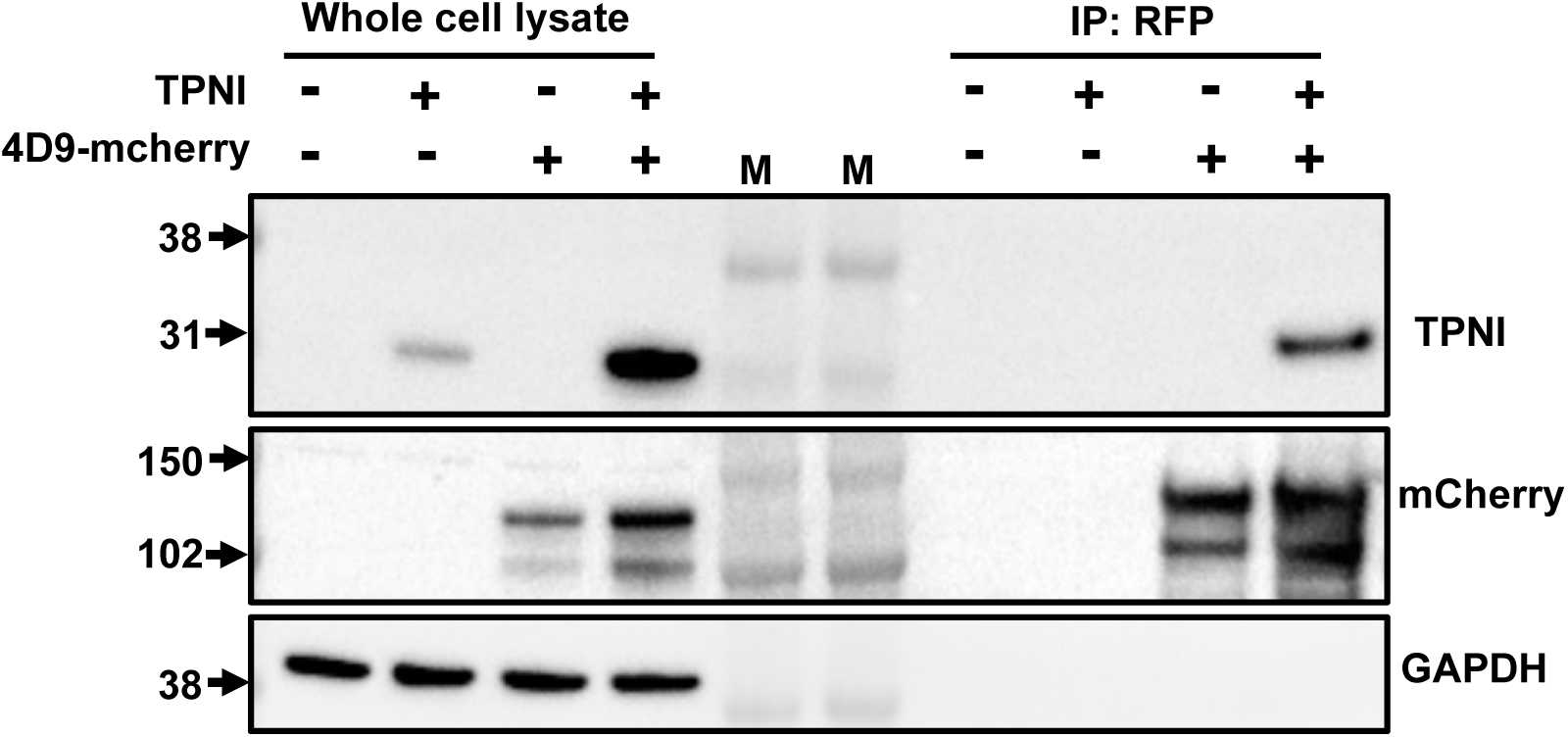
PDE4D9 interacts with TPNI in HEK293T cells. Representative western blots of whole-cell lysates (WCL) and RFP-Trap pull-down fractions from HEK293T cells expressing TPNI and/or PDE4D9-mCherry. Membranes were probed with antibodies against TPNI, mCherry, and GAPDH. M = molecular weight markers. n ≥ 3 biological replicates.

**Extended Data Fig. 5:**
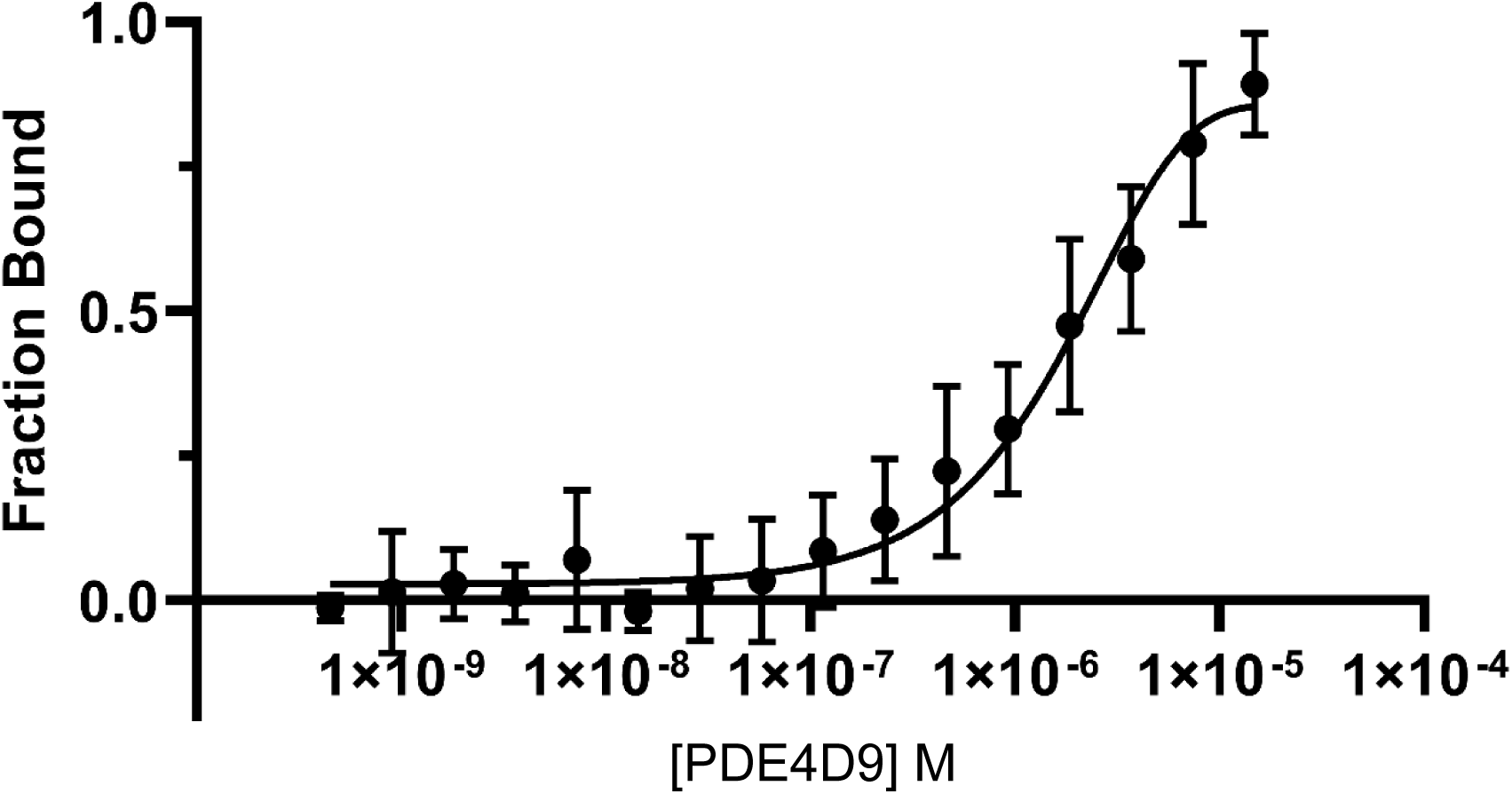
Analysis of direct interaction between TPNI and PDE4D9 by microscale thermophoresis (MST). MST was used to further confirm direct interaction between PDE4D9 and TPNI. TPNI was fluorescently labeled with Red-NHS dye. Approximately 7.75 nM fluorescently labeled TPNI was mixed with serially diluted PDE4D9 (starting at ∼14 µM). MST, measurements reveal that PDE4D9 interacts with purified TPNI with a binding affinity (KD) of approximately 3.03 ± 0.73 µM (mean ± SEM, n = 5).

**Extended Data Fig. 6:**
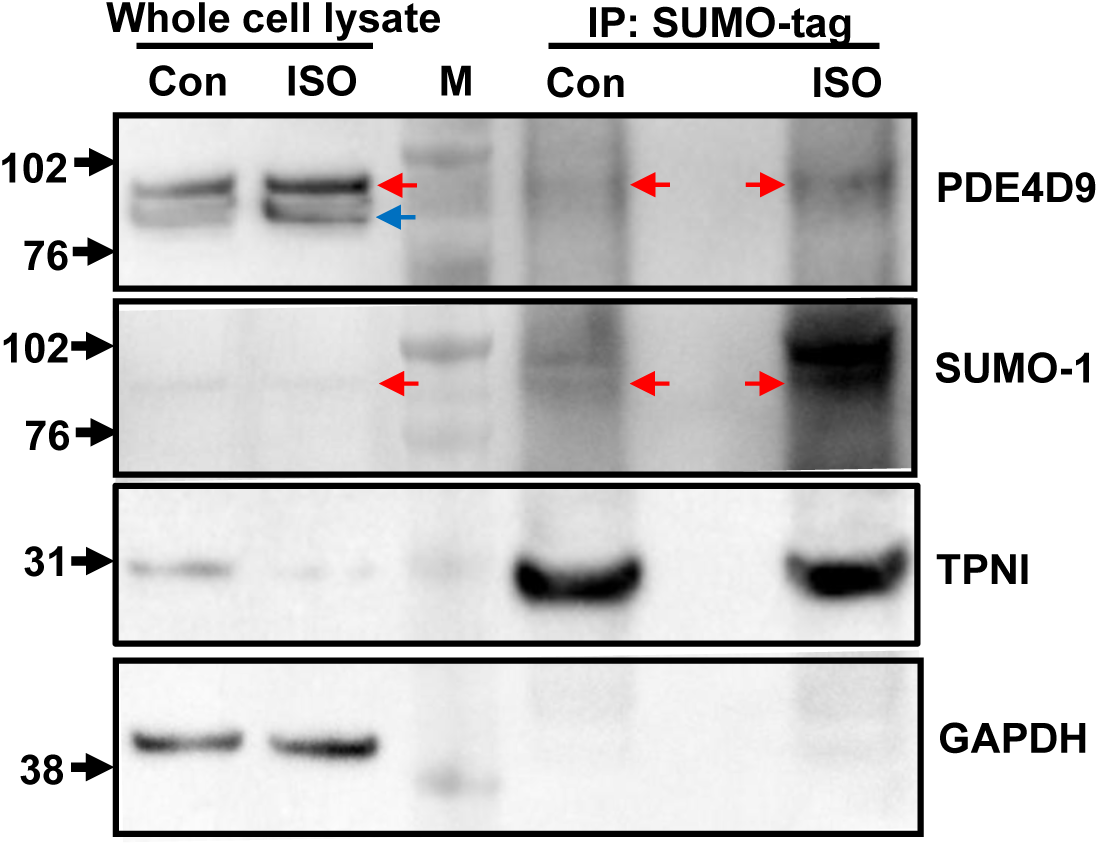
ISO stimulation enriches SUMOylated PDE4D9 associated with TPNI. Primary adult rat ventricular cardiomyocytes (ARVMs) were treated with vehicle or isoproterenol (ISO, 50 nM) for 10 min and lysed in the presence of 20 mM N-ethylmaleimide (NEM) to preserve protein SUMOylation. SUMOylated proteins were enriched using SUMO-Tag Trap Agarose and analyzed by immunoblotting. Whole-cell lysates (WCL) and SUMO-enriched fractions were probed with antibodies against PDE4D9, SUMO-1, and TPNI. PDE4D9 immunoblotting of WCL samples revealed two bands corresponding to lower and higher molecular weight species. In contrast, the IP fraction was enriched for the upper shifted band (red arrow), whereas the lower band (blue arrow) was absent. Immunoblotting with anti-SUMO-1 antibody detected a signal corresponding to the upper shifted species, consistent with this band representing SUMOylated PDE4D9 associated with the troponin complex. ISO treatment further increased enrichment of the SUMOylated species in the IP fraction. n = 2 biological replicates.

**Extended Data Fig. 7.**
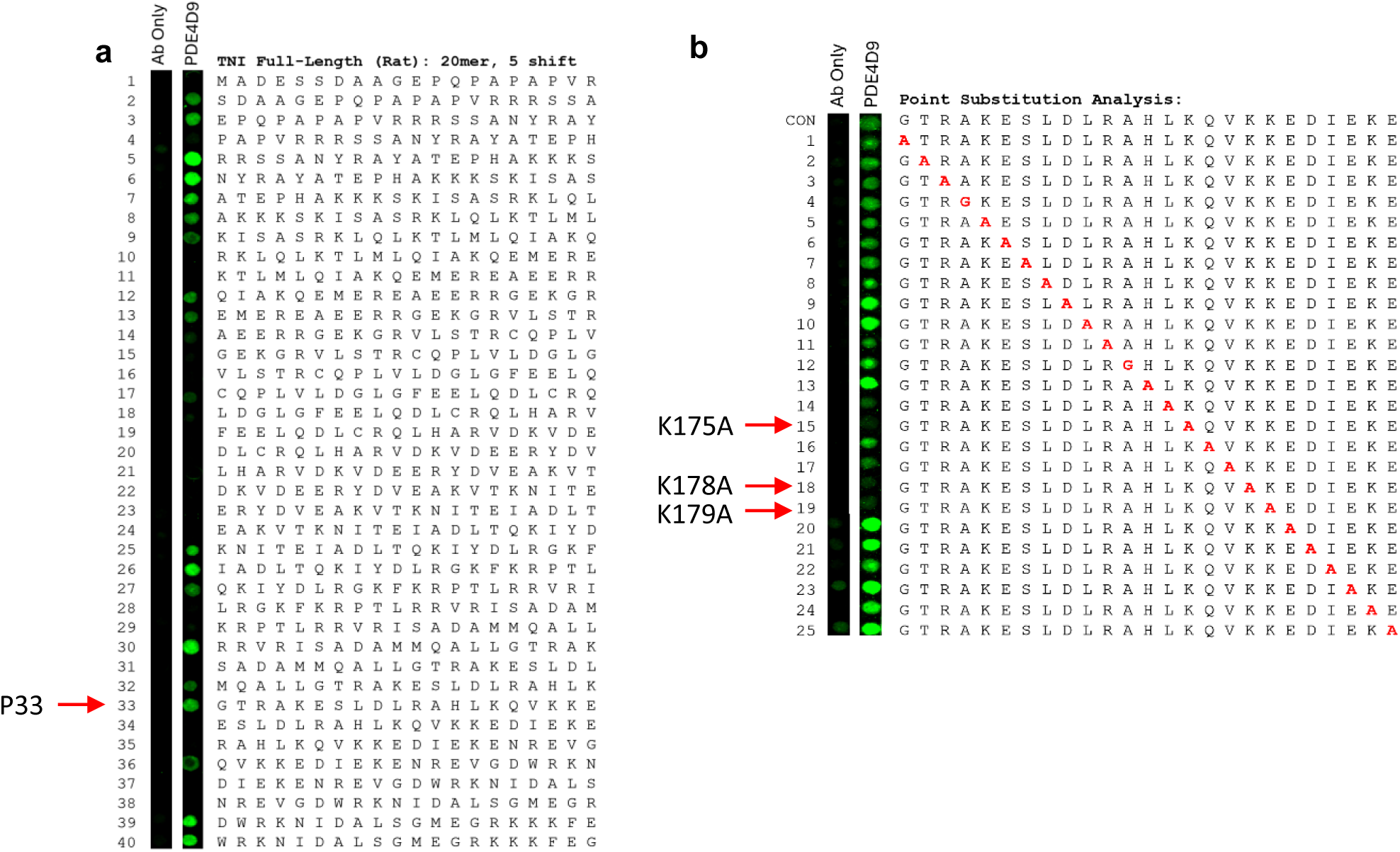
PDE4D9 interacts with TPNI. (**a**) Peptide array of the full-length sequence of rat cardiac TPNI probed with GST-PDE4D9. Red arrow indicates peptide 33, selected for further analysis (**b**) Alanine scanning of the TPNI peptide shown in (a) at position 33 with sequence aa161-185. Red arrows indicate peptides with lysin-to-alanine mutation at position 175, 178 and 179, as indicated.

**Extended Data Fig. 8.**
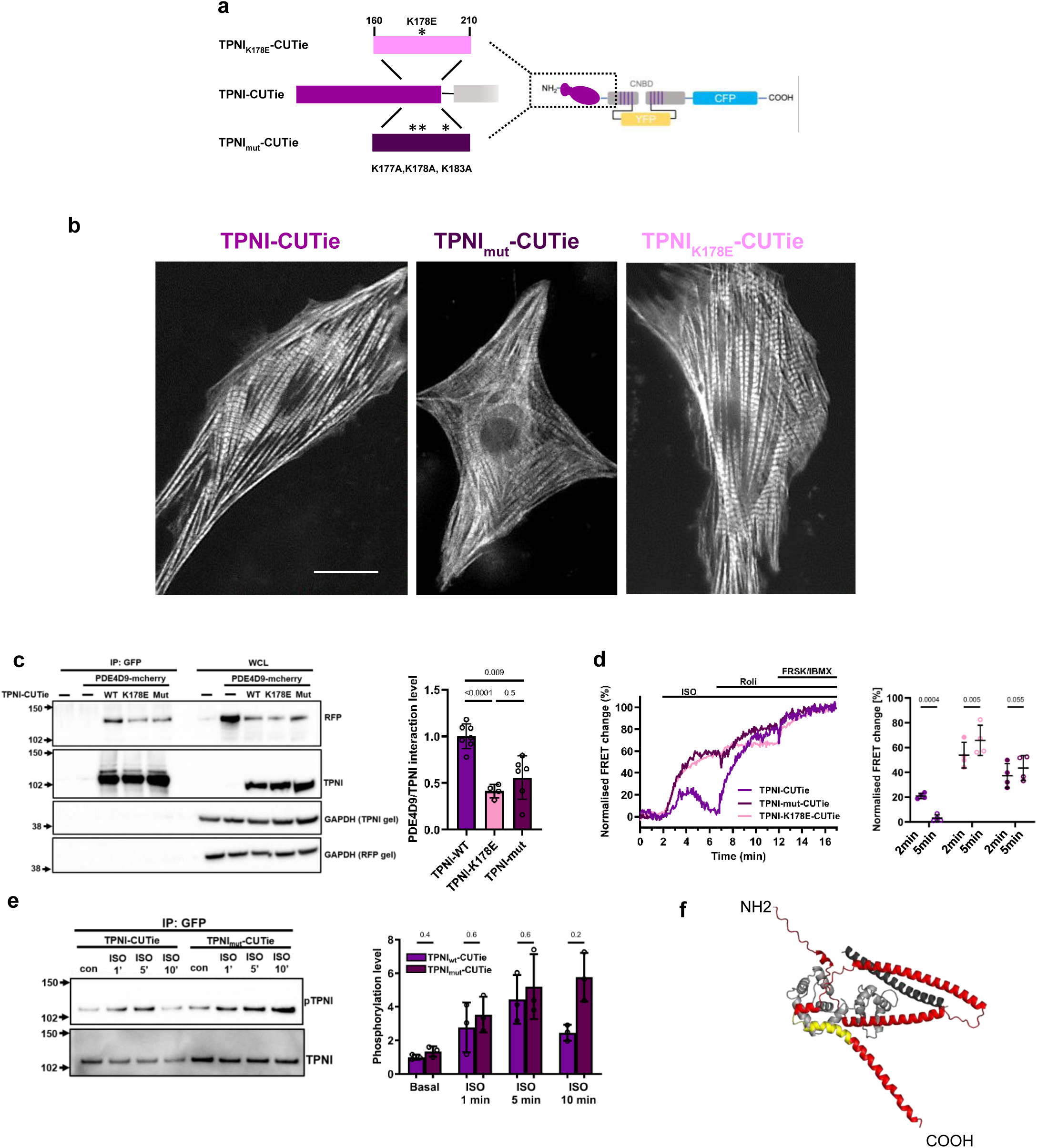
Key amino acid residues involved in the PDE4D9/TPNI interaction. (**a**) Schematic representation showing the mutation sites and (**b**) subcellular localisation in NRVM of TPNImut- and TPNIK178E-CUTie compared to wild type TPNI-CUTie. Scale bar = 10 μm, (**c**) Representative western blot and quantification of GFP pull-down from lysates of HEK293T cells expressing TPNI-, TPNIK178E-, or TPNImut-CUTie and PDE4D9-mcherry. Welch’s ANOVA followed by Dunnett’s T3 multiple-comparison test. WCL = whole cell lysate. (**d**) Representative kinetics and summary of FRET responses recorded at 2 and 5 min following stimulation with ISO (1 nM) in NRVM expressing TPNI-CUTie, TPNImut-CUTie, or TPNIK178E-CUTie. Paired t-tests. (**e**) Representative western blot and quantification of GFP pull-down and phosphorylation levels of the TPNI moiety in TPNI-CUTie and TPNImut-CUTie expressed in NRVM and collected at the indicated time points following stimulation with ISO (1 nM). Welch’s t-tests with Benjamini–Hochberg correction. Each data point represents an independent biological replicate. Data are presented as mean ± SD. n ≥ 3 biological replicates per group. (**f**) Structural model of TPNI (red) in complex with TPNC (light grey) and TPNT (dark grey) generated using AlphaFold. The sequence corresponding to peptide P33 within TPNI is highlighted in yellow.

**Extended Data Figure 9:**
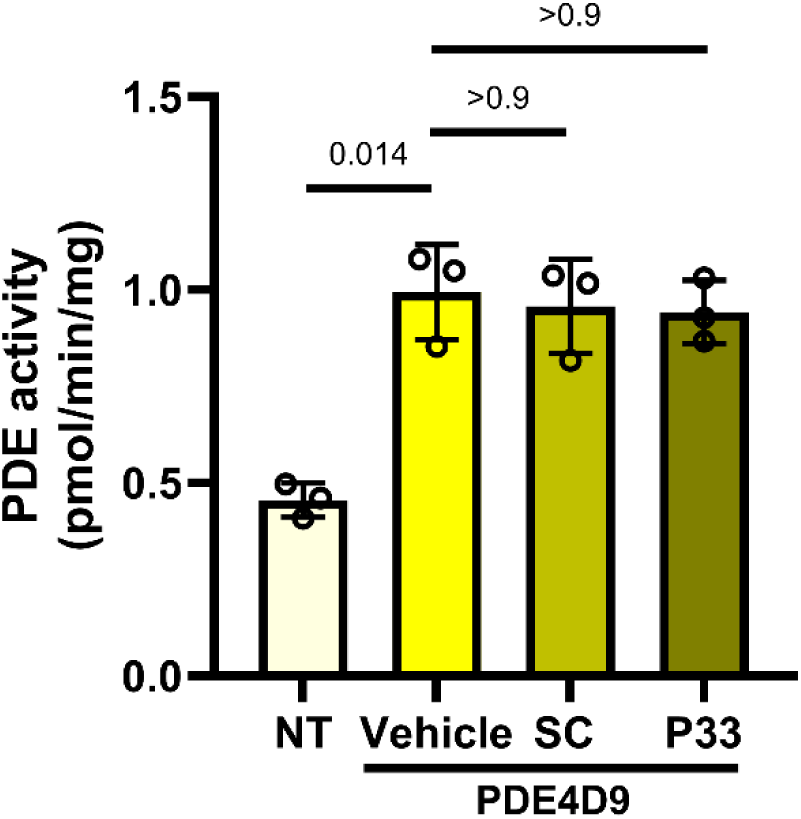
P33 does not directly inhibit PDE4D9 catalytic activity. PDE activity measured in HEK293 cells transfected with PDE4D9 and treated with vehicle (DMSO), scrambled control peptide (SC, 10 μM), or P33 (10 μM) for 2 h prior to stimulation with forskolin (10 μM) for 10 min. Non-transfected (NT) cells served as controls. PDE activity was quantified using a phosphodiesterase activity assay. Statistical significance was assessed using Welch’s ANOVA followed by Dunnett’s T3 multiple-comparison test. Each data point represents an independent biological replicate. Data are presented as mean ± SD. n = 3 independent experiments per group.

**Extended Data Figure 10.**
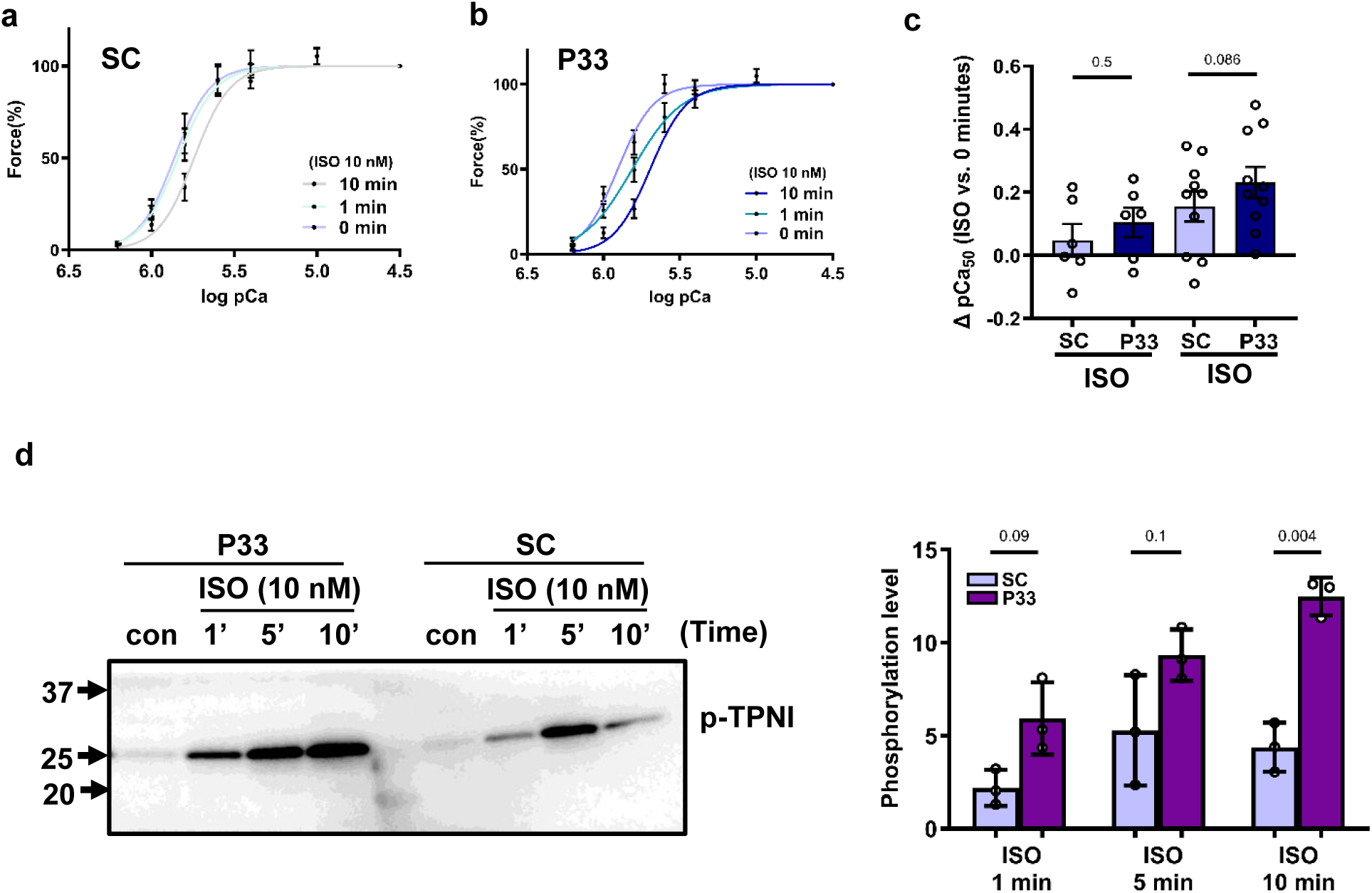
P33 treatment enhances myofilament Ca²⁺ sensitivity. (**a, b**) Force–pCa relationships of skinned cardiomyocytes treated with scrambled control peptide (SC) or P33 and subsequently stimulated with isoprenaline (ISO, 10 nM) for 0, 1, or 10 min. Force was normalised to maximal force and plotted against pCa. (**c**) Change in pCa₅₀ relative to the corresponding 0 min condition (ΔpCa₅₀). Paired t-tests. Individual symbols represent single cardiomyocytes and bars indicate mean ± SEM. Data were obtained from two independent isolations (N = 2) and 10 cardiomyocytes (n = 10). (**d**) Representative immunoblot and quantification of cardiac troponin I phosphorylation in SC- and P33-treated cardiomyocytes following stimulation with ISO (10 nM) for the indicated times. Welch’s t-tests with Benjamini–Hochberg correction. Each data point represents an independent biological replicate (n = 3). Data are presented as mean ± SD. pCa₅₀, negative logarithm of the free Ca²⁺ concentration producing 50% maximal force.

**Extended Data Fig. 11.**
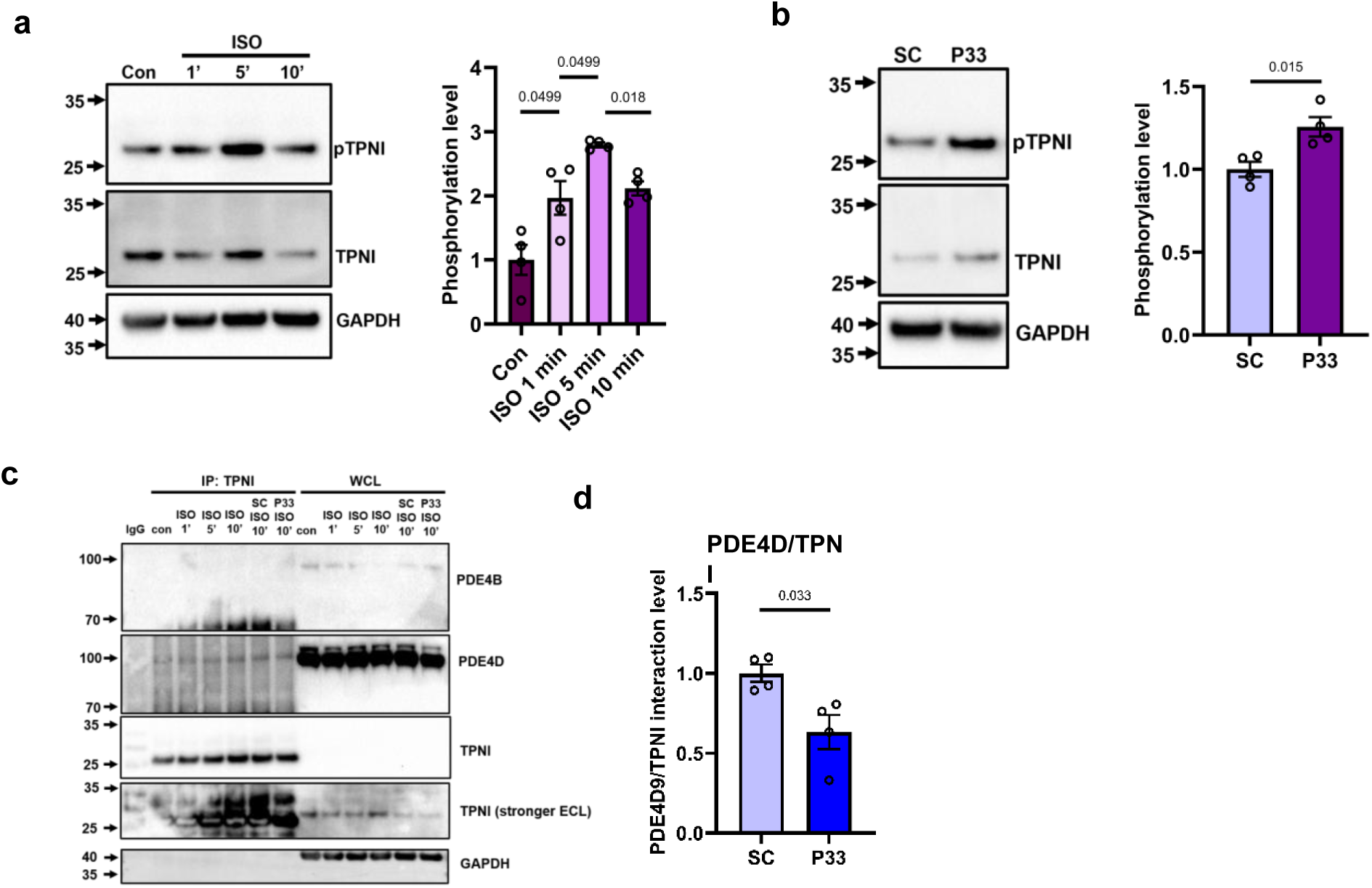
Regulation of TPNI phosphorylation by PDE4D9 in hiPS-CM. (**a**) Representative western blot and quantification of lysates from hiPSC-CMs treated with ISO (1 nM) for 1, 5, or 10 min showing the time course of TPNI phosphorylation. Welch’s t-tests with Benjamini–Hochberg correction. (**b**) Representative western blot and quantification of lysates from hiPSC-CMs pre-treated with P33 peptide or scrambled control peptide (SC, 10 μM) followed by stimulation with ISO (1 nM) for 5 min. (**c**) Representative western blot and (**d**) quantification of co-immunoprecipitation experiments using lysates from hiPSC-CMs pre-treated with P33 peptide or scrambled control peptide (SC, 10 μM) and stimulated with ISO (1 nM) for 10 min. Welch’s t-tests. Each data point represents an independent biological replicate. WCL = whole cell lysate. Data are presented as mean ± SD. N = 4 biological replicates per group.

**Extended Data Fig. 12.**
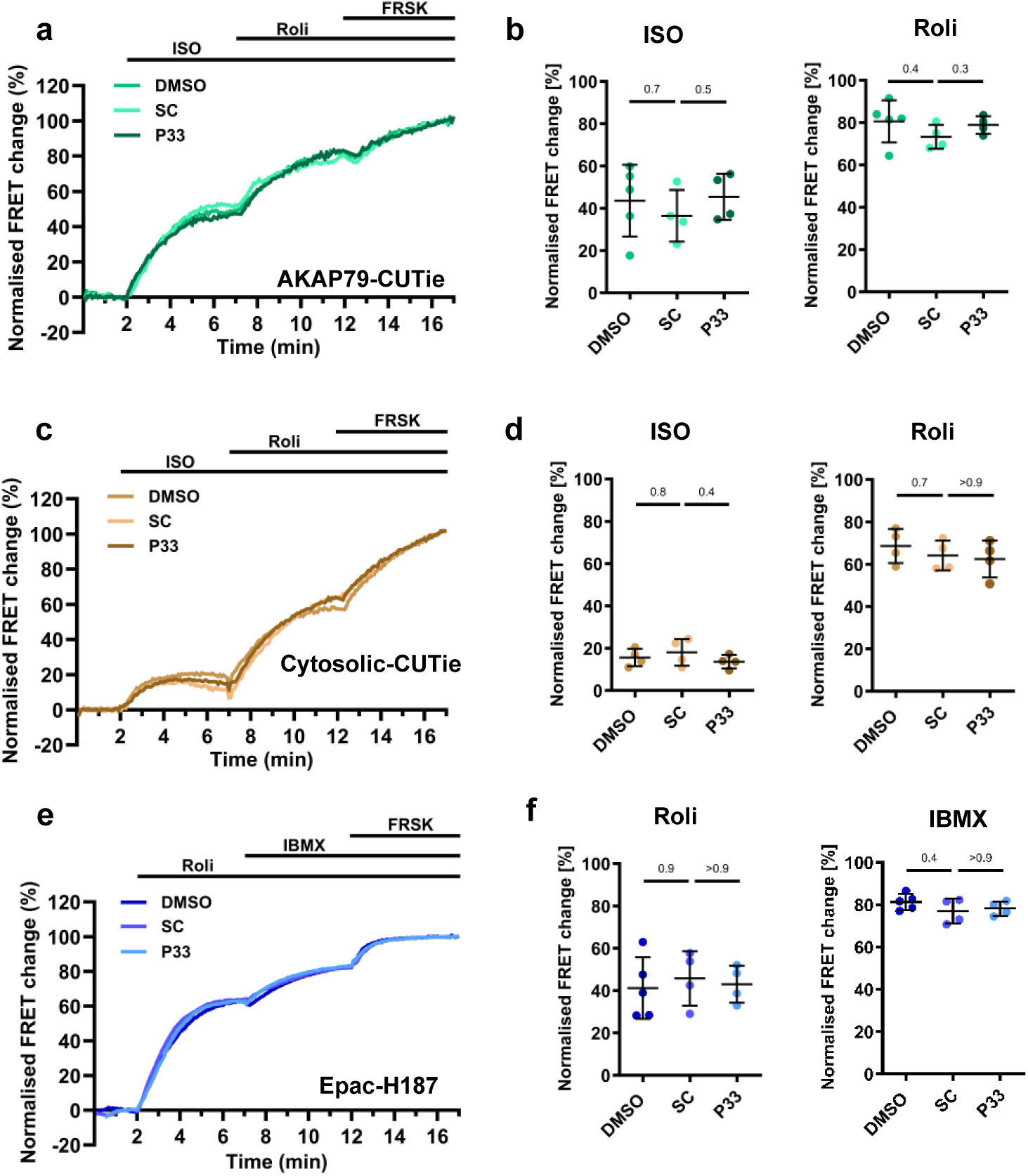
Effect of P33 on overall PDE activity or on the cAMP response in the bulk cytosol and at the plasmalemma. (**a**) Representative kinetics of FRET change recorded in NRVM expressing the plasmalemma-targeted cAMP reporter AKAP79-CUTie and pre-treated with P33, SC peptides (10 μM) or DMSO as control, upon application of ISO (1 nM) followed by Rolipram (10 μM) and saturating treatment (**b**) Summary of response amplitude in experiments as described in (a). (**c**) Representative kinetics of FRET change recorded in NRVM pre-treated with P33 (10 μM), SC peptides (10 μM) or DMSO as control and expressing the cytosolic cAMP reporter CUTie on application of ISO (1 nM) followed by rolipram (10 μM) and saturating treatment. (**d**) Summary of data collected as in (c). (**e**) Representative kinetics of FRET change recorded in NRVM pre-treated with P33 (10 μM), SC peptides (10 μM) or DMSO. Cells express the cytosolic cAMP reporter Epac-H187 and are treated with Rolipram (10 μM) followed by IBMX (100 μM) and saturating treatment. (**f**) Summary of data collected as in (e). In all cases, data points are values recorded at 5 min after treatment. Welch’s ANOVA followed by Dunnett’s T3 multiple-comparison test. Each data point represents an independent biological replicate. Data are presented as mean ± SD. n ≥ 4 biological replicates per group.

**Extended Data Fig. 13.**
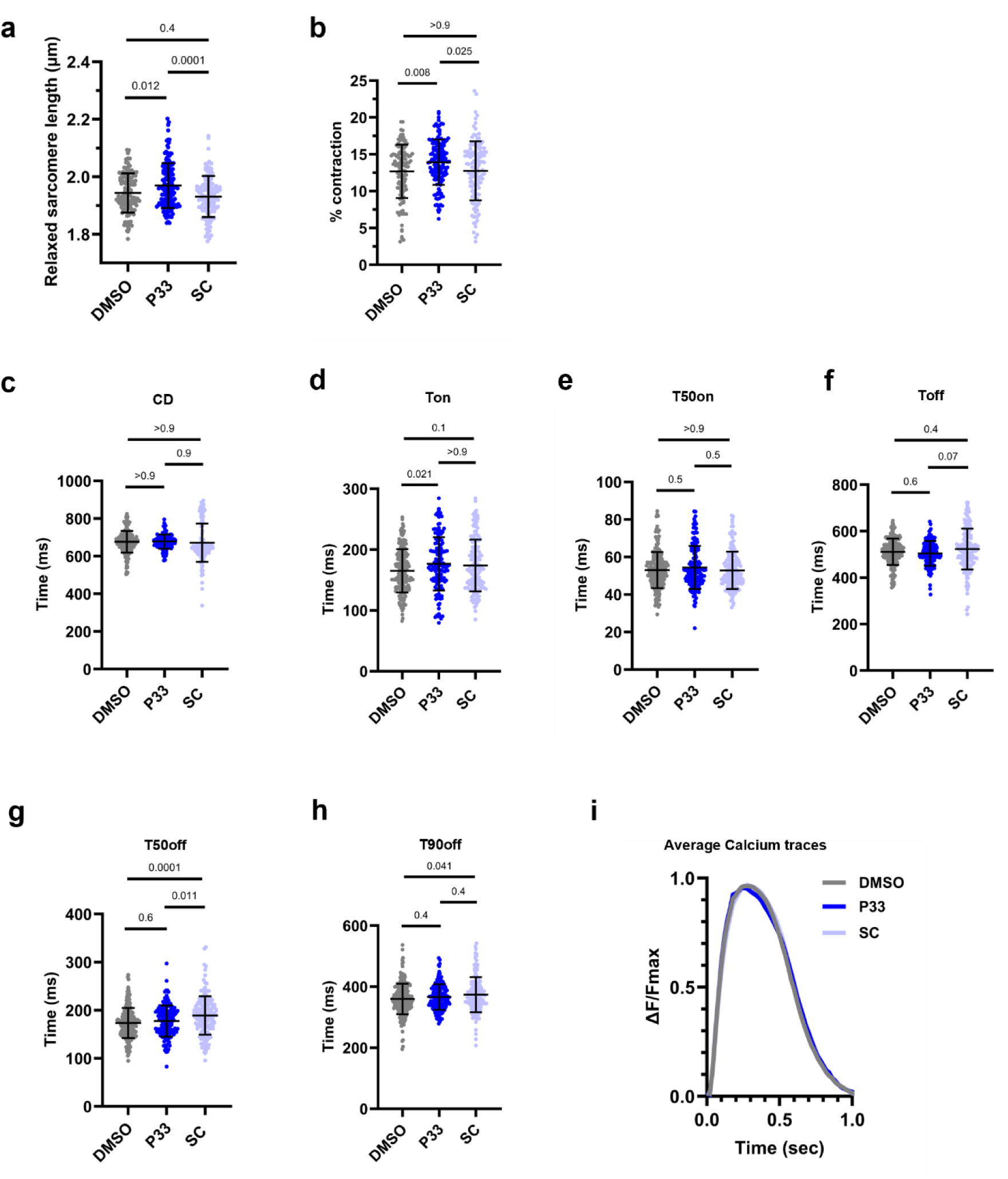
hiPSC-derived cardiomyocytes treated for 2h with P33 (10 μM), SC peptide (10 μM), or DMSO were analysed with SarcTrack to measure (**a**) contractility and (**b**) cardiomyocyte shortening. CalTrack analysis of the same cells shows (**c-h**) Calcium transient parameters and (**i**) average normalised Calcium transient waveforms. Statistical significance was assessed using Welch’s ANOVA followed by Dunnett’s T3 multiple-comparison test. n = 4 differentiations. Data are presented as mean ± SD. CD = total Calcium duration; Ton = time to peak; T50on = time to 50% peak; Toff = time of decay; T50off = time to 50% of the baseline; T90off = time to 90% of baseline.

**Extended Data Fig. 14.**
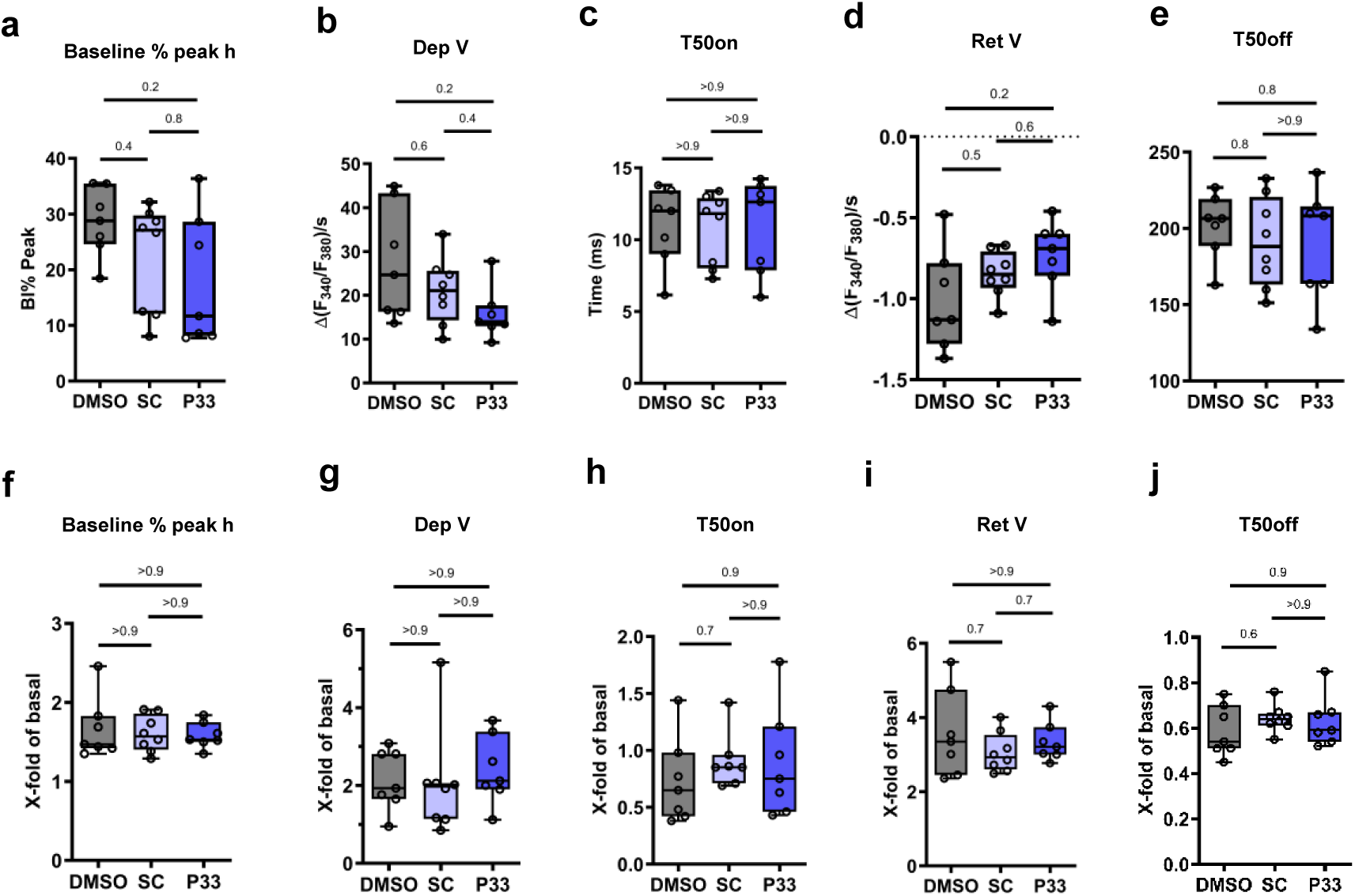
Effect of P33 on the amplitude and duration of the Calcium transient in ARVM. Analysis of Calcium transient parameters in unstimulated condition in the presence of DMSO, SC peptide (10 μM), or P33 peptide (10 μM). Electrically evoked Calcium transients were measured in monolayers under control conditions for 2 min. Monotonic transient analysis was performed to evaluate transient parameters for the peak (**a**), departure (**b, c**), and return phase (**d, e**) of the Calcium transients. (f-j) Electrically evoked Calcium transients were measured in monolayers under basal conditions for 2 min, and after ISO (5 nM) application for further 2 min in the presence of SC peptide (10 μM) or P33 peptide (10 μM). Following monotonic transient analysis, the ISO effect was calculated as ratio of mean transient parameter under ISO exposure divided by mean transient parameter under basal conditions for each monolayer. Therefore, data are shown as x-fold of basal. ISO effect is shown for parameters describing peak (f), departure (g, h), and return phase (i, j) of Calcium transients. Welch’s ANOVA followed by Dunnett’s T3 multiple-comparison test. Each data point represents an independent biological replicate. Data are shown as min-to-max whiskers box plots. n ≥ 3 biological replicates per group. Bl% peak = baseline as a percentage of peak height, the percent change during the transient; Dep V = departure velocity, the maximal rate of change during Calcium release phase of transient; Ret V = return velocity, the maximal rate of the return phase of the transient; T50on = time to 50% peak (Calcium elevation); T50off = time to 50% of the baseline (Calcium reuptake).

**Extended Data Fig. 15.**
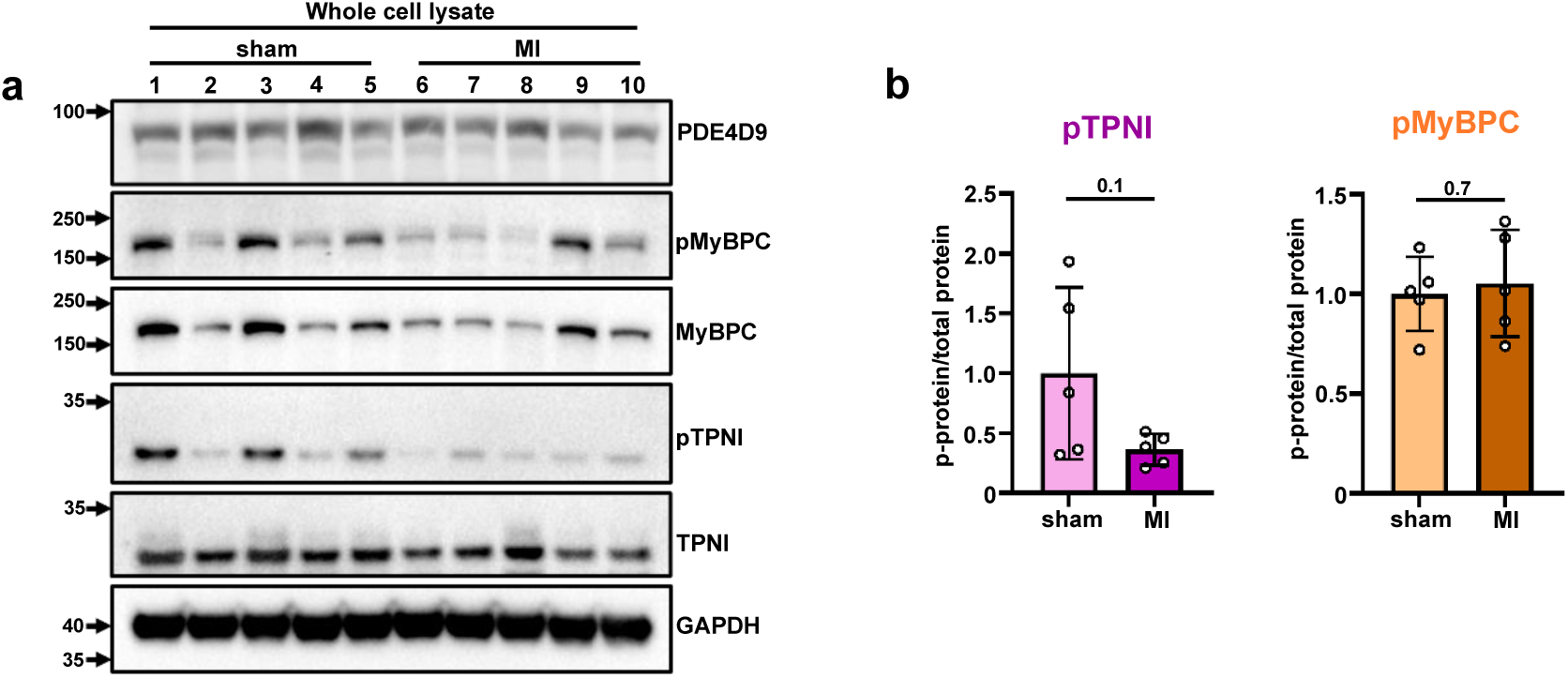
(**a**) Representative western blot and (**b**) quantification of TPNI and MyBPC phosphorylation in whole cell lysates (WCL) from sham or myocardial infarcted (MI) mouse hearts. Welch’s t-tests. Lane numbers and data points represent individual animals. n = 5 animals per group. Data are presented as mean ± SD.

**Extended Data Fig. 16.**
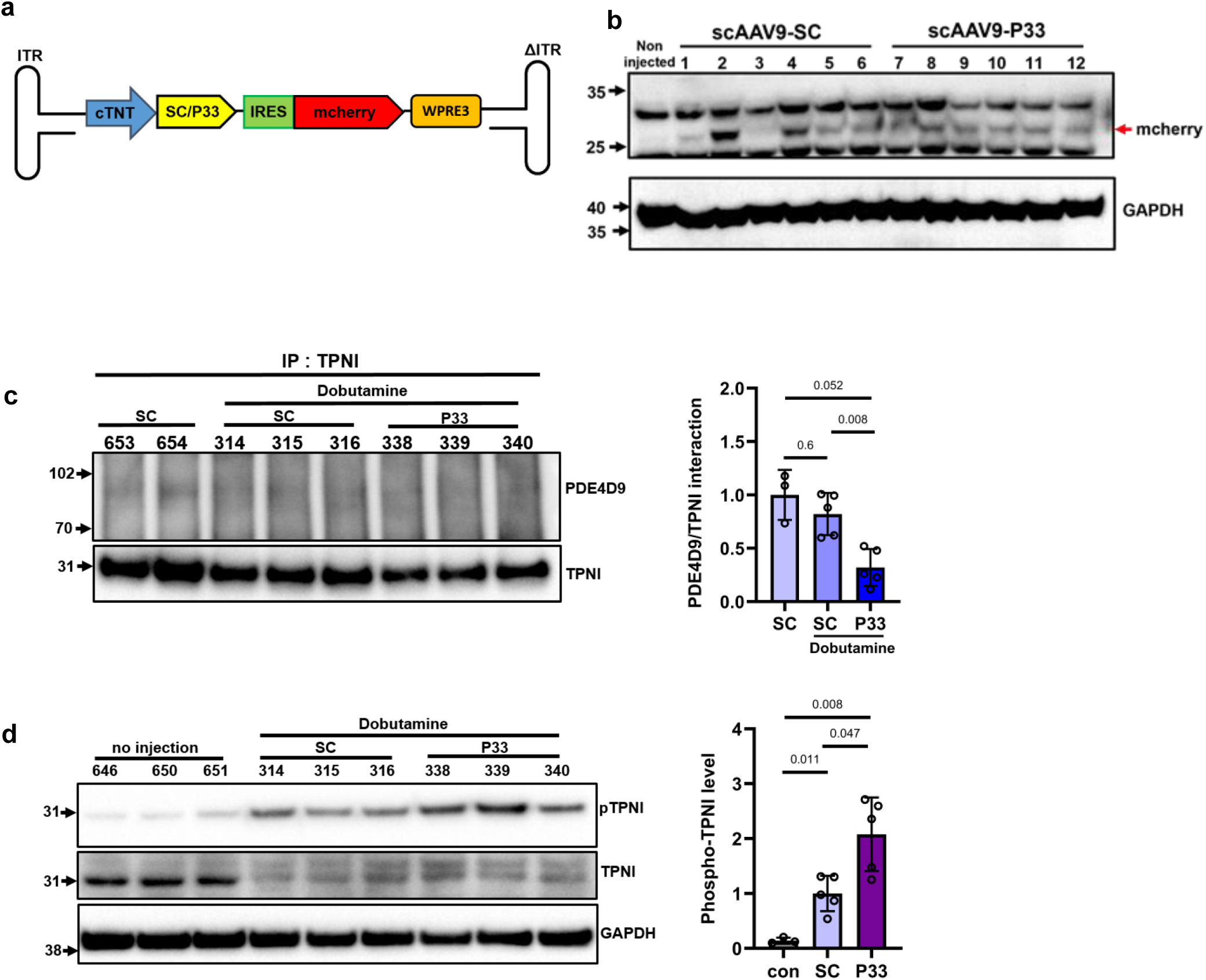
scAAV9-P33 injection attenuates PDE4D9/TPNI interaction and enhances TPNI phosphorylation. (**a**) Schematic of the self-complementary AAV9 (scAAV9) cassette carrying SC or P33 coding sequences under the control of the cardiac specific Troponin T promoter (cTNT) and bicistronic (IRES) mCherry expression system with WPRE3 fragment at the 3’-untranslated region to enhance expression efficiency. (**b**) Western blot of lysates from mouse left ventricular (LV) tissues harvested 3 weeks after scAAV9-SC or scAAV9-P33 injections, probed with mCherry antibody. (**c**) Representative western blot and quantifications in samples as in (b) showing the effect of expressing SC or P33 on the PDE4D9/TPNI interaction. (**d**) Whole cell lysates from the same samples as in (b) showing the effect of SC or P33 on the TPNI phosphorylation level. Welch’s ANOVA followed by Dunnett’s T3 multiple-comparison test. Lane numbers and data points represent individual mice. n ≥ 3 mice per group. Data are presented as mean ± SD.

**Extended Data Fig. 17:**
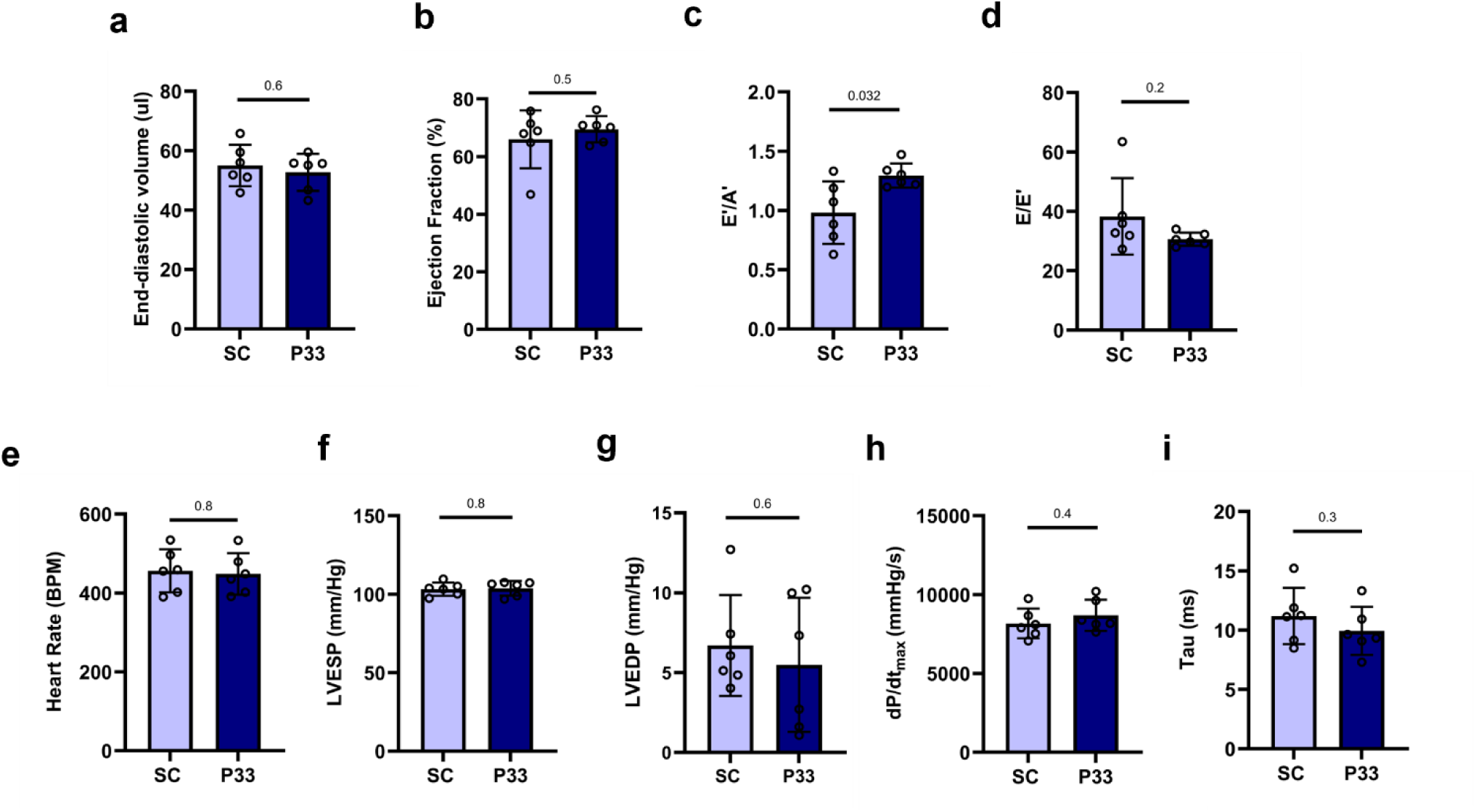
P33 enhance cardiac relaxation *in vivo*. Echocardiophaphy performed **t**hree weeks post injection in healthy mice injected with AAV9 carrying SC or P33 coding sequences, showing **(a)** end diastolic volume, **(b)** LV ejection fraction, the diastolic parameters **(c)** E’/A’ and **(d)** E/E’ derived from combining Pulse-wave Doppler and Tissue Doppler imaging. Left ventricular haemodynamics measured under basal condition for **(e)** heart rates, **(f)** left ventricular end-systolic pressure (LVESP), **(g)** left ventricular end-diastolic pressure (LVEDP), **(h)** the maximum rate of pressure rise (dP/dt_max_) as a measure of contractility, and **(i)** isovolumetric constant, tau, as a measure of relaxation. Statistical significance was assessed using Welch’s t-tests. Each data point represents an individual animal. n = 6 mice per group. Data are presented as mean ± SD.

**Extended Data Figure 18.**
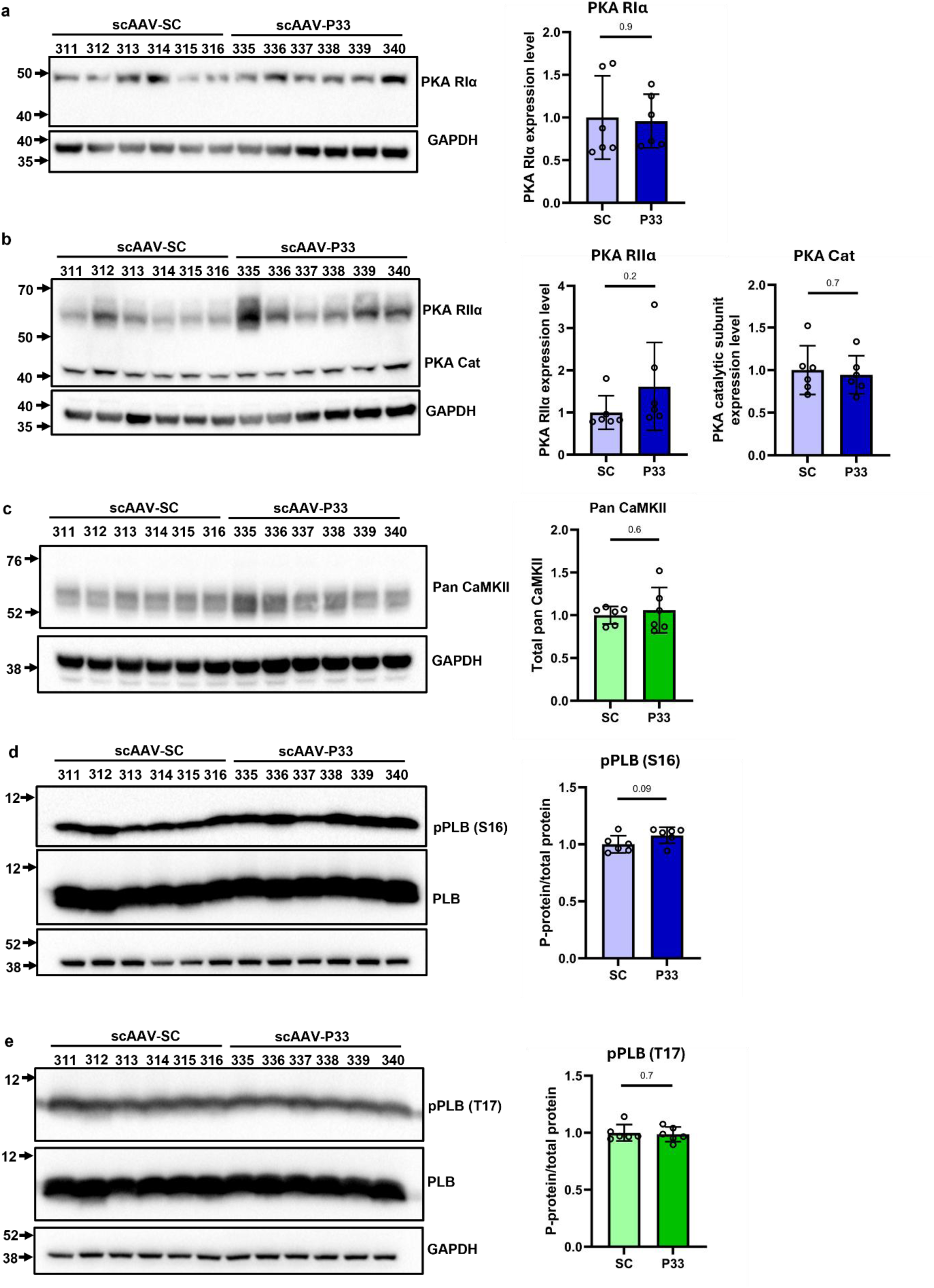
P33 does not alter the abundance of PKA or CaMKII signalling proteins. Western blot analysis and quantification of (a) protein kinase A type Iα regulatory subunit (PKA RIα), (b) protein kinase A type IIα regulatory subunit (PKA RIIα) and catalytic subunit (PKA Cat), and (c) CaMKII in left ventricular (LV) tissue lysates collected from healthy mice 3 weeks after scAAV9-SC or scAAV9-P33 injection. In the same samples, (d) phospholamban (PLB) phosphorylation at Ser16 and (e) Thr17 is shown. Welch’s t-tests. Lane numbers and data points represent individual mice. n = 6 mice per group. Data are presented as mean ± SD.

**Extended Data Figure 19.**
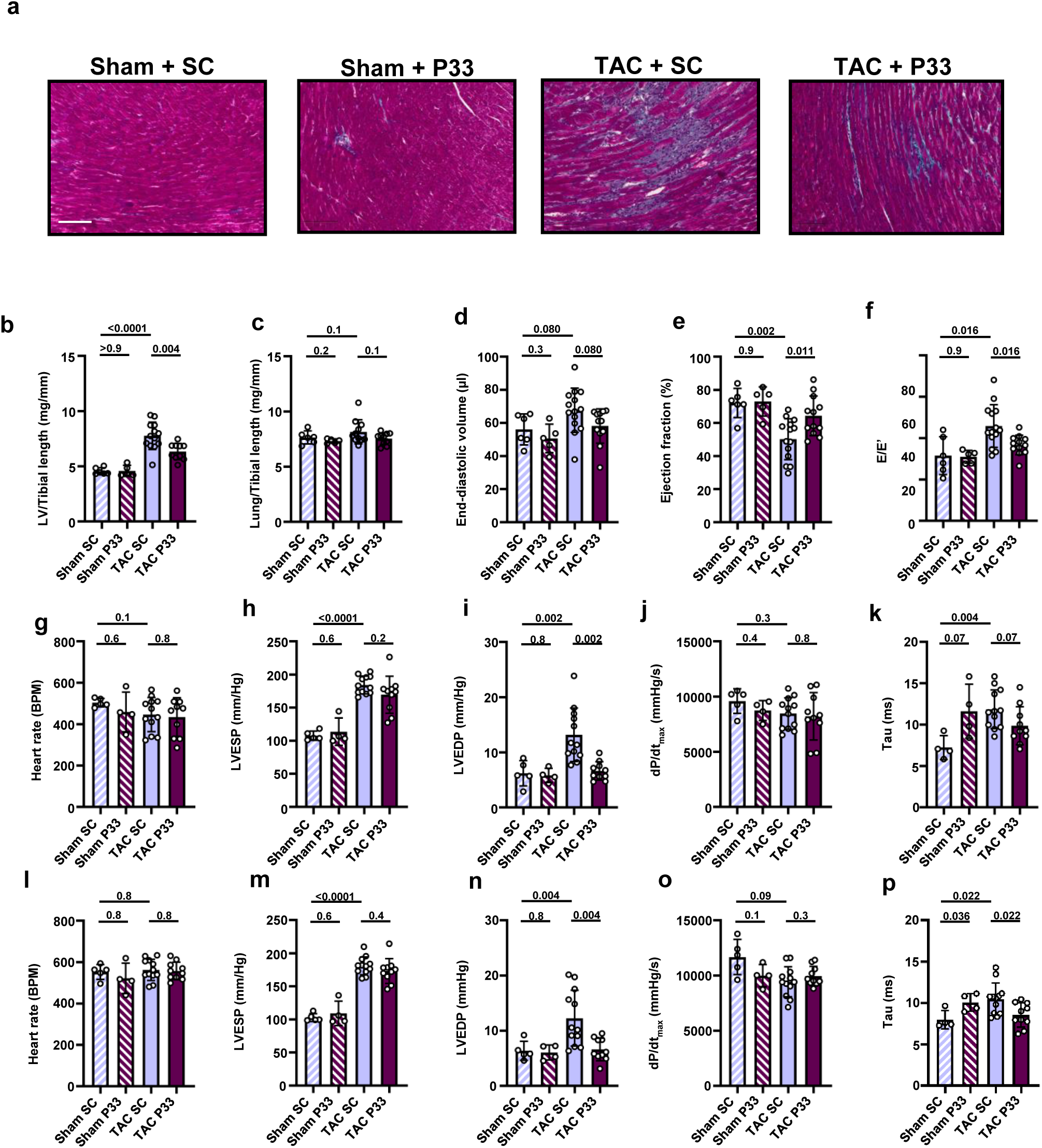
P33 protects from pressure overload-induced cardiac dysfunction in mice subjected to TAC. (**a**) Representative Masson’s trichrome staining of tissue from female mice at week 4 after TAC (bottom; scale bar = 100 μm) (**b**) Left ventricle and (**c**) lung weights obtained post-mortem from male mice and normalised to tibial length. (**d**) Echocardiography recorded in male mice showing LV end diastolic volume, (**e**) LV ejection fraction and (**f**) the diastolic parameter E/e’ derived from combining Pulse-wave Doppler and Tissue Doppler imaging. Haemodynamics measured in male mice under (**g-k**) basal and (**l-p**) cardiac stressor dobutamine (32 ng/g BW/min) for heart rates, left ventricular end-systolic pressure (LVESP), left ventricular end-diastolic pressure (LVEDP), the maximum rate of pressure rise (dP/dt_max_) as a measure of contractility, isovolumetric constant, tau, as a measure of relaxation. Sham-SC N≥ 5, Sham-P33 N ≥ 4, TAC-SC ≥ 12, TAC-P33 ≥ 10. Prespecified comparisons were performed between Sham-SC and Sham-P33, Sham-SC and TAC-SC, and TAC-SC and TAC-P33 using Welch’s t-tests with Benjamini–Hochberg correction. All values are mean ± SD.

**Extended Data Figure 20.**
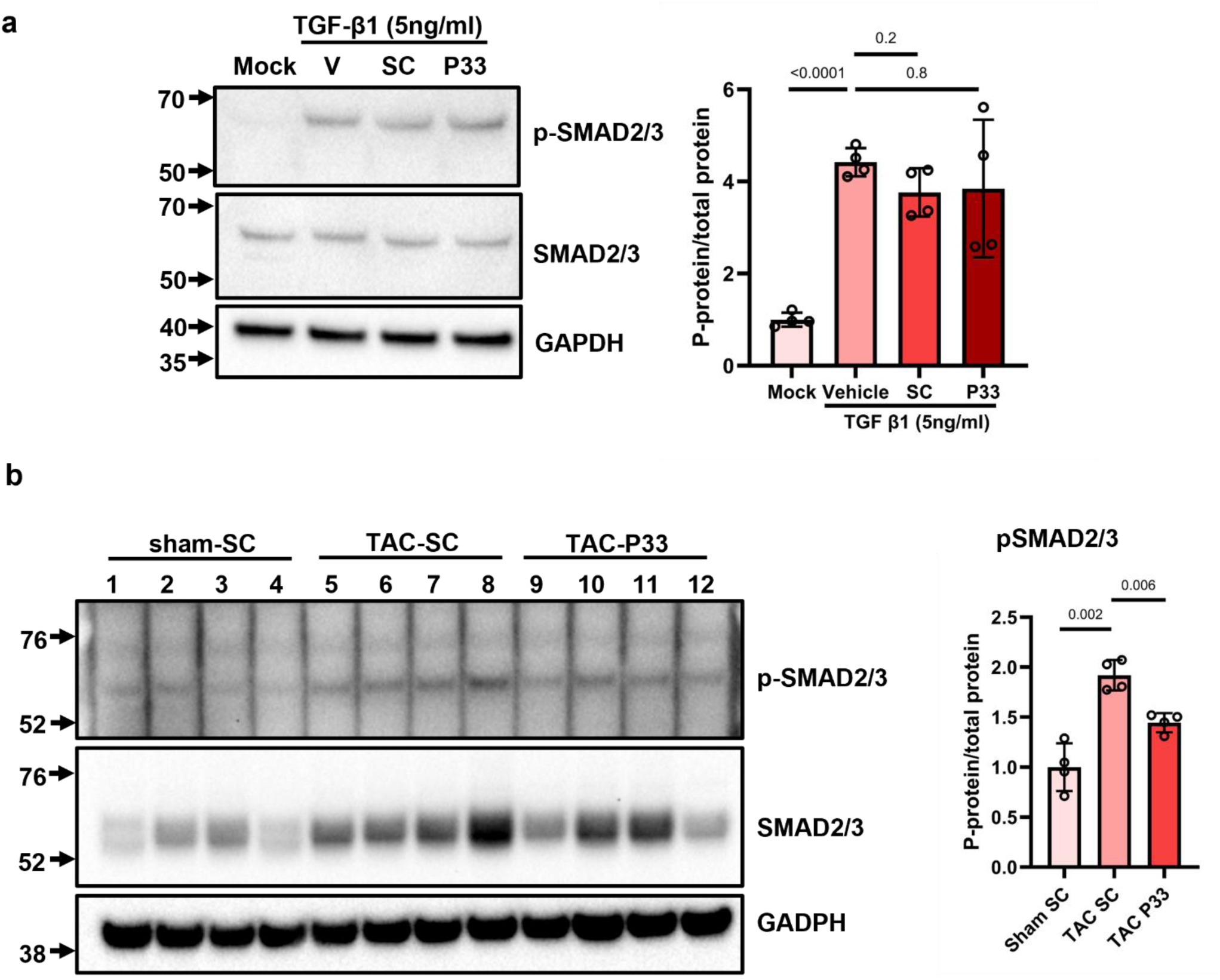
Representative western blot analysis and quantification of phospho-SMAD2/3 and total SMAD2/3 in (**a**) neonatal rat fibroblasts stimulated with TGF-β1 (5 ng/mL) for 30 min in the presence of SC or P33. Mock, PBS-treated control without TGF-β1 stimulation; V, vehicle (DMSO) control with TGF-β1 stimulation; SC and P33, peptide-treated groups with TGF-β1 stimulation. Welch’s ANOVA followed by Dunnett’s T3 multiple-comparison test. n = 4 independent experiments. (**b**) Representative western blot analysis and quantification of LV tissue lysates collected 4 weeks after sham or TAC surgery. Welch’s ANOVA followed by Dunnett’s T3 multiple-comparison test. For panels (b) and (c), each lane and data point represents an individual mouse. Data are presented as mean ± SD.

**Extended Data Figure 21.**
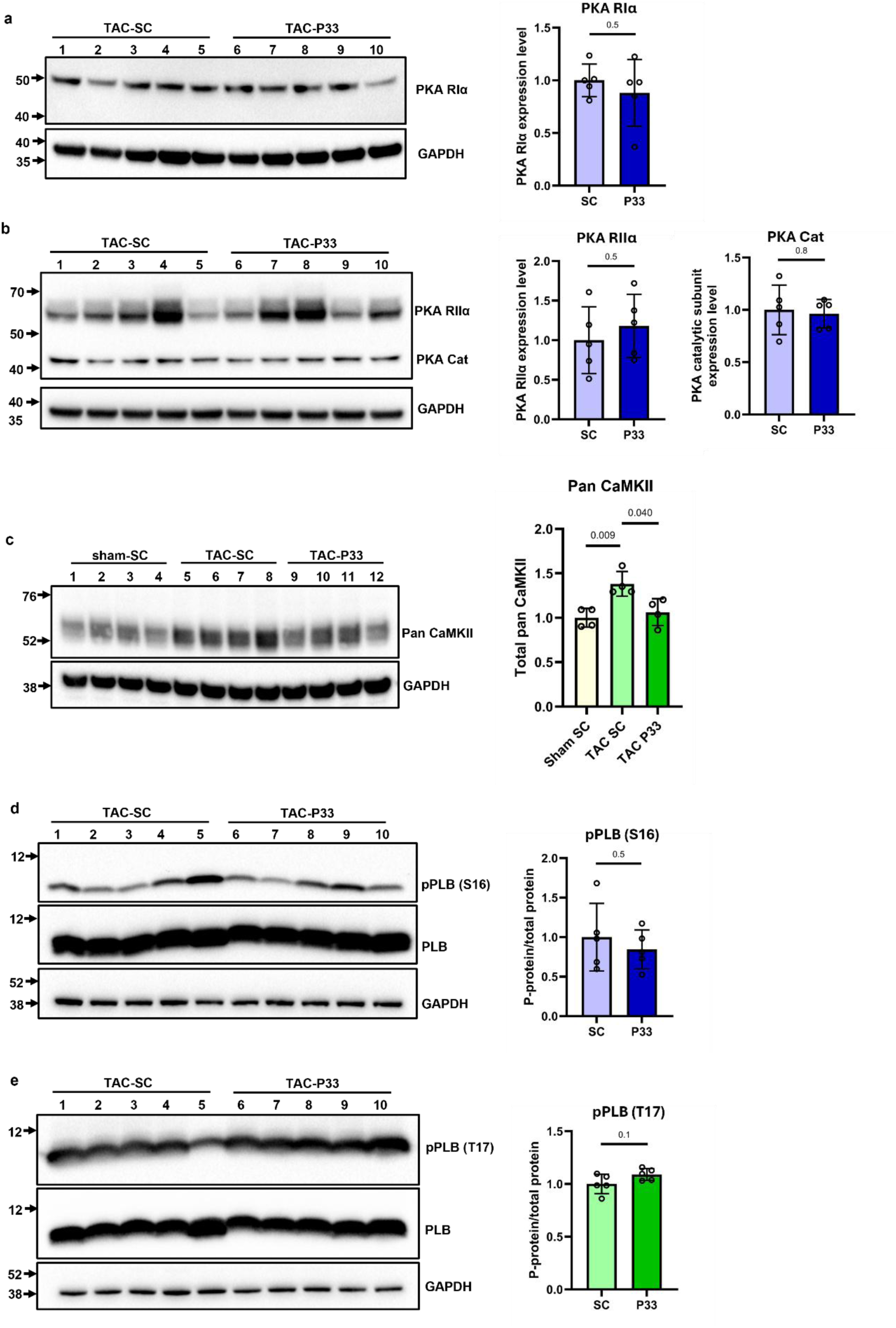
P33 does not alter the abundance of major PKA and CaMKII signalling proteins in TAC hearts. Western blot analysis and quantification of (**a**) PKA RIα, (**b**) PKA RIIα and PKA catalytic subunit (PKA Cat) in left ventricular (LV) tissue lysates collected from mice 4 weeks after TAC surgery and 3 weeks after scAAV9-SC or scAAV9-P33 injection (treatment protocol illustrated in Fig. 5a). Statistical significance was assessed using Welch’s t-tests. (**c**) Representative western blot and quantification of pan-CaMKII in LV tissue lysates collected from sham-operated mice treated with SC and TAC mice treated with either SC or P33. Welch’s ANOVA followed by Dunnett’s T3 multiple-comparison test. (**d**) Phospholamban (PLB) phosphorylation at Ser16 and (**e**) Thr17 measured in the same LV tissue lysates as in (a) and (b). Welch’s t-tests. Lane numbers and data points represent individual mice. n = 6 mice per group. Data are presented as mean ± SD.

**Extended Data Figure 22.**
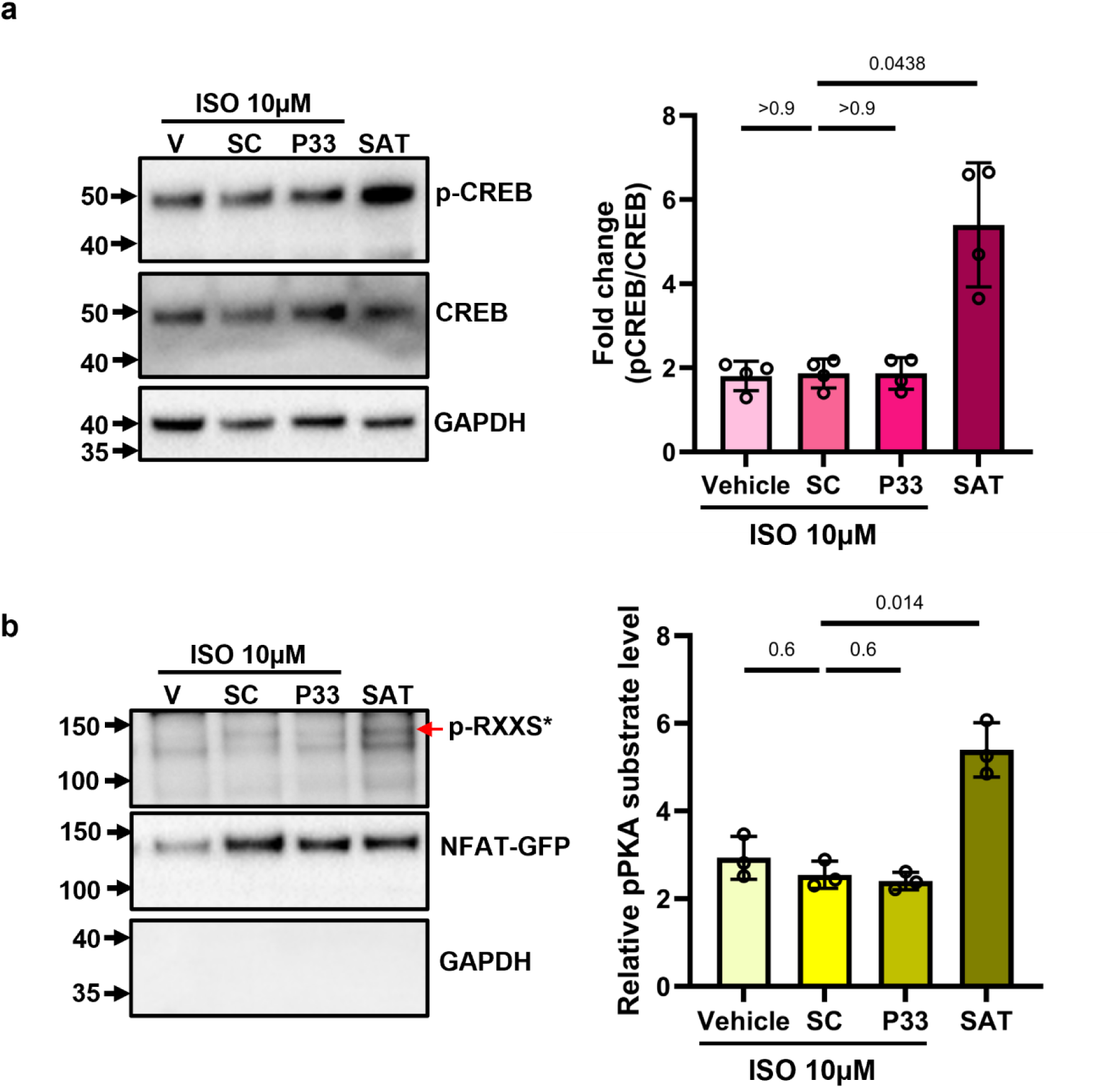
Representative western blots and quantification of (a) phospho-CREB and total CREB in NRVMs stimulated with isoproterenol (ISO, 10 μM) or saturating stimulus (SAT) for 10 min in the presences of SC or P33. V, vehicle (DMSO) control; SC and P33, peptide-treated groups; SAT, saturating stimulus (25 μM FRSK + 100 μM IBMX). Quantification was normalised to the basal group without ISO stimulation, which was set to 1. Welch’s ANOVA followed by Dunnett’s T3 multiple-comparisons test was used for statistical analysis. n = 4 independent experiments. Data are presented as mean ± SD. (**b**) Representative GFP-pull down experiment and quantification from lysate of NRVMs expressing NFAT-GFP. Cells were treated with isoproterenol (ISO, 10 μM) or saturating stimulus (SAT) for 10 min in the presences of SC or P33. Blots were probed with phospho-PKA substrate and GFP, relative phosphorylation levels were normalised to immunoprecipitated NFAT-GFP. Quantification was normalised to the basal group without ISO stimulation, which was set to 1. Welch’s ANOVA followed by Dunnett’s T3 multiple-comparisons test was used for statistical analysis. n = 3 independent experiments. Data are presented as mean ± SD.

**Extended Data Figure 23:**
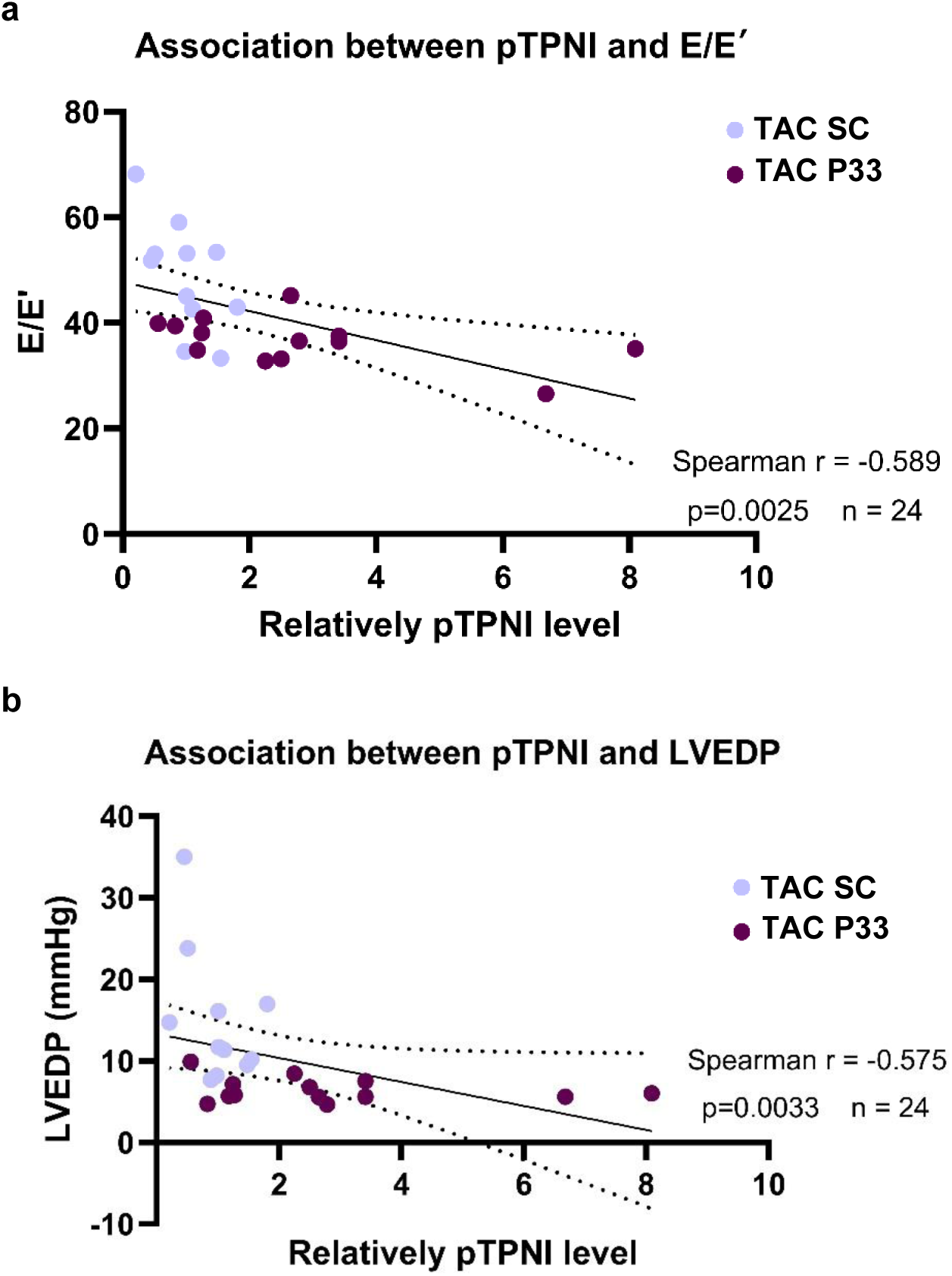
Relation between TPNI phosphorylation (a) E/E′ and (b) LVEDP as measured in male mice subject to TAC surgery and threated for three weeks with either P33 or SC control. Each data point represents an individual animal. Shown are regressions ± 95% confidence interval and *P* values for relations. Correlations were assessed using Spearman’s rank correlation.

**Extended Data Fig. 24.**
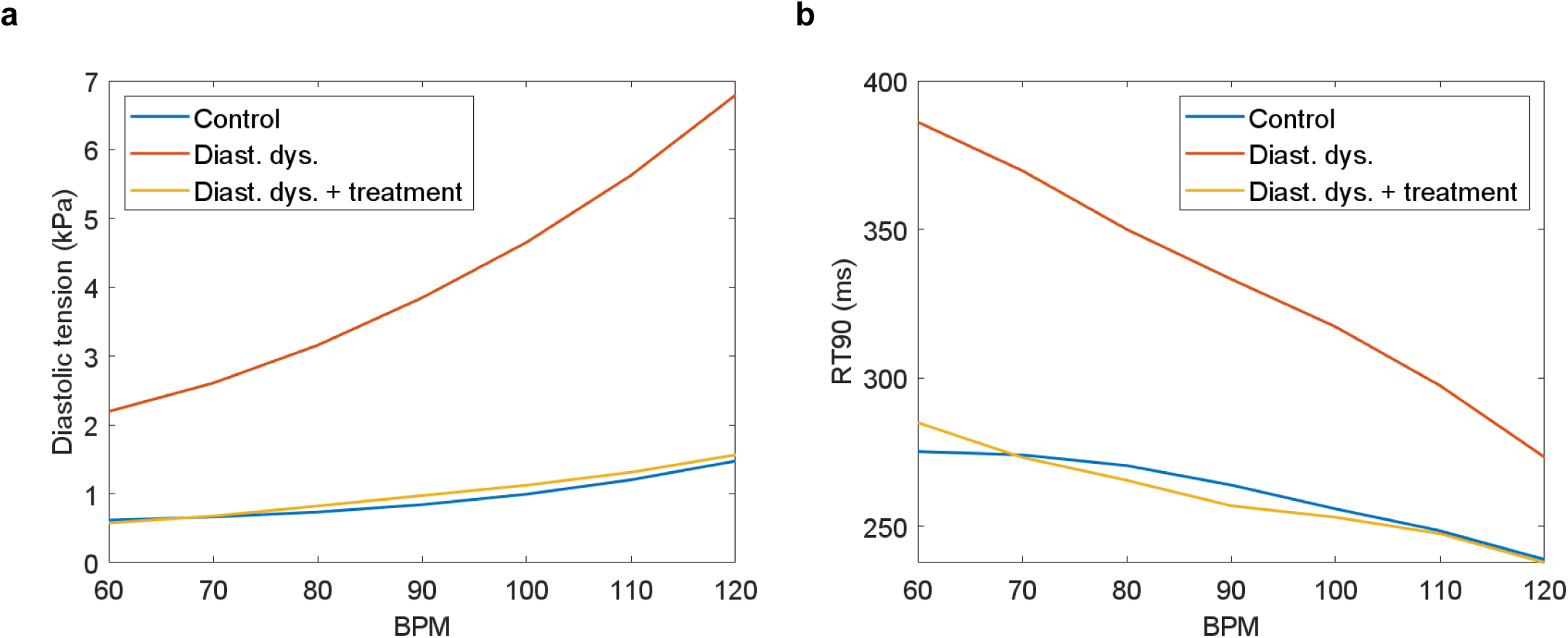
Disrupting PDE4-Troponin interaction normalizes diastolic function. (**a**) Diastolic tension is elevated across heart rates given in beats per minute in a model of diastolic dysfunction; this is normalized by treatment which reduces troponin sensitivity to Calcium. (**b**) A similar pattern in a different index of diastolic dysfunction: time from peak developed force to 90% recovery to baseline value. Here, the baseline is defined as the last value observed in a beat at the given pacing rate.

**Extended Data Table 1.**
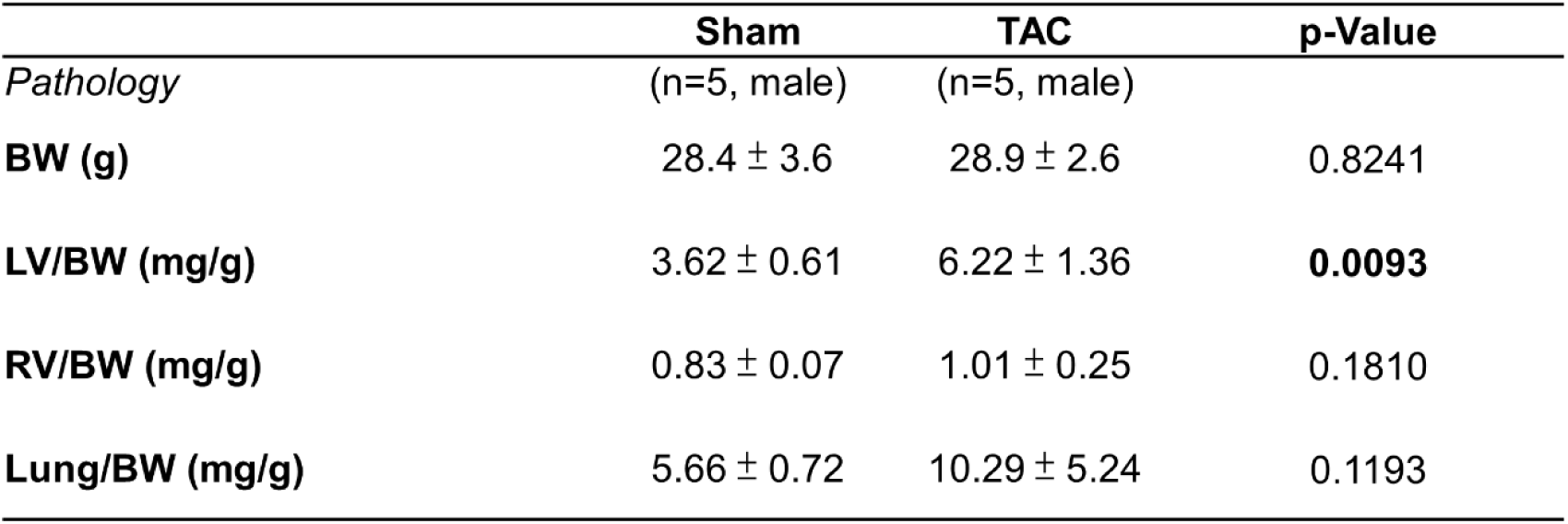
Pathological parameters measured 6.5 weeks after sham or TAC surgery. BW, body weight; LV, left ventricle weight; RV, right ventricle weight. Data are presented as mean ± SD. P-values were calculated using Welch’s t-test for comparisons between the SC and P33 groups.

**Extended Data Table 2.** Echocardiography parameters measured 3 weeks after scAAV9-SC or scAAV9-P33 injections in healthy mice. BW, body weight; LVEDV, left ventricular end-diastolic volume; LVESV, left ventricular end-systolic volume; EF, ejection fraction; FS, fraction shortening; SV, stroke volume; CO, cardiac output; Myo CSA, myocardial cross-sectional area; FAC, fractional area change; E, early diastolic peak filing velocity; E’, peak early relaxation velocity of myocardial diastolic motion. Data are presented as mean ± SD. P-values were calculated using Welch’s t-test for comparisons between the SC and P33 groups.

|  | **SC** | **P33** | **p-value** |
| --- | --- | --- | --- |
| *Echocardiography* | (n=6, male) | (n=6, male) |  |
| **BW (g)** | 28.0 ± 1.1 | 26.3 ± 1.8 | 0.0808 |
| **LVEDV (μl)** | 54.99 ± 7.01 | 52.69 ± 6.24 | 0.5613 |
| **LVESV (μl)** | 18.91 ± 6.90 | 16.22 ± 3.70 | 0.4254 |
| **EF (%)** | 65.96 ± 10.06 | 69.47 ± 4.54 | 0.4618 |
| **FS (%)** | 23.15 ± 6.66 | 22.15 ± 4.59 | 0.7711 |
| **SV (μl)** | 36.09 ± 6.27 | 36.47 ± 3.61 | 0.8997 |
| **CO (ml/min)** | 17.62 ± 4.33 | 18.71 ± 1.93 | 0.5916 |
| **Myo CSA (mm<sup>2</sup>)** | 11.03 ± 1.26 | 10.85 ± 1.40 | 0.8162 |
| **FAC (%)** | 62.30 ± 9.89 | 66.40 ± 6.90 | 0.4265 |
| **E/E'** | 38.29 ± 12.90 | 30.60 ± 2.20 | 0.2067 |

**Extended Data Table 3.** Left ventricular haemodynamic parameters measured 3 weeks after scAAV9-SC or scAAV9-P33 injections in healthy mice. dP/dt_max_, maximum rate of pressure rise (index of contractility); dP/dt_min_, maximum rate of pressure decline (index of relaxation); Tau, isovolumetric relaxation constant. Data are presented as mean ± SD. P-values were calculated using Welch’s t-test for comparisons between the SC and P33 groups.

|  | **SC** | **P33** | **p-value** |
| --- | --- | --- | --- |
| *Baseline* | (n=6, male) | (n=6, male) |  |
| **Heart rate (BPM)** | 456 ± 55 | 448 ± 53 | 0.8010 |
| **LV systolic pressure (mmHg)** | 103.2 ± 4.2 | 103.8 ± 4.7 | 0.8285 |
| **LV diastolic pressure (mmHg)** | 6.7 ± 3.2 | 5.5 ± 4.2 | 0.5855 |
| **dP/dt<sub>max</sub> (mmHg/s)** | 8166 ± 941 | 8686 ± 993 | 0.3732 |
| **dP/dt<sub>min</sub> (mmHg/s)** | 7133 ± 759 | 7618 ± 1109 | 0.4004 |
| **Tau (ms)** | 11.2 ± 2.4 | 9.9 ± 2.0 | 0.3499 |
| *Dobutamine* | (n=6, male) | (n=6, male) |  |
| **Heart rate (BPM)** | 546 ± 36 | 530 ± 49 | 0.5467 |
| **LV systolic pressure (mmHg)** | 98.1 ± 3.7 | 99.4 ± 6.5 | 0.6849 |
| **LV diastolic pressure (mmHg)** | 7.0 ± 3.0 | 6.5 ± 3.7 | 0.8149 |
| **dP/dt<sub>max</sub> (mmHg/s)** | 11573 ± 547 | 12513 ± 1572 | 0.2141 |
| **dP/dt<sub>min</sub> (mmHg/s)** | 8093 ± 913 | 7378 ± 978 | 0.2201 |
| **Tau (ms)** | 8.9 ± 1.4 | 10.4 ± 1.1 | 0.0581 |

**Extended Data Table 4.** Pathological parameters and echocardiography parameters measured in female mice 4 weeks after sham or TAC surgery following with scAAV9-SC/P33 injections 1-week post-surgery. BW, body weight; LVEDV, left ventricular end-diastolic volume; LVESV, left ventricular end-systolic volume; EF, ejection fraction; FS, fraction shortening; SV, stroke volume; CO, cardiac output; Myo CSA, myocardial cross-sectional area; FAC, fractional area change; E, early diastolic peak filling velocity; E’, peak early diastolic myocardial relaxation velocity. Data is mean ± standard deviation. Welch t-test with Benjamini-Hochberg correction for multiple testing. p-values represent the comparison between following groups: ψ = Sham SC vs. Sham P33; # = sham SC vs. TAC SC; * = TAC SC vs TAC P33.

|  | **Sham SC** | **Sham P33** | **TAC SC** | **TAC P33** | **p-Value** |
| --- | --- | --- | --- | --- | --- |
| *Pathology* | (n=5, female) | (n=5, female) | (n=19, female) | (n=19, female) |  |
| **BW (g)** | 21.1 ± 0.6 | 21.7 ± 0.9 | 23.5 ± 1.1 | 23.3 ± 1.2 | 0.4326<sup>ψ</sup>, **0.0001**<sup>#</sup>,<br>0.6580* |
| **LV/Tibial-L (mg/mm)** | 4.46 ± 0.67 | 4.18 ± 0.20 | 7.07 ± 0.71 | 6.12 ± 0.67 | 0.4260<sup>ψ</sup>, **0.0003** <sup>#</sup>,<br>**0.0003** * |
| **RV/Tibial-L (mg/mm)** | 1.00 ± 0.08 | 1.03 ± 0.06 | 1.19 ± 0.14 | 1.16 ± 0.08 | 0.5045<sup>ψ</sup>, **0.0068**<sup>#</sup>,<br>0.05045* |
| **Lung/Tibial-L (mg/mm)** | 7.49 ± 0.46 | 7.71 ± 0.10 | 8.17 ± 0.45 | 8.13 ± 0.37 | 0.5366<sup>ψ</sup>, 0.0728<sup>#</sup>,<br>0.7700* |
| *Echocardiography* |  |  |  |  |  |
| **LVEDV (μl)** | 55 ± 10.2 | 52 ± 8.0 | 70 ± 9.9 | 65 ± 11.3 | 0.6106<sup>ψ</sup>, 0.0648<sup>#</sup>,<br>0.1703* |
| **LVESV (μl)** | 21 ± 4.4 | 17 ± 4.3 | 42 ± 8.7 | 33 ± 10.1 | 0.1955<sup>ψ</sup>, **<0.0001** <sup>#</sup>,<br>**0.0105** * |
| **EF (%)** | 65 ± 4.0 | 68 ± 5.0 | 39 ± 8.3 | 52 ± 8.2 | 0.2982<sup>ψ</sup>, **<0.0001** <sup>#</sup>,<br>**0.0001** * |
| **FS (%)** | 21 ± 2.7 | 22 ± 3.5 | 10 ± 3.5 | 16 ± 3.8 | 0.5625<sup>ψ</sup>, **0.0001** <sup>#</sup>,<br>**0.0001** * |
| **SV (μl)** | 34 ± 6.0 | 35 ± 4.6 | 27 ± 6.1 | 34 ± 5.0 | 0.8095<sup>ψ</sup>, 0.0744<sup>#</sup>,<br>**0.0014** * |
| **CO (ml/min)** | 16.3 ± 3.0 | 16.7 ± 2.1 | 12.8 ± 2.9 | 16.2 ± 2.4 | 0.8233<sup>ψ</sup>, 0.0774<sup>#</sup>,<br>**0.0010** * |
| **Myo CSA (mm<sup>2</sup>)** | 10.5 ± 1.5 | 10.5 ± 1.7 | 15.0 ± 1.4 | 13.2 ± 1.5 | 0.9792<sup>ψ</sup>, **0.0014** <sup>#</sup>,<br>**0.0014** * |
| **FAC (%)** | 55 ± 4.0 | 57 ± 5.1 | 35 ± 5.8 | 43 ± 6.1 | 0.5012<sup>ψ</sup>, **<0.0001** <sup>#</sup>,<br>**0.0003** * |
| **E/E'** | 31.59 ± 6.01 | 26.28 ± 2.88 | 45.00 ± 6.58 | 33.48 ± 7.09 | 0.1272<sup>ψ</sup>, **0.0054** <sup>#</sup>,<br>**<0.0001** * |

**Extended Data Table 5.** LV ventricular haemodynamic parameters measured in female mice 4 weeks after sham or TAC surgery following with scAAV9-SC/P33 injections 1-week post-surgery. dP/dt_max_, maximum rate of pressure rise (index of contractility); dP/dt_min_, maximum rate of pressure decline (index of relaxation); Tau, isovolumetric relaxation constant. Data is mean ± standard deviation. Welch t-test with Benjamini-Hochberg correction for multiple testing. p-values represent the comparison between following groups: ψ = Sham SC vs. Sham P33; # = sham SC vs. TAC SC; * = TAC SC vs TAC P33.

|  | **Sham SC** | **Sham P33** | **TAC SC** | **TAC P33** | **p-Value** |
| --- | --- | --- | --- | --- | --- |
| *Baseline* | (n=5, female) | (n=5, female) | (n=16, female) | (n=17, female) |  |
| **Heart rate (BPM)** | 464 ± 59 | 482 ± 65 | 471 ± 64 | 489 ± 78 | 0.8180<sup>ψ</sup>, 0.8180<sup>#</sup>,<br>0.8180* |
| **LV systolic pressure (mmHg)** | 106.6 ± 1.6 | 105 ± 4.5 | 166.6 ± 15.8 | 169.0 ± 16.6 | 0.6677<sup>ψ</sup>, <0.0001<sup>#</sup>,<br>0.6677* |
| **LV diastolic pressure (mmHg)** | 5.7 ± 0.7 | 5.9 ± 4.1 | 10.4 ± 5.0 | 5.9 ± 1.7 | 0.9076<sup>ψ</sup>, **0.0046<sup>#</sup>**,<br>**0.0046\*** |
| **dP/dt<sub>max</sub> (mmHg/s)** | 9002 ± 1023 | 8762 ± 1999 | 8339 ± 1487 | 8890 ± 1453 | 0.8189<sup>ψ</sup>, 0.4358<sup>#</sup>,<br>0.4358* |
| **dP/dt<sub>min</sub> (mmHg/s)** | 7243 ± 854 | 7480 ± 2386 | 7690 ± 1982 | 9360 ± 2242 | 0.8425<sup>ψ</sup>, 0.7276<sup>#</sup>,<br>0.0909* |
| **Tau (ms)** | 9.8 ± 1.7 | 11.0 ± 5.0 | 12.0 ± 4.1 | 8.4 ± 1.3 | 0.6201<sup>ψ</sup>, 0.1629<sup>#</sup>,<br>**0.0122\*** |
| *Dobutamine* |  |  |  |  |  |
| **Heart rate (BPM)** | 509 ± 60 | 535 ± 58 | 522 ± 61 | 564 ± 33 | 0.6867<sup>ψ</sup>, 0.6867<sup>#</sup>,<br>0.0706* |
| **LV systolic pressure (mmHg)** | 101.4 ± 2.7 | 107.4 ± 11.6 | 169.0 ± 20.2 | 176.4 ± 16.4 | 0.3188<sup>ψ</sup>, <0.0001<sup>#</sup>,<br>0.3188* |
| **LV diastolic pressure (mmHg)** | 6.1 ± 1.1 | 6.6 ± 2.6 | 10.0 ± 4.5 | 6.0 ± 1.9 | 0.6981<sup>ψ</sup>, **0.0088<sup>#</sup>**,<br>**0.0088\*** |
| **dP/dt<sub>max</sub> (mmHg/s)** | 10556 ± 1179 | 11028 ± 2505 | 9647 ± 2098 | 10973 ± 1260 | 0.7172<sup>ψ</sup>, 0.3655<sup>#</sup>,<br>0.1164* |
| **dP/dt<sub>min</sub> (mmHg/s)** | 6448 ± 1075 | 7895 ± 1430 | 8623 ± 2237 | 10576 ± 2241 | 0.1112<sup>ψ</sup>, **0.0267<sup>#</sup>**,<br>**0.0267\*** |
| **Tau (ms)** | 10.8 ± 1.9 | 9.6 ± 2.2 | 9.7 ± 2.4 | 8.0 ± 1.3 | 0.3827<sup>ψ</sup>, 0.3827<sup>#</sup>,<br>**0.0796\*** |

**Extended Data Table 6.** Pathological parameters and echocardiography parameters measured in male mice 4 weeks after sham or TAC surgery following with scAAV9-SC/P33 injections 1-week post-surgery. BW, body weight; LVEDV, left ventricular end-diastolic volume; LVESV, left ventricular end-systolic volume; EF, ejection fraction; FS, fraction shortening; SV, stroke volume; CO, cardiac output; Myo CSA, myocardial cross-sectional area; FAC, fractional area change; E, early diastolic peak filling velocity; E’, peak early diastolic myocardial relaxation velocity. Data is mean ± standard deviation. Welch t-test with Benjamini-Hochberg correction for multiple testing. p-values represent the comparison between following groups: ψ = Sham SC vs. Sham P33; # = sham SC vs. TAC SC; * = TAC SC vs TAC P33.

|  | **Sham SC** | **Sham P33** | **TAC SC** | **TAC P33** | **p-Value** |
| --- | --- | --- | --- | --- | --- |
| *Pathology* | (n=6, male) | (n=5, male) | (n=13, male) | (n=10, male) |  |
| **BW (g)** | 26.5 ± 2.0 | 26.0 ± 1.7 | 27.4 ± 2.5 | 26.2 ± 1.6 | 0.6344<sup>ψ</sup>, 0.6344<sup>#</sup>,<br>0.5001* |
| **LV/Tibial-L (mg/mm)** | 4.56 ± 0.31 | 4.58 ± 0.50 | 7.76 ± 1.24 | 6.34 ± 0.74 | 0.9287<sup>ψ</sup>, **<0.0001<sup>#</sup>**,<br>**0.0039\*** |
| **RV/Tibial-L (mg/mm)** | 1.36 ± 0.10 | 1.25 ± 0.10 | 1.60 ± 0.29 | 1.48 ± 0.32 | 0.1123<sup>ψ</sup>, 0.0561<sup>#</sup>,<br>0.1123* |
| **Lung/Tibial-L (mg/mm)** | 7.66 ± 0.59 | 7.33 ± 0.20 | 9.17 ± 2.68 | 8.31 ± 2.57 | 0.2488<sup>ψ</sup>, 0.1095<sup>#</sup>,<br>0.1095* |
| *Echocardiography* | (n=6, male) | (n=5, male) | (n=14, male) | (n=12, male) |  |
| **LVEDV (μl)** | 56 ± 9.4 | 51 ± 8.6 | 68 ± 13.4 | 58 ± 10.3 | 0.3427<sup>ψ</sup>, 0.0803<sup>#</sup>,<br>0.0803* |
| **LVESV (μl)** | 16 ± 7.1 | 14 ± 7.1 | 32 ± 14.8 | 21 ± 8.8 | 0.6582<sup>ψ</sup>, **0.0127<sup>#</sup>**,<br>**0.0490\*** |
| **EF (%)** | 72 ± 8.8 | 73 ± 8.8 | 50 ± 12.3 | 64 ± 12.0 | 0.8702<sup>ψ</sup>, **0.0017<sup>#</sup>**,<br>**0.0105\*** |
| **FS (%)** | 24 ± 3.0 | 25 ± 4.2 | 9 ± 2.0 | 18 ± 1.5 | 0.6850<sup>ψ</sup>, **<0.0001<sup>#</sup>**,<br>**<0.0001\*** |
| **SV (μl)** | 40 ± 5.0 | 36 ± 3.4 | 33 ± 7.0 | 37 ± 6.5 | 0.2053<sup>ψ</sup>, 0.0736<sup>#</sup>,<br>0.2053* |
| **CO (ml/min)** | 20.1 ± 3.3 | 17.9 ± 1.8 | 16.0 ± 3.4 | 17.6 ± 3.4 | 0.2391<sup>ψ</sup>, 0.0941<sup>#</sup>,<br>0.2391* |
| **Myo CSA (mm<sup>2</sup>)** | 12.3 ± 1.5 | 12.5 ± 1.7 | 18.3 ± 2.5 | 16.8 ± 2.1 | 0.8797<sup>ψ</sup>, **<0.0001<sup>#</sup>**,<br>0.1586* |
| **FAC (%)** | 58 ± 11.9 | 53 ± 7.1 | 39 ± 6.6 | 50 ± 9.3 | 0.4213<sup>ψ</sup>, **0.0169<sup>#</sup>**,<br>**0.0131\*** |
| **E/E'** | 31.54 ± 9.12 | 30.97 ± 3.10 | 45.84 ± 10.27 | 36.7 ± 4.73 | 0.8906<sup>ψ</sup>, **0.0159<sup>#</sup>**,<br>**0.0159\*** |

**Extended Data Table 7.** LV ventricular haemodynamic parameters measured in male mice 4 weeks after sham or TAC surgery following with scAAV9-SC/P33 injections 1-week post-surgery. dP/dt_max_, maximum rate of pressure rise (index of contractility); dP/dt_min_, maximum rate of pressure decline (index of relaxation); Tau, isovolumetric relaxation constant. Data is mean ± standard deviation. Welch t-test with Benjamini-Hochberg correction for multiple testing. p-values represent the comparison between following groups: ψ = Sham SC vs. Sham P33; # = sham SC vs. TAC SC; * = TAC SC vs TAC P33.

|  | **Sham SC** | **Sham P33** | **TAC SC** | **TAC P33** | **p-Value** |
| --- | --- | --- | --- | --- | --- |
| *Baseline* | (n=5, male) | (n=4, male) | (n=12, male) | (n=10, male) |  |
| **Heart rate (BPM)** | 504 ± 21 | 459 ± 96 | 447 ± 82 | 435 ± 93 | 0.6319<sup>ψ</sup>, 0.1246<sup>#</sup>,<br>0.7544* |
| **LV systolic pressure (mmHg)** | 107.7 ± 7.0 | 113.6 ± 20.7 | 184.1 ± 13.5 | 169.5 ± 27.9 | 0.6179<sup>ψ</sup>, **<0.0001<sup>#</sup>**,<br>0.2300* |
| **LV diastolic pressure (mmHg)** | 6.2 ± 2.3 | 5.8 ± 1.3 | 13.2 ± 4.8 | 6.6 ± 1.7 | 0.7648<sup>ψ</sup>, **0.0018<sup>#</sup>**,<br>**0.0018\*** |
| **dP/dt<sub>max</sub> (mmHg/s)** | 9587 ± 1118 | 8728 ± 907 | 8460 ± 1427 | 8210 ± 2150 | 0.3656<sup>ψ</sup>, 0.3408<sup>#</sup>,<br>0.7573* |
| **dP/dt<sub>min</sub> (mmHg/s)** | 7607 ± 1999 | 6784 ± 1333 | 8079 ± 1558 | 8873 ± 2361 | 0.6533<sup>ψ</sup>, 0.6533<sup>#</sup>,<br>0.6533* |
| **Tau (ms)** | 7.2 ± 1.4 | 11.6 ± 3.3 | 11.8 ± 2.3 | 9.9 ± 2.3 | 0.0678<sup>ψ</sup>, **0.0038<sup>#</sup>**,<br>0.0678* |
| *Dobutamine* |  |  |  |  |  |
| **Heart rate (BPM)** | 553 ± 36 | 522 ± 74 | 563 ± 51 | 557 ± 43 | 0.7916<sup>ψ</sup>, 0.7916<sup>#</sup>,<br>0.7916* |
| **LV systolic pressure (mmHg)** | 102.9 ± 6.0 | 109.1 ± 18.5 | 180.9 ± 13.5 | 173.1 ± 18.7 | 0.5573<sup>ψ</sup>, **<0.0001<sup>#</sup>**,<br>0.4290* |
| **LV diastolic pressure (mmHg)** | 6.4 ± 1.7 | 6.0 ± 1.4 | 12.3 ± 5.1 | 6.6 ± 2.1 | 0.7726<sup>ψ</sup>, **0.0044<sup>#</sup>**,<br>**0.0044\*** |
| **dP/dt<sub>max</sub> (mmHg/s)** | 11677 ± 1587 | 9990 ± 1002 | 9440 ± 1360 | 9953 ± 876 | 0.1422<sup>ψ</sup>, 0.0895<sup>#</sup>,<br>0.2992* |
| **dP/dt<sub>min</sub> (mmHg/s)** | 7570 ± 1883 | 7233 ± 905 | 8780 ± 1731 | 9849 ± 2228 | 0.7363<sup>ψ</sup>, 0.3851<sup>#</sup>,<br>0.3851* |
| **Tau (ms)** | 8.0 ± 1.1 | 10.1 ± 1.1 | 10.5 ± 1.9 | 8.5 ± 1.5 | **0.0364<sup>ψ</sup>**, **0.0220<sup>#</sup>**,<br>**0.0220\*** |

**Extended Data Table 8.** Clinical characteristics of the patients employed for the FRET experiments in the study. LVEF, left ventricle ejection fraction; ICM, ischemic cardiomyopathy; DCM, dilated cardiomyopathy; LVAD, left ventricular assist device implantation; Impella, percutaneous ventricular assist device.

| **Clinical features** |  |
| --- | --- |
| Age, years | 51.19 ± 12.80 |
| Male gender, n (%) | 11 (68.75) |
| Body mass index, kg/m<sup>2</sup> | 26.44 ± 6.61 |
| **Cardiovascular disease** |  |
| Sinus rhythm, n (%) | 7 (43.75%) |
| Atrial fibrillation, n (%) | 9 (56.25%) |
| LVEF, % | 26.31 ± 6.38 |
| ICM, n (%) | 1 (6.25%) |
| DCM, n (%) | 1 (6.25%) |
| **Surgical intervention** |  |
| Heart transplant, n (%) | 8 (50.00%) |
| LVAD/Impella, n (%) | 5 (31.25%) |
| Valvular surgery, n (%) | 5 (31.25%) |
| Myectomy, n (%) | 2 (12.50%) |
| **Medication** |  |
| ACE inhibitors, n (%) | 5 (31.25%) |
| Beta-blockers, n (%) | 12 (75.00%) |
| Diuretics, n (%) | 12 (75.00%) |
| Calcium antagonists, n (%) | 1 (6.25%) |
| Statins, n (%) | 6 (37.50%) |
| AT<sub>1</sub> receptor inhibitors, n (%) | 5 (31.25%) |

**Extended Data Table 9.** Clinical characteristics of the patients employed for co-immunoprecipitation experiments in the study. Surgical procedure was in all cases heart transplant and samples were obtained from the explanted (Control) or from the implanted hearts (HF). LVEF, left ventricle ejection fraction; HF, heart failure with reduced ejection fraction; ICM, ischemic cardiomyopathy; DCM, dilated cardiomyopathy. Continuous variables are presented as mean ± SEM and categorical variables as counts.

| Clinical features |  |  |
| --- | --- | --- |
|  | Control (n = 3) | HF (n = 7) |
| Age, years | 48.67 ± 28.10 | 59.43 ± 22.46 |
| Male gender, n (%) | 1 (33.33%) | 5 (71.43%) |
| Body mass index, kg/m<sup>2</sup> | 25.00 ± 17.68 | 26.43 ± 9.99 |
| Cardiovascular disease |  |  |
| Sinus rhythm, n (%) | 3 (100.00%) | 4 (57.14%) |
| Atrial fibrillation, n (%) | 0 (0.00%) | 3 (42.86%) |
| LVEF, % | 58.33 ± 33.68 | 23.43 ± 0.05 |
| ICM, n (%) | 0 (0.00%) | 4 (57.14%) |
| DCM, n (%) | 0 (0.00%) | 2 (28.57%) |
| Medication |  |  |
| ACE inhibitors, n (%) | 0 (0.00%) | 3 (42.86%) |
| Beta-blockers, n (%) | 0 (0.00%) | 7 (100.00%) |
| Diuretics, n (%) | 0 (0.00%) | 6 (85.71%) |
| Calcium antagonists, n (%) | 0 (0.00%) | 1 (14.29%) |
| Statins, n (%) | 0 (0.00%) | 3 (42.86%) |
| AT<sub>1</sub> receptor inhibitors, n (%) | 0 (0.00%) | 1 (14.29%) |

## References

1 Bers, D. M. Cardiac excitation-contraction coupling. Nature 415, 198–205 (2002). 10.1038/415198a

2 Bristow, M. R. et al. Decreased catecholamine sensitivity and beta-adrenergic-receptor density in failing human hearts. N Engl J Med 307, 205–211 (1982). 10.1056/NEJM198207223070401

3 Neubauer, S. The failing heart--an engine out of fuel. N Engl J Med 356, 1140–1151 (2007). 10.1056/NEJMra063052

4 Mishra, S. & Kass, D. A. Cellular and molecular pathobiology of heart failure with preserved ejection fraction. Nat Rev Cardiol 18, 400–423 (2021). 10.1038/s41569-020-00480-6

5 Zaccolo, M. & Kovanich, D. Nanodomain cAMP signalling in cardiac pathophysiology: potential for developing targeted therapeutic interventions. Physiol Rev (2024). 10.1152/physrev.00013.2024

6 Surdo, N. C. et al. FRET biosensor uncovers cAMP nano-domains at beta-adrenergic targets that dictate precise tuning of cardiac contractility. Nat Commun 8, 15031 (2017). 10.1038/ncomms15031

7 Scott, J. D. & Santana, L. F. A-kinase anchoring proteins: getting to the heart of the matter. Circulation 121, 1264–1271 (2010).

8 Kamel, R., Leroy, J., Vandecasteele, G. & Fischmeister, R. Cyclic nucleotide phosphodiesterases as therapeutic targets in cardiac hypertrophy and heart failure. Nat Rev Cardiol 20, 90–108 (2023). 10.1038/s41569-022-00756-z

9 Kokkonen, K. & Kass, D. A. Nanodomain Regulation of Cardiac Cyclic Nucleotide Signaling by Phosphodiesterases. Annu Rev Pharmacol Toxicol 57, 455–479 (2017). 10.1146/annurev-pharmtox-010716-104756

10 Lomas, O. et al. Adenoviral transduction of FRET-based biosensors for cAMP in primary adult mouse cardiomyocytes. Methods Mol Biol 1294, 103–115 (2015). 10.1007/978-1-4939-2537-7_8

11 Zaccolo, M. & Pozzan, T. Discrete microdomains with high concentration of cAMP in stimulated rat neonatal cardiac myocytes. Science 295, 1711–1715 (2002).

12 Stangherlin, A. et al. cGMP signals modulate cAMP levels in a compartment-specific manner to regulate catecholamine-dependent signaling in cardiac myocytes. Circ Res 108, 929–939 (2011). 10.1161/CIRCRESAHA.110.230698

13 Henderson, D. J. et al. The cAMP phosphodiesterase-4D7 (PDE4D7) is downregulated in androgen-independent prostate cancer cells and mediates proliferation by compartmentalising cAMP at the plasma membrane of VCaP prostate cancer cells. Br J Cancer 110, 1278–1287 (2014). 10.1038/bjc.2014.22

14 Cooke, S. F. et al. Disruption of the pro-oncogenic c-RAF-PDE8A complex represents a differentiated approach to treating KRAS-c-RAF dependent PDAC. Sci Rep 14, 8998 (2024). 10.1038/s41598-024-59451-3

15 Brown, K. M. et al. Phosphodiesterase-8A binds to and regulates Raf-1 kinase. Proc Natl Acad Sci U S A 110, E1533–1542 (2013). 10.1073/pnas.1303004110

16 Amartely, H., Iosub-Amir, A. & Friedler, A. Identifying protein-protein interaction sites using peptide arrays. J Vis Exp, e52097 (2014). 10.3791/52097

17 Maddock, H. L., Mocanu, M. M. & Yellon, D. M. Adenosine A(3) receptor activation protects the myocardium from reperfusion/reoxygenation injury. American journal of physiology. Heart and circulatory physiology 283, H1307–1313 (2002). 10.1152/ajpheart.00851.2001

18 Tran, P. et al. Profiling the Biomechanical Responses to Workload on the Human Myocyte to Explore the Concept of Myocardial Fatigue and Reversibility: Rationale and Design of the POWER Heart Failure Study. J Cardiovasc Transl Res 17, 275–286 (2024). 10.1007/s12265-023-10391-9

19 Gharanei M, B. M., Linekar A, Wallis R, Chuizbaian O, Maddock H.. Development of an in vitro platform using the human primary cardiomyocyte work loop assay to screen for drug-induced effects on cardiac contractility. Journal of Pharmacological and Toxicological Methods 93 (2018). 10.1016/j.vascn.2018.01.428

20 Helmes, M. et al. Mimicking the cardiac cycle in intact cardiomyocytes using diastolic and systolic force clamps; measuring power output. Cardiovasc Res 111, 66–73 (2016). 10.1093/cvr/cvw072

21 Margara, F. et al. Mechanism based therapies enable personalised treatment of hypertrophic cardiomyopathy. Sci Rep 12, 22501 (2022). 10.1038/s41598-022-26889-2

22 Psaras, Y. et al. CalTrack: High-Throughput Automated Calcium Transient Analysis in Cardiomyocytes. Circ Res 129, 326–341 (2021). 10.1161/CIRCRESAHA.121.318868

23 Toepfer, C. N. et al. SarcTrack. Circ Res 124, 1172–1183 (2019). 10.1161/CIRCRESAHA.118.314505

24 Zufferey, R., Donello, J. E., Trono, D. & Hope, T. J. Woodchuck hepatitis virus posttranscriptional regulatory element enhances expression of transgenes delivered by retroviral vectors. J Virol 73, 2886–2892 (1999). 10.1128/JVI.73.4.2886-2892.1999

25 Choi, J. H. et al. Optimization of AAV expression cassettes to improve packaging capacity and transgene expression in neurons. Mol Brain 7, 17 (2014). 10.1186/1756-6606-7-17

26 Lygate CA, N. S. Surgically induced chronic heart failure. A Handbook of Mouse Models of Cardiovascular Disease, 333–348 (2006).

27 Lygate, C. A. et al. Serial high resolution 3D-MRI after aortic banding in mice: band internalization is a source of variability in the hypertrophic response. Basic Res Cardiol 101, 8–16 (2006). 10.1007/s00395-005-0546-3

28 Faller, K. M. E. et al. Impaired cardiac contractile function in arginine:glycine amidinotransferase knockout mice devoid of creatine is rescued by homoarginine but not creatine. Cardiovasc Res 114, 417–430 (2018). 10.1093/cvr/cvx242

29 Bankhead, P. et al. QuPath: Open source software for digital pathology image analysis. Sci Rep 7, 16878 (2017). 10.1038/s41598-017-17204-5

30 Subramaniam, G. et al. Integrated Proteomics Unveils Nuclear PDE3A2 as a Regulator of Cardiac Myocyte Hypertrophy. Circ Res 132, 828–848 (2023). 10.1161/CIRCRESAHA.122.321448

31 Mongillo, M. et al. Fluorescence resonance energy transfer-based analysis of cAMP dynamics in live neonatal rat cardiac myocytes reveals distinct functions of compartmentalized phosphodiesterases. Circ Res 95, 67–75 (2004).

32 Li, X., Huston, E., Lynch, M. J., Houslay, M. D. & Baillie, G. S. Phosphodiesterase-4 influences the PKA phosphorylation status and membrane translocation of G-protein receptor kinase 2 (GRK2) in HEK-293beta2 cells and cardiac myocytes. Biochem J 394, 427–435 (2006). 10.1042/BJ20051560

33 Richter, W., Jin, S. L. & Conti, M. Splice variants of the cyclic nucleotide phosphodiesterase PDE4D are differentially expressed and regulated in rat tissue. Biochem J 388, 803–811 (2005). 10.1042/BJ20050030

34 McCahill, A. et al. In resting COS1 cells a dominant negative approach shows that specific, anchored PDE4 cAMP phosphodiesterase isoforms gate the activation, by basal cyclic AMP production, of AKAP-tethered protein kinase A type II located in the centrosomal region. Cell Signal 17, 1158–1173 (2005).

35 Li, X. et al. Selective SUMO modification of cAMP-specific phosphodiesterase-4D5 (PDE4D5) regulates the functional consequences of phosphorylation by PKA and ERK. Biochem J 428, 55–65 (2010). 10.1042/BJ20091672

36 Tibbo, A. J. et al. Phosphodiesterase type 4 anchoring regulates cAMP signaling to Popeye domain-containing proteins. J Mol Cell Cardiol 165, 86–102 (2022). 10.1016/j.yjmcc.2022.01.001

37 Yumoto, F. et al. Drastic Ca2+ sensitization of myofilament associated with a small structural change in troponin I in inherited restrictive cardiomyopathy. Biochemical and biophysical research communications 338, 1519–1526 (2005). 10.1016/j.bbrc.2005.10.116

38 Klarenbeek, J., Goedhart, J., van Batenburg, A., Groenewald, D. & Jalink, K. Fourth-generation epac-based FRET sensors for cAMP feature exceptional brightness, photostability and dynamic range: characterization of dedicated sensors for FLIM, for ratiometry and with high affinity. PloS one 10, e0122513 (2015). 10.1371/journal.pone.0122513

39 Layland, J. et al. Essential role of troponin I in the positive inotropic response to isoprenaline in mouse hearts contracting auxotonically. J Physiol 556, 835–847 (2004). 10.1113/jphysiol.2004.061176

40 Kentish, J. C. et al. Phosphorylation of troponin I by protein kinase A accelerates relaxation and crossbridge cycle kinetics in mouse ventricular muscle. Circ Res 88, 1059–1065 (2001). 10.1161/hh1001.091640

41 Herron, T. J., Korte, F. S. & McDonald, K. S. Power output is increased after phosphorylation of myofibrillar proteins in rat skinned cardiac myocytes. Circ Res 89, 1184–1190 (2001). 10.1161/hh2401.101908

42 Fentzke, R. C. et al. Impaired cardiomyocyte relaxation and diastolic function in transgenic mice expressing slow skeletal troponin I in the heart. J Physiol 517 (Pt 1), 143–157 (1999). 10.1111/j.1469-7793.1999.0143z.x

43 Barbato, J. C., Huang, Q. Q., Hossain, M. M., Bond, M. & Jin, J. P. Proteolytic N-terminal truncation of cardiac troponin I enhances ventricular diastolic function. J Biol Chem 280, 6602–6609 (2005). 10.1074/jbc.M408525200

44 Venkataraman, R. et al. Myofilament calcium de-sensitization and contractile uncoupling prevent pause-triggered ventricular tachycardia in mouse hearts with chronic myocardial infarction. Journal of molecular and cellular cardiology 60, 8–15 (2013). 10.1016/j.yjmcc.2013.03.022

45 de Waard, M. C. et al. Early exercise training normalizes myofilament function and attenuates left ventricular pump dysfunction in mice with a large myocardial infarction. Circ Res 100, 1079–1088 (2007). 10.1161/01.RES.0000262655.16373.37

46 van der Velden, J. et al. Alterations in myofilament function contribute to left ventricular dysfunction in pigs early after myocardial infarction. Circ Res 95, e85–95 (2004). 10.1161/01.RES.0000149531.02904.09

47 McConnell, B. K., Moravec, C. S., Morano, I. & Bond, M. Troponin I phosphorylation in spontaneously hypertensive rat heart: effect of beta-adrenergic stimulation. Am J Physiol 273, H1440–1451 (1997). 10.1152/ajpheart.1997.273.3.H1440

48 Wolff, M. R., Buck, S. H., Stoker, S. W., Greaser, M. L. & Mentzer, R. M. Myofibrillar calcium sensitivity of isometric tension is increased in human dilated cardiomyopathies: role of altered beta-adrenergically mediated protein phosphorylation. J Clin Invest 98, 167–176 (1996). 10.1172/JCI118762

49 Bodor, G. S. et al. Troponin I phosphorylation in the normal and failing adult human heart. Circulation 96, 1495–1500 (1997). 10.1161/01.cir.96.5.1495

50 Zakhary, D. R., Moravec, C. S., Stewart, R. W. & Bond, M. Protein kinase A (PKA)-dependent troponin-I phosphorylation and PKA regulatory subunits are decreased in human dilated cardiomyopathy. Circulation 99, 505–510 (1999).

51 van der Velden, J. et al. Increased Ca2+-sensitivity of the contractile apparatus in end-stage human heart failure results from altered phosphorylation of contractile proteins. Cardiovasc Res 57, 37–47 (2003). 10.1016/s0008-6363(02)00606-5

52 Jani, V. P. et al. Severe obesity in human HFpEF alters contractile protein function and organization. Science, eadz7118 (2026). 10.1126/science.adz7118

53 McCarty, D. M., Monahan, P. E. & Samulski, R. J. Self-complementary recombinant adeno-associated virus (scAAV) vectors promote efficient transduction independently of DNA synthesis. Gene Ther 8, 1248–1254 (2001). 10.1038/sj.gt.3301514

54 Tomek, J. et al. T-World: A highly general computational model of a human ventricular myocyte. bioRxiv (2025). 10.1101/2025.03.24.645031

55 Richter, W. et al. Conserved expression and functions of PDE4 in rodent and human heart. Basic Res Cardiol 106, 249–262 (2011). 10.1007/s00395-010-0138-8

56 Nikolaev, V. O., Bunemann, M., Hein, L., Hannawacker, A. & Lohse, M. J. Novel single chain cAMP sensors for receptor-induced signal propagation. J. Biol. Chem. 279, 37215–37218 (2004).

57 Darvish, M. et al. Heart failure: assessment of the global economic burden. Eur Heart J (2025). 10.1093/eurheartj/ehaf323

58 Authors/Task Force, M., et al. 2023 Focused Update of the 2021 ESC Guidelines for the diagnosis and treatment of acute and chronic heart failure: Developed by the task force for the diagnosis and treatment of acute and chronic heart failure of the European Society of Cardiology (ESC) With the special contribution of the Heart Failure Association (HFA) of the ESC. Eur J Heart Fail 26, 5–17 (2024). 10.1002/ejhf.3024

59 Redfield, M. M. & Borlaug, B. A. Heart Failure With Preserved Ejection Fraction: A Review. JAMA 329, 827–838 (2023). 10.1001/jama.2023.2020

60 Redfield, M. M., et al. Burden of systolic and diastolic ventricular dysfunction in the community: appreciating the scope of the heart failure epidemic. JAMA 289, 194–202 (2003). 10.1001/jama.289.2.194

61 Chahal, H. et al. Heart failure risk prediction in the Multi-Ethnic Study of Atherosclerosis. Heart 101, 58–64 (2015). 10.1136/heartjnl-2014-305697

62 Lam, C. S. P., Voors, A. A., de Boer, R. A., Solomon, S. D. & van Veldhuisen, D. J. Heart failure with preserved ejection fraction: from mechanisms to therapies. Eur Heart J 39, 2780–2792 (2018). 10.1093/eurheartj/ehy301

63 Messer, A. E., Jacques, A. M. & Marston, S. B. Troponin phosphorylation and regulatory function in human heart muscle: dephosphorylation of Ser23/24 on troponin I could account for the contractile defect in end-stage heart failure. J Mol Cell Cardiol 42, 247–259 (2007). 10.1016/j.yjmcc.2006.08.017

64 Feng, H. Z., Huang, X. & Jin, J. P. N-terminal truncated cardiac troponin I enhances Frank-Starling response by increasing myofilament sensitivity to resting tension. J Gen Physiol 155 (2023). 10.1085/jgp.202012821

65 Thompson, B. R., Martindale, J. & Metzger, J. M. Sarcomere neutralization in inherited cardiomyopathy: small-molecule proof-of-concept to correct hyper-Ca2+-sensitive myofilaments. American journal of physiology. Heart and circulatory physiology 311, H36–43 (2016). 10.1152/ajpheart.00981.2015

66 Joyce, W. et al. Genetic excision of the regulatory cardiac troponin I extension in high-heart rate mammal clades. Science 385, 1466–1471 (2024). 10.1126/science.adi8146

67 Dominic, K. L., Schmidt, A. V., Granzier, H., Campbell, K. S. & Stelzer, J. E. Mechanism-based myofilament manipulation to treat diastolic dysfunction in HFpEF. Front Physiol 15, 1512550 (2024). 10.3389/fphys.2024.1512550

68 Landim-Vieira, M. & Pinto, J. R. Can evolution-based studies inform modern medicine? Science 385, 1420–1421 (2024). 10.1126/science.ads2585

69 Sumandea, C. A. et al. Cardiac troponin T, a sarcomeric AKAP, tethers protein kinase A at the myofilaments. The Journal of biological chemistry 286, 530–541 (2011). 10.1074/jbc.M110.148684

70 Mendler, L., Braun, T. & Muller, S. The Ubiquitin-Like SUMO System and Heart Function: From Development to Disease. Circ Res 118, 132–144 (2016). 10.1161/CIRCRESAHA.115.307730

71 Fertig, B. et al. SUMOylation does not affect cardiac troponin I stability but alters indirectly the development of force in response to Ca(2). FEBS J 289, 6267–6285 (2022). 10.1111/febs.16537

72 Fentzke, R. C. et al. Impaired cardiomyocyte relaxation and diastolic function in transgenic mice expressing slow skeletal troponin I in the heart. J Physiol 517 (Pt 1), 143–157 (1999). 10.1111/j.1469-7793.1999.0143z.x

73 Hamo, C. E. et al. Heart failure with preserved ejection fraction. Nat Rev Dis Primers 10, 55 (2024). 10.1038/s41572-024-00540-y

## References

1 Tomek, J. et al. T-World: A highly general computational model of a human ventricular myocyte. bioRxiv (2025). 10.1101/2025.03.24.645031

2 Adeniran, I., MacIver, D. H., Hancox, J. C. & Zhang, H. Abnormal calcium homeostasis in heart failure with preserved ejection fraction is related to both reduced contractile function and incomplete relaxation: an electromechanically detailed biophysical modeling study. Front Physiol 6, 78 (2015). 10.3389/fphys.2015.00078

3 Kaplan, A. D., Boyman, L., Ward, C. W., Lederer, W. J. & Greiser, M. Ryanodine receptor stabilization therapy suppresses Ca(2+)- based arrhythmias in a novel model of metabolic HFpEF. J Mol Cell Cardiol 195, 68–72 (2024). 10.1016/j.yjmcc.2024.07.006

4 Rouhana, S. et al. Early calcium handling imbalance in pressure overload-induced heart failure with nearly normal left ventricular ejection fraction. Biochim Biophys Acta Mol Basis Dis 1865, 230–242 (2019). 10.1016/j.bbadis.2018.08.005

5 Selby, D. E., Palmer, B. M., LeWinter, M. M. & Meyer, M. Tachycardia-induced diastolic dysfunction and resting tone in myocardium from patients with a normal ejection fraction. J Am Coll Cardiol 58, 147–154 (2011). 10.1016/j.jacc.2010.10.069

6 Sequeira, V. et al. Perturbed length-dependent activation in human hypertrophic cardiomyopathy with missense sarcomeric gene mutations. Circ Res 112, 1491–1505 (2013). 10.1161/CIRCRESAHA.111.300436

